# Juvenile protein restriction compresses sexual dimorphism in mice

**DOI:** 10.1101/2025.07.09.663702

**Authors:** Amélie Joly, Lucas Rebiffé, Anne Lambert, Estelle Caillon, Sandrine Hughes, Benjamin Gillet, Mickael Zergane, Isabelle Rahioui, Pedro Da Silva, Chantal Rigaud, Ingrid Plotton, Justine Bruse, Hervé Guillou, Anne Fougerat, François Leulier, Filipe De Vadder

## Abstract

Sexual dimorphism is canonically understood as a fixed adult trait, scaled by gonadal hormones at puberty, but whether its magnitude is shaped earlier in life by juvenile nutrition during the post-weaning window has not been mechanistically investigated in mammals. We used dietary protein dilution in juvenile mice to study the plasticity of sexual dimorphism implementation. Sexual dimorphism failed to develop across somatic growth, energy metabolism, and hepatic molecular identity: as in undernourished children, males were more stunted than females and resembled the female baseline in all three levels. Androgens contributed to the male liver feminization, whereas the somato-metabolic compression persisted after orchidectomy. The reproductive axis diverged instead: males entered puberty without detectable delay while females deferred maturation, yet both sexes showed reduced reproductive output. Mammalian sexual dimorphism is therefore not a fixed adult property but a developmentally plastic trait, with the two sexes displaying distinct adaptation strategies along the somatic and reproductive axes.

## INTRODUCTION

Childhood undernutrition affects 149 million children under five worldwide, with stunting, defined as height-for-age more than two standard deviations below the median, as its most prevalent and consequential manifestation (*1*). Although stunting is multifactorial, dietary protein is a major nutritional determinant of linear growth (*2*), and stunted children show lower circulating essential amino acids, including the branched-chain amino acids (*3*). This burden is not distributed evenly between the sexes: across decades of demographic surveys, and confirmed by a recent systematic review and meta-analysis of more than 700,000 children, boys are consistently more stunted, wasted, and underweight than girls under nutritional stress (*4*, *5*). This disproportionate male vulnerability is present from early life, yet its proximate mechanisms remain unexplained, and no human study has resolved a specific developmental window in which it arises (*6*). In many species somatic growth is sexually dimorphic, and the magnitude of this dimorphism is itself diet-sensitive. Male traits collapse toward female values when juvenile resources are limiting, and the compression is asymmetric, the male response accounting for most of the cross-diet variability in dimorphism magnitude, consistent with the larger sex being the more developmentally plastic (*7–9*). In insects, sexual identity is specified cell-autonomously by the *doublesex*/*fruitless* cascade, rather than by a sex-determining hormone (*10*), and the condition dependence of exaggerated traits is traceable, in some species, to their heightened tissue-specific sensitivity to insulin/IGF signaling (*11*). In mammals, where sex-specific traits are instead scaled by gonadal and somatotropic signals, the underlying architecture of nutritional plasticity is unknown.

This plasticity should be most consequential during the mammalian post-weaning juvenile window. Indeed, this is a singular developmental period in which rapid somatic growth and the onset of puberty unfold together, when dimorphism is installed, and when nutrition can act on both at once. The somatotropic axis is a plausible substrate for this male vulnerability: its output is sexually dimorphic, installed during this same window, and directly tuned by nutrition. In the liver, where this program is best characterized, pulsatile growth hormone (GH) secretion from the male pituitary drives STAT5b in a phasic mode and opens hundreds of male-biased chromatin sites in phase with each pulse (*12*, *13*). The female axis, with more continuous GH exposure, is comparatively insensitive to perturbations of GH dynamics (*14*). This axis is itself nutritionally gated: hepatic insulin-like growth factor 1 (IGF-1) output and the feedback that shapes GH pulsatility depend on protein and energy status, so that dietary restriction lowers somatotropic drive and can shift GH secretion toward a more continuous, feminine pattern (*15*). Dietary protein restriction suppresses the somatotropic axis through a hepatic post-receptor defect in GH action (*16*), and we previously showed this for low-protein feeding from weaning (*17*, *18*), but whether sexual dimorphism still develops when this axis is nutritionally constrained has not been tested.

Here we profiled the development of sexual dimorphism in C57BL/6N mice fed a control diet (CD) or a matched-lipid protein-diluted variant (low-protein diet, LPD) from weaning to early adulthood, at three nested levels (somatic growth, energy metabolism, and hepatic molecular identity) together with reproductive maturation. Somatic, metabolic and hepatic sexual dimorphisms failed to develop under LPD, the males being compressed towards the female baseline. Puberty was differentially affected: females but not males showed a delay in reproductive maturation. Circulating testosterone nonetheless fell in LPD males. To dissect the contribution of gonadal androgens, we performed prepubertal orchidectomy concurrently with the dietary challenge, resolving the male response into androgen-dependent and androgen-independent programs. These findings establish, in a mammal, an asymmetric compression of sexual dimorphism, recasting its magnitude as a developmentally plastic rather than fixed adult property.

## RESULTS

### Sexual dimorphism in somatic growth fails to develop under juvenile protein restriction

To model sustained protein restriction during juvenile development, we fed ad libitum male and female C57BL/6N mice either a standard AIN-93G control diet (CD; 18.5% kcal protein, 17.5% kcal lipid, 64.0% kcal carbohydrate; 3.82 kcal/g) or a protein-diluted variant (LPD; 4.8% kcal protein, 18.0% kcal lipid, 77.2% kcal carbohydrate; 3.62 kcal/g) (**Table S1**). Throughout, we use “low-protein diet” (LPD) as shorthand for this targeted reduction of the protein-to-carbohydrate ratio at matched lipid. Under CD, somatic dimorphism developed progressively, males outgrowing females from post-natal day 35 (P35) in body length (**Fig. 1A**) and body weight (**Fig. S1A**). We calculated Cohen’s d as a measure of effect size, reflecting the biological difference between two groups, a higher Cohen’s d indicating a greater difference between groups. Under LPD, male and female growth trajectories converged and somatic sexual dimorphism failed to develop. Indeed, sex differences in body length and weight rose to d ∼ 4 by P42 under CD but stayed near d ∼ 0 throughout under LPD (**Figs. 1B, S1B**), the diet effect being much larger in males (d ∼ 5 from P35) than in females (d ∼ 2, declining to ∼1 by P56) (Figs. 1C, S1C). The same pattern was observed for femur length (**Fig. 1D-F**) and growth rate (**Fig. S1D**).

**Figure 1.**
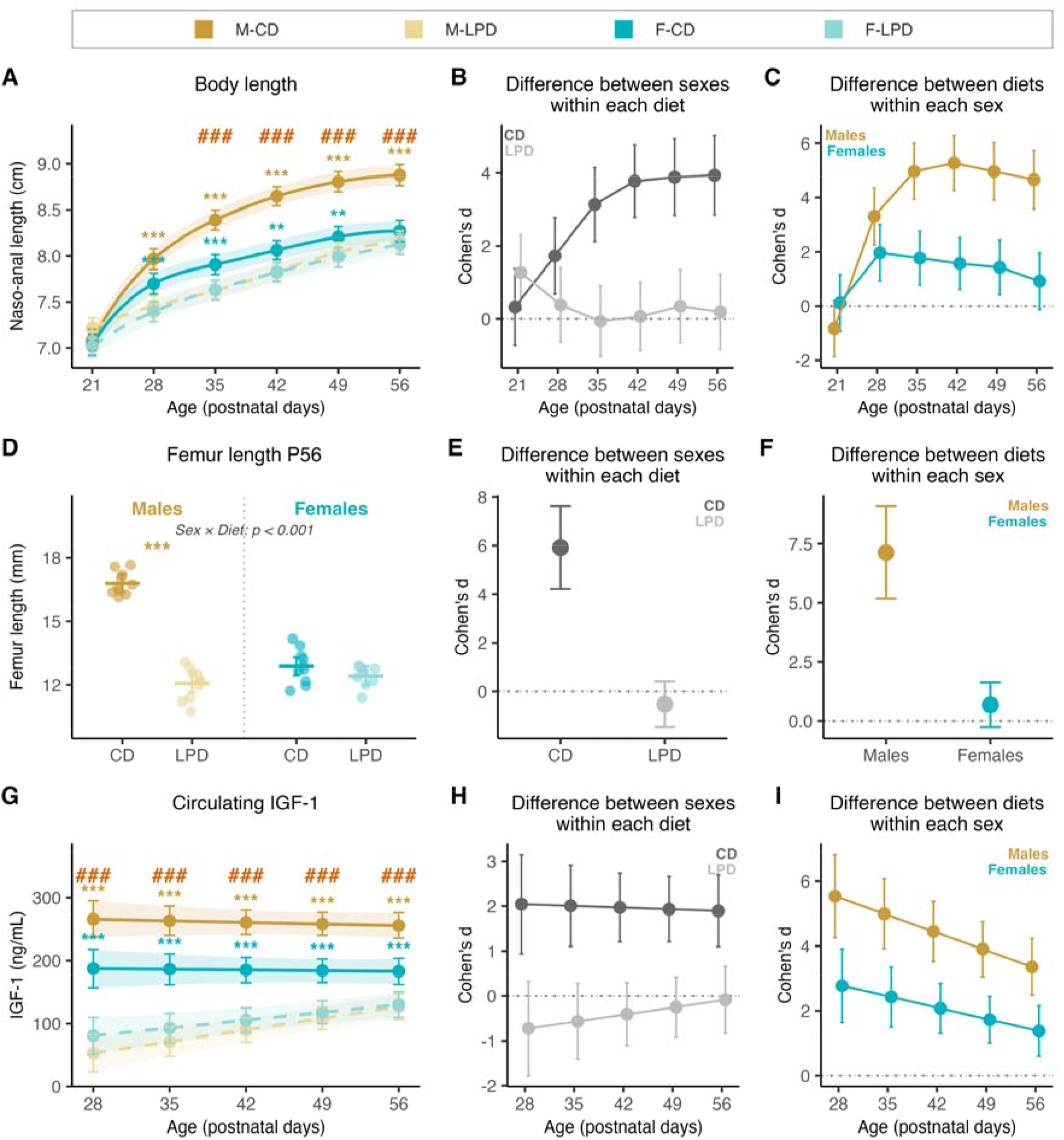
Sexual dimorphism in somatic growth fails to develop under juvenile protein restriction. C57BL/6N males and females were fed a control diet (CD) or a low-protein, matched-lipid diet (LPD) from weaning (post-natal day 21, P21) to P56. (**A**) Naso-anal length from P21 to P56 (n = 82 mice: 20 M-CD, 21 M-LPD, 20 F-CD, 21 F-LPD; 489 measurements), with (**B**) the sex difference within each diet and (**C**) the diet effect within each sex, measured as Cohen’s d with 95% CI. (**D**) Femur length at P56 (n = 39: 10 M-CD, 10 M-LPD, 10 F-CD, 9 F-LPD), with (**E**) and (**F**) as in (**B**) and (**C**). (**G**) Circulating IGF-1 from P28 to P56 (n = 132 mice, 196 samples), with (**H**) and (**I**) as in (**B**) and (**C**). Longitudinal traits (**A**, **G**) are estimated marginal means with 95% CI from linear mixed-effects models (see Methods); femur length (**D**) shows individual mice with the estimated marginal mean and 95% CI from a type III ANOVA on Sex × Diet. Colored asterisks, within-sex diet contrast (yellow, males; teal, females); red hash marks, Sex × Diet interaction at that age. *p < 0.05, **p < 0.01, ***p < 0.001; #p < 0.05, ##p < 0.01, ###p < 0.001. Full statistical output is in **Data S4**.

Organ weights showed no evidence of group-specific allometric scaling (**Fig. S1E**), consistent with a coherent loss of body-size dimorphism rather than tissue-specific effects.

Compared to control animals, IGF-1 fell more in males than in females under LPD (**Figs. 1G, 1I**), with the dimorphism observed in CD (d ∼ 2) collapsing under LPD (d ∼ 0; **Fig. 1H**), consistent with the somatic phenotype.

Food intake, scaled to lean mass, showed no evidence of a diet or sex difference (**Fig. S1F**). Whereas adult female mice mildly increase intake on low-protein diets, consistent with protein leverage (*19*), juveniles showed no evidence of such compensation in food intake (**Fig. S1G**), leaving a comparable ∼75% reduction in protein intake in both sexes. The differential growth response between males and females under LPD is therefore unlikely to reflect differential protein intake.

Across length, skeletal growth, and circulating IGF-1 (**Fig. 1**), the sex difference that builds under CD collapses under LPD. Somatic dimorphism is therefore a phenotype that fails to assemble under LPD, where it is compressed towards the female position.

### Sexual dimorphism in whole-body metabolism is compressed under LPD

We addressed next whether this convergence extends to dimorphic metabolic phenotypes. We profiled body composition by NMR (at 38 days) and whole-body metabolism by indirect calorimetry (from 39 to 41 days), when somatic dimorphism is fully established under CD (**Fig. 1**). Body composition was reshaped by LPD, affecting the relationship between body weight and both lean and fat mass (**Fig. 2A–B**). Sexual dimorphism in lean mass under CD, with males having more lean mass than females at comparable body weight, was buffered by LPD. On the contrary, adiposity, for which there was no evidence of dimorphism under CD, rose specifically in females under LPD. Whole-body energy expenditure, modeled at equal lean mass, rose under LPD in both sexes across both phases of the circadian cycle (**Figs. 2C, S2A–B**), consistent with the increased basal energetic cost per unit of tissue under reduced protein-to-carbohydrate ratios (*20*). The respiratory exchange ratio (RER) rose in parallel, indicating a shift toward carbohydrate utilization (**Figs. 2D, S2C**). This rise diverged between the sexes in the light phase (Sex × Diet p = 0.015; **Fig. 2D**), where LPD males lost the daytime drop into lipid oxidation that LPD females retained.

**Figure 2.**
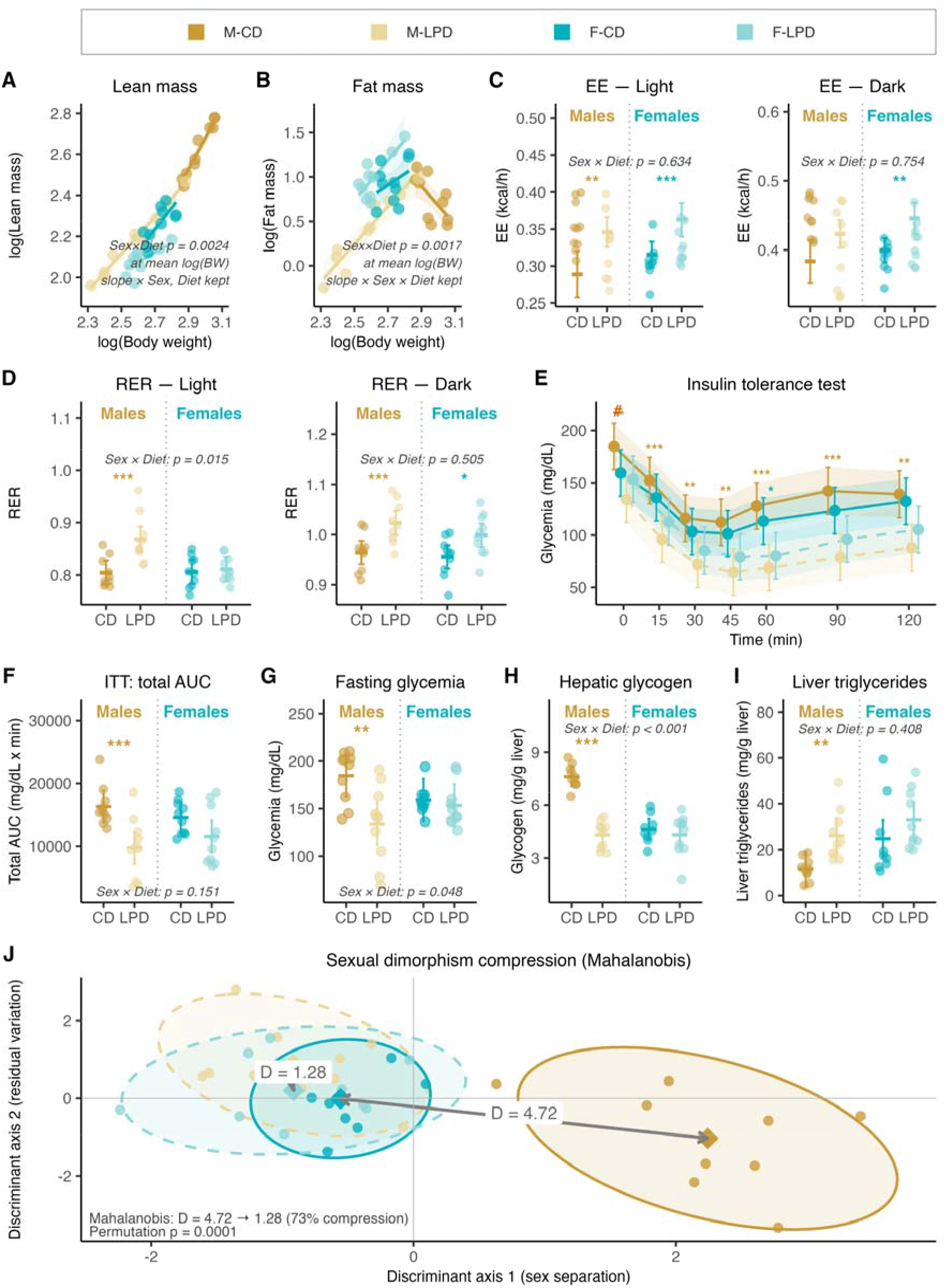
Juvenile protein restriction compresses whole-body metabolic sexual dimorphism. Mice fed CD or LPD from P21 underwent NMR body composition analysis (P38) and indirect calorimetry (P39 to P41). (**A, B**) Allometric scaling of lean mass (**A**) and fat mass (**B**) on body weight, log-log (n = 40, 10 per group). (**C**) Energy expenditure and (**D**) respiratory exchange ratio in the light and dark phases (n = 39: 10 M-CD, 9 M-LPD, 10 F-CD, 10 F-LPD; one LPD male excluded from the calorimetry recordings, cumulative food intake four times the next highest value). (**E**) Insulin tolerance test at P49 (0.5 U/kg i.p., n = 40) and (**F**) its total area under the curve; four mice reaching the humane endpoint were carried forward. (**G**) Fasting glycemia at P49 (n = 40). (**H**) Hepatic glycogen and (**I**) hepatic triglycerides at P56 (n = 39). (**J**) Multivariate compression of dimorphism: linear discriminant analysis of five somato-metabolic traits (body weight, fasting glycemia and insulin, hepatic glycogen and triglycerides) in 38 mice with complete data. Diamonds, centroids; ellipses, 95% regions; arrows, Mahalanobis distance between the sexes (D = 4.72 under CD, 1.28 under LPD; 73% compression, permutation p = 0.0001). Models: log-log ANCOVA with model selection on the slopes, retained model and Sex × Diet p-value at mean log body weight shown on the panel (**A, B**); lean-mass-adjusted mixed model with phase (**C, D**); mixed model on Sex × Diet × time (**E**); type III ANOVA on Sex × Diet (**F** to **I**). Colored asterisks, within-sex diet contrast (yellow, males; teal, females). *p < 0.05, **p < 0.01, ***p < 0.001. Full statistical output is in **Data S4**.

The overall RER increase pointing to a substrate-level signal, we next probed glucose homeostasis, which is known to be sexually dimorphic (*21*). Under CD, males tended to fast at higher glycemia than females but we found no evidence of a sex difference in fasting glycemia or insulin tolerance (**Fig. 2E–G**). Under LPD this ordering reversed (Sex × Diet p = 0.048; **Fig. 2G**): LPD males fasted below LPD females and showed the larger glycemic response to insulin (**Fig. 2E–F**). Similar to linear growth, the diet effect was far larger in males than in females. The response to an oral glucose tolerance test and corresponding insulin levels followed the same pattern, again driven by a strong male response to LPD, with no evidence of a diet effect in females (**Fig. S2D–E**). We then turned to the liver, a key integrator of somatotropic and metabolic signals. Once again, the established mammalian dimorphism in hepatic glycogen storage (*22*), apparent under CD, collapsed to the female baseline under LPD (Sex × Diet p < 0.001; **Fig. 2H**), consistent with the carbohydrate-shifted RER. Hepatic triglyceride content and lipid-droplet geometry, by contrast, responded concordantly in both sexes with increased hepatic lipid storage under LPD (**Figs. 2I, S2F**). To quantify all these signals jointly, we performed discriminant analysis on five somato-metabolic traits selected under CD for dimorphism and non-redundancy (see methods for details). The Mahalanobis distance (D) between sex centroids, which measures how far apart sexes are given the spread within each group, fell from D = 4.72 under CD to D = 1.28 under LPD, indicating a 73% compression of the somato-metabolic sexual dimorphism (**Figs. 2J, S2G**). This quantification confirms that the development of a strong growth and metabolic sexual dimorphism is largely prevented by low-protein feeding during the juvenile period.

### Hepatic sexual dimorphism contracts asymmetrically under juvenile protein restriction

Hepatic sexual dimorphism is itself a readout of the sex-biased somatotropic axis (*23*). We thus profiled, in both sexes, the nutritional signal reaching the liver and the hepatic response it elicits. Despite a fourfold dietary protein dilution, the fall in portal amino acid concentrations was partial and selective (**Fig. S3A**): branched-chain amino acids (valine, leucine, isoleucine) and phenylalanine dropped by 29–46% under LPD in both sexes, while several non-essential amino acids (notably glutamine and arginine) were maintained or rose, consistent with partial upstream metabolic compensation, likely catabolic. Plotting the LPD-induced log_2_ fold-change in males against females (**Fig. 3A**) reveals that this response was concordant between the sexes (Pearson correlation coefficient r = 0.81, portal compartment), with no evidence of a Sex × Diet interaction for any of the 19 amino acids across the four vein and nutritional-state combinations (**Fig. S3A**). The portal blood entering each liver therefore carries the same low-BCAA/phenylalanine signal under LPD, with no detectable sex difference in its amplitude or composition.

**Figure 3.**
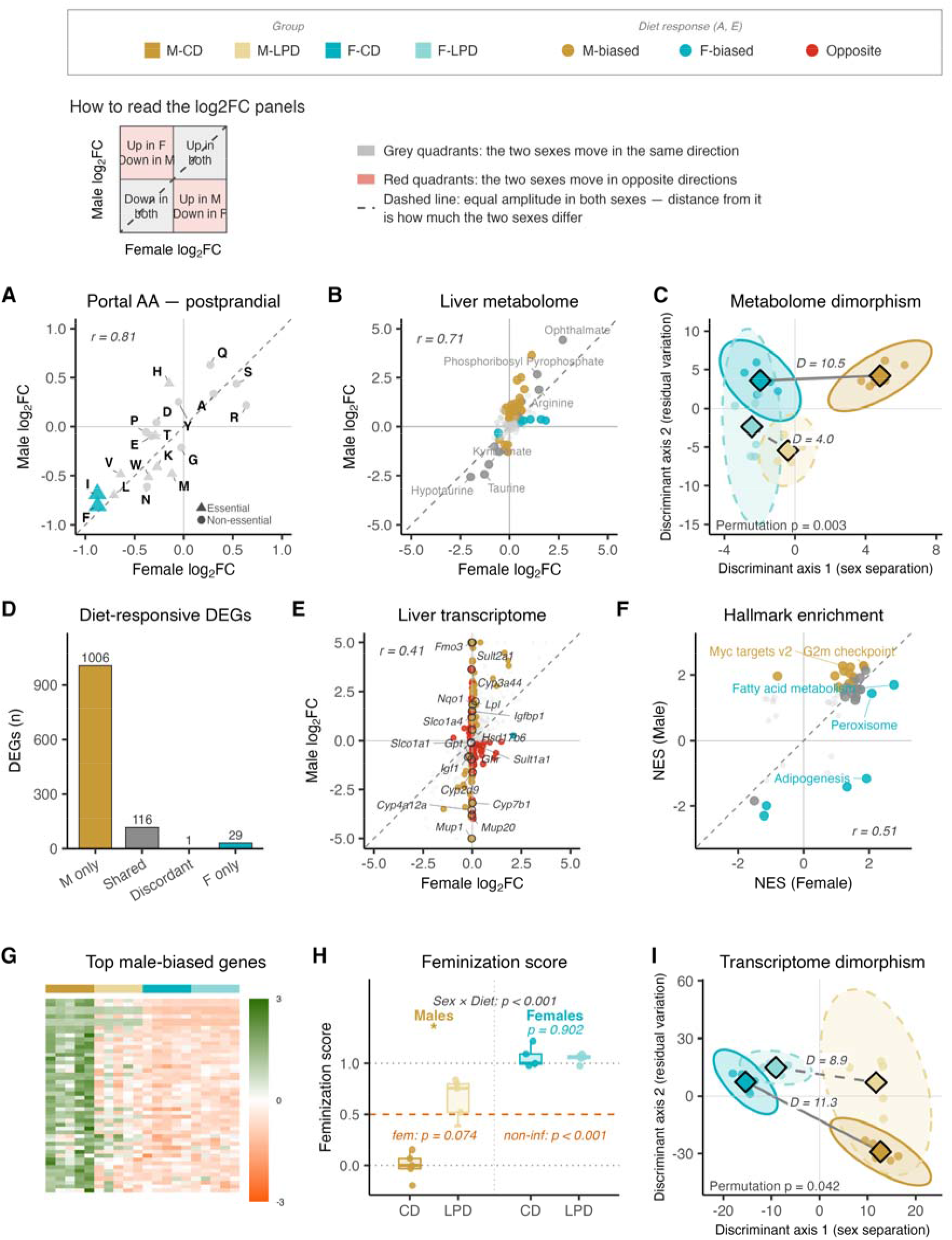
Hepatic sexual dimorphism compresses asymmetrically under juvenile protein restriction. Portal serum, hepatic polar metabolome and liver transcriptome were profiled at P56. Teal, female-associated diet response; yellow, male-associated; grey, shared or nonsignificant; red, divergent between sexes (**A, E**). The key explains the log_2_ fold-change panels. (**A**) Per-amino-acid log2 fold change (LPD versus CD) in postprandial portal serum, females (x) versus males (y) (n = 28, 7 per group). (**B**) Per-metabolite log2 fold change in liver (n = 24, 6 per group; 165 metabolites). (**C**) Metabolome dimorphism by linear discriminant analysis; diamonds, centroids; ellipses, 95% regions; D, Mahalanobis distance between the sexes (10.5 under CD, 4.0 under LPD; permutation p = 0.003). (**D**) Diet-responsive genes by category, (**E**) per-gene shrunken log_2_ fold change and (**F**) Hallmark gene-set enrichment scores, females versus males (RNA-seq, n = 5 per group). (**G**) Top 50 male-biased genes under CD; rows, genes; columns, mice; color, per-gene z score. (**H**) Feminization score: projection onto the M-CD to F-CD discriminant axis (321 strictly sex-biased genes), LPD projected out of sample, CD by leave-one-out, rescaled to median(M-CD) = 0, median(F-CD) = 1. (**I**) Transcriptome dimorphism as in (**C**), on the 1,255 sex-biased genes (D = 11.3 versus 8.9, P = 0.042). Correlations are Pearson’s r; amino acids and metabolites by `limma`, genes by `DESeq2` (Benjamini-Hochberg p < 0.05, |log_2_ fold change| > 1); (**H**) by aligned rank transform ANOVA, with one-sided t tests against the 0.5 midpoint in orange. Full statistical output is in **Data S4**.

To compare how the liver of each sex responds to this common signal, we profiled 165 polar metabolites of central-carbon and amino-acid metabolism. Plotting the LPD-induced log_2_ fold-changes in males against females (**Fig. 3B**) showed a concordant response (no evidence of a Sex × Diet interaction for any of the 165 metabolites), with male amplitudes systematically larger. Male and female livers thus mount the same biochemical adaptation to LPD, spanning the urea cycle, glutathione/taurine and one-carbon metabolism, and the pentose phosphate pathway. These changes differ in intensity rather than in nature, though from sex-distinct baselines, since the hepatic metabolome is itself dimorphic under CD. The Mahalanobis distance between sex centroids (as in **Fig. 2J**) fell from 10.5 to 4.0 (a 62% compression), driven mainly by displacement of the male centroid toward the female position (**Fig. 3C**).

To examine the transcriptional response underlying this remodeling, we profiled the hepatic transcriptome by bulk RNA-seq. Principal component analysis separated samples primarily by sex, with male LPD livers displaced toward the female position (**Fig. S3B**). Differential expression was strongly asymmetric: 1,123 genes responded to LPD in males versus 146 in females (**Figs. 3D, S3C–D**). Among those, 117 genes were responsive in both sexes, with 116 displaying a concordant response, and 238 genes showed a Sex × Diet interaction (**Fig. 3D**; **Data S2**). However, most of these genes responded strongly in males with negligible amplitude in females (**Fig. 3E**). Gene-set enrichment revealed that proliferative and acute-stress programs were engaged more strongly in males while lipid-metabolism programs were more activated in females (**Figs. 3F, S3E–F**). Hepatic transcriptional plasticity to LPD is therefore sexually dimorphic in magnitude, not in direction. This comparatively muted female response to LPD is consistent with prior reports in adult C57BL/6J liver (*24*).

Superimposed on this, the sex-biased hepatic transcriptional identity itself was asymmetrically eroded. Canonical male-biased targets of the pulsatile GH/STAT5b axis (e.g. *Cyp4a12a*, *Cyp2d9*, *Mup1/20*) were strongly down-regulated in LPD males, while female-biased genes (e.g. *Fmo3*, *Sult2a*) were reciprocally induced in LPD males toward their female level (**Figs. 3E, 3G, S3D**; **Data S2**). The male liver thus shifts toward the female transcriptional state with repression of male-biased genes and induction of female-biased ones, mirroring the transcriptional feminization produced when the male-determining repressor BCL6 is removed from the liver (*25*). To quantify this, we calculated a transcriptional feminization score for each liver (where M-CD = 0 and F-CD = 1). LPD males shifted toward the female pole, even if feminization was incomplete. On the contrary, LPD females stayed at the female pole (**Fig. 3H**). Accordingly, the Mahalanobis distance over the 1,255 sex-biased genes contracted by 21% in LPD compared to CD (D = 11.3 to 8.9; **Fig. 3I**). The compression is therefore not a symmetric meeting in the middle but a specific displacement of the male hepatic transcriptome toward the female state.

We then asked whether this hepatic transcriptional feminization is developmentally restricted by re-analyzing, through our pipeline, published transcriptomic data from adult C57BL/6J mice fed a low-protein diet (*24*), with no new adult cohort (**Fig. S3G–J**). Even if the designs differed (Green: 7% whey-based vs 21% protein; Joly: 4.8% casein vs AIN-93G), allowing a qualitative comparison only, the two components dissociated by age. Indeed, in males, the nutrient-stress response was attenuated only 1.7-to-9.7-fold in adults relative to juvenile amplitude, whereas the GH/STAT5b program was attenuated 56-to-147-fold. Although the overall response to low protein was much stronger in juveniles than in adults (**Fig. S3I**), the nutrient-stress response, driven by the stress-activated transcription factor ATF4 together with PPARα (*26*), was detectable and more induced in males than in females in both datasets (**Fig. S3G-H**).The dimorphic GH/STAT5b program, by contrast, was largely lost in adults. Strikingly, we found no evidence of male hepatic transcriptional feminization in adults (**Figs. 3H, S3J**). The nutrient-stress response to protein restriction is thus preserved across ages, whereas the sex-dimorphic GH/STAT5b remodeling and subsequent hepatic transcriptional feminization is confined to the juvenile window.

Hepatic FGF21 has been proposed as a mechanism of undernutrition-related GH resistance and consequent growth defect (*27*). In line with this idea, LPD strongly induced hepatic *Fgf21* in both sexes (**Fig. S3K**) and raised circulating FGF21 (**Fig. S4A**). However, we found no evidence that hepatocyte-specific *Fgf21* deletion (*28*), largely abolishing the LPD-induced rise in circulating FGF21 levels, rescued any LPD-associated male phenotype (**Fig. S4**), which does not support its role in GH resistance in males. In females, however, hepatic *Fgf21* deletion partially restored somatic growth by P56 (**Fig. S4B–F**). The same hepatokine thus appears to have a sex-restricted function: it constrains female somatic growth under LPD with no detectable contribution to male stunting.

Therefore, juvenile protein restriction compresses sexual dimorphism by displacing the male phenotypes toward the female’s, the compression scaling with biological integration (73% somato-metabolic, 62% metabolome, 21% transcriptome). Dimorphism behaves as an integrated, multivariate trait, most convergent in composite physiological readouts and only partially traceable to the transcriptome.

### Puberty timing in males appears unaffected, but androgen-dependent outputs are reduced

The male hepatic transcriptomic program is governed by synergistic effects of both gonadal androgens and the GH/STAT5b axis to install and maintain male hepatic transcriptomic identity. Because that androgen tone rises at male puberty, an event unfolding within the same post-weaning window we perturb, we turned next to male reproductive maturation. We first assessed balanopreputial separation (BPS), a non-invasive marker of pubertal initiation (*29*). There was no evidence that the age at BPS differed between diets (**Fig. 4A**)or that testis mass differed between diets at equivalent body weight (**Fig. 4B**), hence no sign of impaired testis development. There was weak evidence that LPD reduced sperm count and seminal vesicle weight, two markers of androgenic activity *in vivo* (*30*), each by about a third (**Fig. 4C–D**). Protein restriction also sharply reduced hepatic expression of the *Mup* cluster (**Fig. 4E**), which encodes the liver-derived major urinary proteins that mediate pheromone-based chemical signaling, predominant in males (*31*). Accordingly, urinary MUP protein was markedly reduced (**Fig. 4F**). LPD males thus enter puberty without detectable delay, but with reduced androgen-dependent outputs.

**Figure 4.**
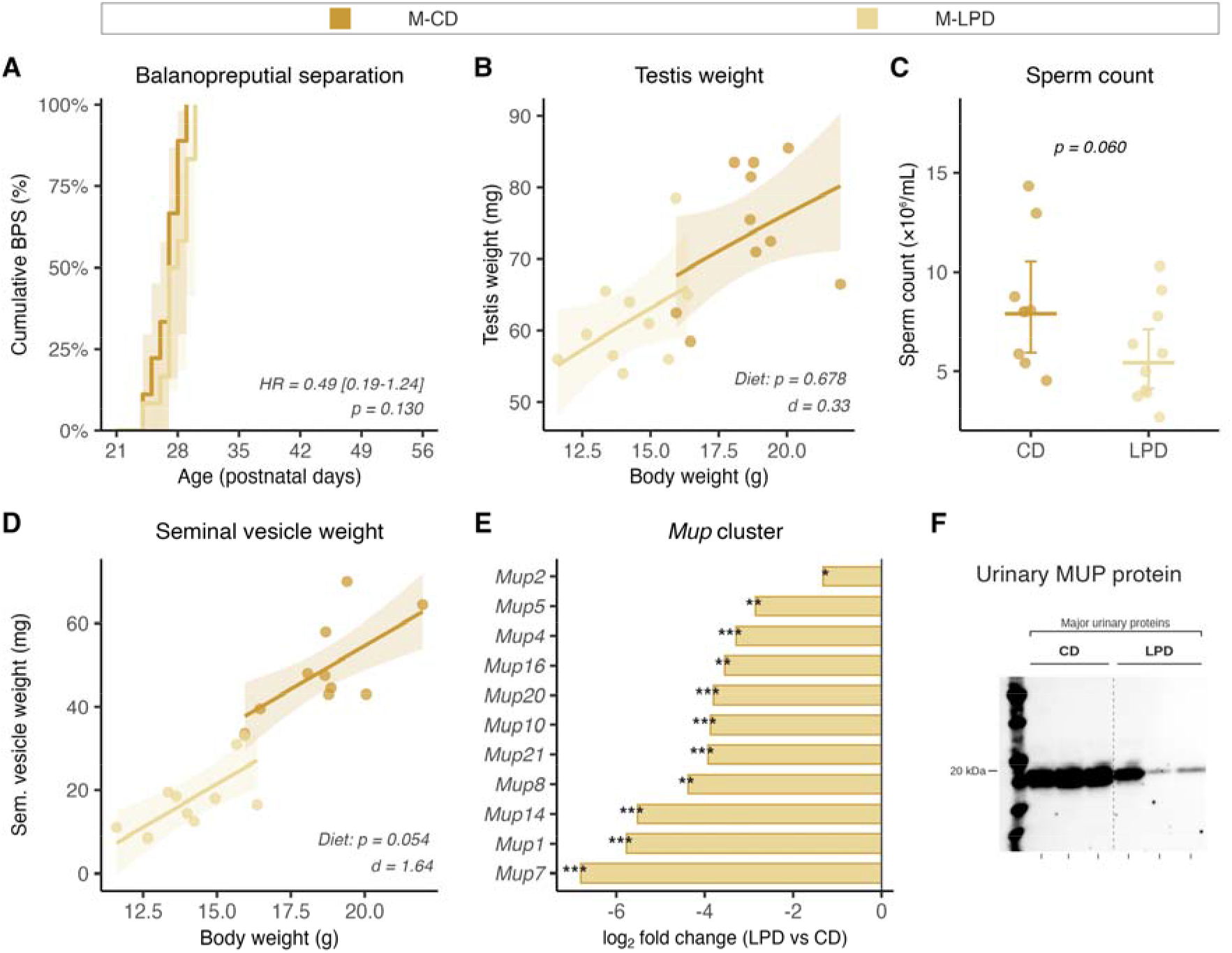
Male puberty advances on time, but the hepatic pheromone output is silenced. Males were fed CD or LPD from P21. (**A**) Cumulative incidence of balanopreputial separation from P21 to P56 (n = 21: 9 CD, 12 LPD); shaded areas, 95% CI of the Kaplan-Meier estimates; HR, hazard ratio from a Cox proportional hazards model with 95% CI; p-value, log-rank test. (**B**) Testis weight at P56 against body weight (n = 20, 10 per diet). (**C**) Epididymal sperm count at P56 (n = 19: 9 CD, 10 LPD). (**D**) Seminal vesicle weight at P56 against body weight (n = 20, 10 per diet). (**E**) Hepatic expression of the Mup cluster, shrunken log_2_ fold change (LPD versus CD) in the male liver RNA-seq of Fig. 3 (n = 5 per diet). (**F**) Urinary major urinary proteins at P56 (Western blot, three males per diet, equal urine volume per lane). Panel (**F**) is qualitative: the CD bands are saturated at the exposure used, so the quantitative statement rests on (**E**); the uncropped membrane is provided as **Fig. S8**. Organ weights (**B, D**), ANCOVA with body weight as covariate on the Box-Cox scale; sperm count (**C**), type III ANOVA on the log scale; diet p-value and Cohen’s d on each panel. Points are individual mice, with the estimated marginal mean and 95% CI in (**C**). Asterisks in (**E**), DESeq2-adjusted p-value (*p < 0.05, **p < 0.01, ***p < 0.001). Full statistical output is in **Data S4**.

### Prepubertal orchidectomy separates androgen-dependent and androgen-independent components of hepatic feminization

The gonadal and somatotropic (GH/IGF-1) axes are tightly interconnected, and sex steroids shape the male GH secretory pattern that drives growth (*32*). To understand how potentially impaired gonadal function contributes to the somatic growth and metabolic phenotype observed in LPD-fed males, we combined both dietary arms (CD and LPD) with prepubertal orchidectomy (SHAM and ORX) performed at weaning. Orchidectomy (ORX) drove circulating testosterone below the limit of quantification in 13 of 18 castrated males and in none of the 19 sham males, a fall of at least 88% relative to the median of control-fed sham males. Among castrated males, residual testosterone remained quantifiable in 5 of 10 fed CD but in none of the 8 fed LPD (p = 0.036). LPD alone lowered it by 38% in sham males (**Fig. 5A**), consistent with the partial loss of androgen-associated traits described in **Fig. 4**. We found no evidence that ORX modified the somatic response to LPD: for body weight, body length, organ weights and femur length, no Diet × Surgery interaction was detected (**Figs. 5B–E, S5A–E**). In line with this, the drop in circulating IGF-1 induced by LPD occurred independently of ORX surgery (**Fig. 5F**). Glucose tolerance and fasting glycemia showed no evidence of an ORX effect (**Fig. S5F–G**). The somatic and metabolic feminization of the male upon LPD thus persisted without the testes.

**Figure 5.**
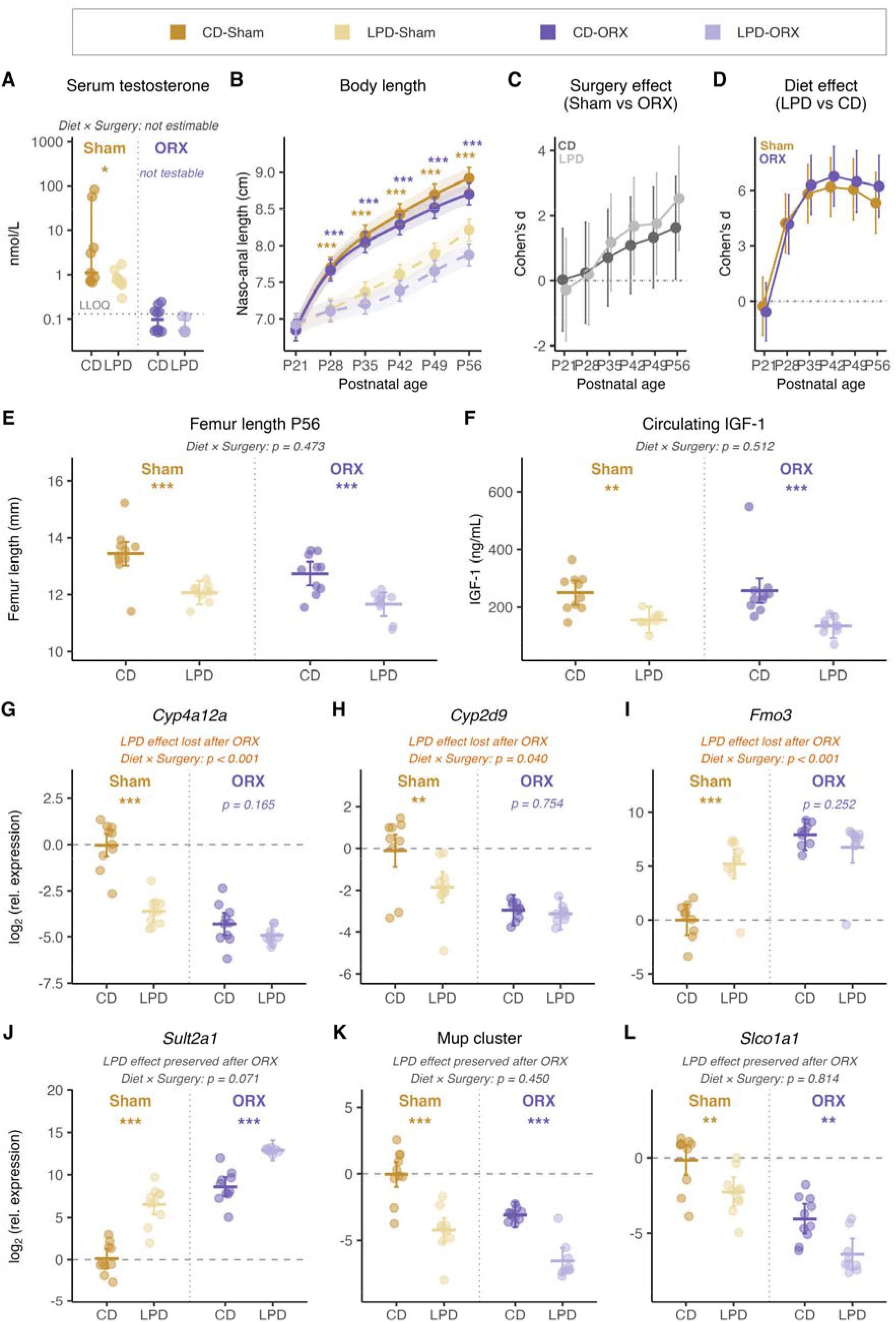
Prepubertal orchidectomy separates two parallel mechanisms remodeling the male liver under juvenile protein restriction. Males underwent bilateral orchidectomy (ORX) or sham surgery at P21 and were fed CD or LPD until P56 (n = 40, 10 per Diet × Surgery group). (**A**) Serum testosterone at P56 (n = 37: 9 CD-Sham, 10 LPD-Sham, 10 CD-ORX, 8 LPD-ORX), log scale; bars are medians with bootstrap 95% CI. The dotted line is the limit of quantification (0.13 nmol/L); values below it are censored. Diet effect, exact Wilcoxon rank-sum in Sham males; all LPD-ORX values lie below the limit, so no contrast is estimable in that stratum. (**B**) Naso-anal length from P21 to P56 (240 measurements), with (**C**) the surgery effect within each diet and (**D**) the diet effect within each surgery, measured as Cohen’s d with 95% CI. (**E**) Femur length (n = 40) and (**F**) circulating IGF-1 (n = 39) at P56. (**G** to **I**) Hepatic transcripts whose diet response is lost after ORX and (**J** to **L**) transcripts whose diet response is preserved, by RT-qPCR (n = 39: 10 CD-Sham, 10 LPD-Sham, 10 CD-ORX, 9 LPD-ORX), as log_2_ relative expression. Body length (**B**) is estimated marginal means with 95% CI from a linear mixed model (see Methods); panels (**E**) to (**L**) show individual mice with the estimated marginal mean and 95% CI from a type III ANOVA on Diet × Surgery. Colored asterisks, within-surgery diet contrast (yellow, Sham; purple, ORX; *p < 0.05, **p < 0.01, ***p < 0.001); the Diet × Surgery interaction p-value is given on panels (**E**) to (**L**). Full statistical output is in **Data S4**.

We then asked whether the male-specific hepatic transcriptional response to LPD established in **Fig. 3** is mediated by gonadal androgens. First, we assessed expression of non-dimorphic genes related to the somatotropic axis, and saw that, as expected, both *Ghr* and *Igf1* responded to LPD, with no evidence of an ORX effect (**Fig. S5H**). Because the LPD male liver is transcriptionally feminized (**Fig. 3**), we then profiled a panel of sexually dimorphic hepatic transcripts selected based on the Rampersaud classification (*13*) and assessed if their altered expression in LPD was androgen-dependent. We found two main profiles with both androgen-dependent genes, characterized by an abolished difference between CD and LPD upon ORX (**Figs. 5G–I, S5H**), and androgen-independent genes, with a maintained dietary effect independently of ORX (**Figs. 5J–L, S5H**).

The compression of hepatic transcriptomic dimorphism is thus only partly androgen-driven: the reduced androgenic tone observed in adult LPD males (**Figs. 4C–F, 5A**) contributes to the transcriptomic feminization of the liver but cannot alone account for the suppression of the somatotropic axis, nor for the metabolic and hepatic phenotypes that persist after castration.

### Juvenile protein restriction defers female reproductive maturation

In females, LPD had smaller somatic and hepatic effects. Because gestation and lactation place the bulk of the metabolic cost of reproduction on the female (*33*, *34*), constraints sensed during the post-weaning window may instead be channeled toward reproductive timing.

We therefore examined female pubertal maturation by monitoring vaginal opening (VO) daily from P21. LPD delayed VO by 12.5 days (**Fig. 6A**). It is well known that puberty initiation in females requires reaching a body-size and adiposity threshold rather than a given age (*35*). Consistently, we found no evidence of a difference in body weight at VO (**Fig. 6B**). Causal mediation analysis, aiming to discriminate between direct effect of LPD on VO and the indirect effect of LPD through body weight changes, indicated that the effect of LPD on VO was not mediated by body weight (**Fig. S6A**). Circulating leptin is the canonical adipostatic signal licensing female puberty in rodents (*36*), but at P28, i.e., preceding the median CD VO date, there was no evidence of hypoleptinemia in LPD females (**Fig. S6B**). This is consistent with peripubertal female mice, in which a 30% caloric restriction delays reproductive maturation while lowering circulating leptin only modestly, the frequency of kisspeptin pulses tracking food availability largely independently of leptin (*37*). LPD females thus reach the same body-size gate as CD females, but considerably later. Their juvenile growth rate is reduced (**Fig. S1D**), and age at VO varied inversely with that rate (**Fig. 6C**), all LPD females falling below ∼0.30 g/day, the rate above which most CD females reached VO (**Fig. 6C**, **Fig. S6C–D**).

**Figure 6.**
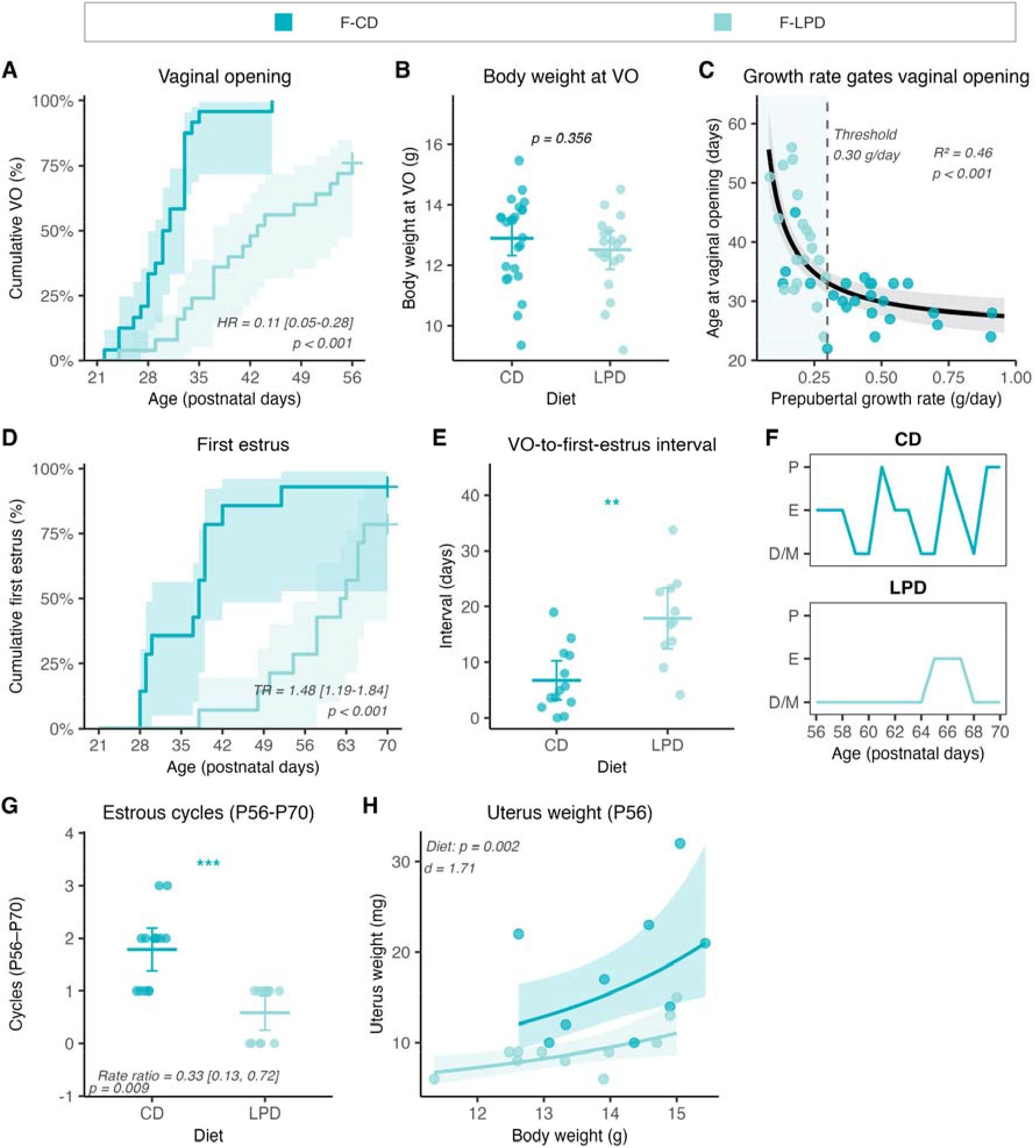
Juvenile protein restriction defers female reproductive maturation. Females were fed CD or LPD from P21 and monitored for vaginal opening (VO), estrous cyclicity and reproductive organs. (**A**) Cumulative incidence of VO from P21 to P56 (n = 49: 24 CD, 25 LPD, 6 censored); shaded areas, 95% CI of the Kaplan-Meier estimates; HR, hazard ratio (Cox model) with 95% CI. (**B**) Body weight at VO (n = 43: 24 CD, 19 LPD; cohort-adjusted contrast). (**C**) Age at VO against prepubertal growth rate (P21 to P28) in the same females (reciprocal model, nonlinear least squares); dashed line, growth-rate threshold from segmented regression (0.30 g/day); shaded band, bootstrap 95% prediction interval (5,000 iterations). (**D**) Cumulative incidence of first estrus from P21 to P70 (n = 28: 14 CD, 14 LPD, 4 censored); TR, time ratio from a Weibull accelerated failure time model. (**E**) Interval from VO to first estrus (n = 24: 13 CD, 11 LPD; Wilcoxon rank-sum test). (**F**) Representative cycles from P56 to P70 in one CD (top) and one LPD female (bottom). P, proestrus; E, estrus; D/M, diestrus or metestrus. (**G**) Complete cycles between P56 and P70 (n = 26: 14 CD, 12 LPD; Poisson GLM, rate ratio). (**H**) Uterus weight at P56 against body weight (n = 20: 9 CD, 11 LPD; ANCOVA on the Box-Cox scale). Points are individual mice with the estimated marginal mean and 95% CI. **p < 0.01, ***p < 0.001. Full statistical output is in **Data S4**.

Reaching VO is only the first of two events that mark female pubertal maturation: it must be followed by the first ovulatory cycle, signaled by the first cornified vaginal smear (first estrus) (*38*). LPD delayed first estrus by 24.5 days (**Fig. 6D**). The interval between VO and first estrus was itself extended, from a median of 5 to 17 days (**Fig. 6E**), suggesting that the delay in first estrus upon LPD is not only attributable to a delay in attaining the weight threshold for VO.

We next tested whether LPD females eventually reach a full adult reproductive state. Across the recording window (P56–P70), CD females completed a median of two estrous cycles, whereas LPD females completed one and frequently held in diestrus or metestrus (**Fig. 6F–G**, **Fig. S6E**). In addition, at equivalent body weight, the uterus of LPD females weighed 38% less than that of CD females (**Fig. 6H**). Because uterine mass tracks circulating estrogen (*39*), this is consistent with the idea that the reproductive axis has not established the cyclic ovarian steroid output that drives adult reproductive function at P56. Thus, it is likely that female pubertal maturation under LPD is delayed through both a growth-gated approach to VO and an incomplete activation of the gonadotropic axis, resulting in abnormal ovulatory cycling.

Because LPD induces a marked rise in hepatic FGF21 (**Fig. S3**), and chronic FGF21 overexpression can disrupt female reproduction (*40*), we asked whether hepatic FGF21 mediates the LPD-induced female reproductive delay. We found no evidence that hepatocyte-specific *Fgf21* deletion shifted VO timing, but it accelerated first estrus under CD; under LPD only mutant females reached first estrus by P56 (**Fig. S6F-G**). The diet effect on uterine weight seen in control females was not detected in mutants, consistent with this earlier cycling onset (**Fig. S6H–I**). Thus, our results suggest that hepatic FGF21 acts as a constitutive tonic brake on the post-VO transition to ovulatory cycling, suggesting that elevated FGF21 levels in LPD could contribute to the impaired female reproductive maturation.

LPD therefore differentially reshapes the somato-metabolic and reproductive axes in males and females during the juvenile period.

### Two contrasting strategies of developmental plasticity lead to a similarly decreased reproductive output

Having established that LPD differentially impacts sexual maturation in males and females, we next tested how LPD affected reproductive output. To evaluate whether the decrease in circulating testosterone (**Fig 5A**) despite normal puberty timing (**Fig. 4A**) compromised basal reproductive function in LPD-fed males, we paired adult males with hyperfertile OF1 females over three non-competitive mating cycles (**Fig 7A–C**). LPD reduced per-cycle conception in males (47% in LPD vs 77% in CD; **Fig. 7A**). Among the fertilized females, there was no evidence of a diet effect on total births or pup weight (**Figs. 7B, S7A–C**). Expected offspring output per mating cycle fell ∼41% in LPD (**Fig. 7C**), this difference arising mostly from the conception stage (**Fig. S7D**). LPD males thus carry a cost expressed almost entirely as reduced per-cycle conception, with litter quality preserved.

**Figure 7.**
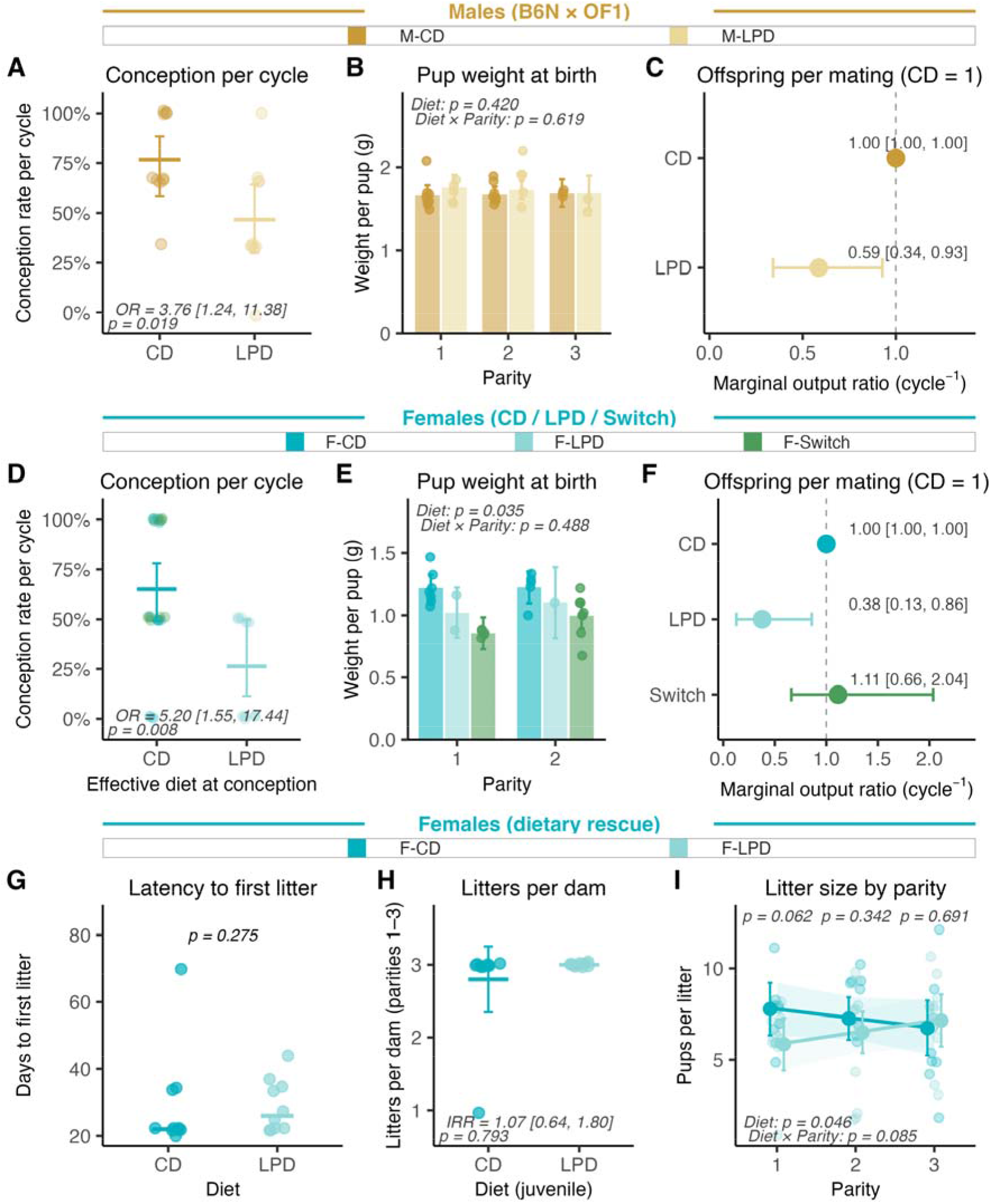
Reproductive output reveals two contrasting strategies of developmental plasticity. Three independent cohorts were assessed: B6N males paired with hyperfertile OF1 females in single-male, noncompetitive cycles (**A** to **C**; n = 20 males, 10 per diet, 60 mating cycles); B6N females in a three-arm design, CD throughout, LPD throughout, or Switch, that is CD before conception and between cycles with LPD restricted to gestation, paired with confirmed-fertile males (**D** to **F**; n = 30 females, 10 per arm, 59 resolved cycles); and B6N females raised on CD or LPD from P21 to P70, then switched to CD and paired with proven studs for three months (**G** to **I**; n = 20 dams, 10 per diet, 58 litters at parities 1 to 3). (**A, D**) Conception per cycle: dots, per-animal rate; bar, estimated marginal mean with 95% CI from a logistic GLM; OR, odds ratio with 95% CI. In (**D**), cycles are classified by the effective diet at fertilization, CD and Switch being pooled at conception. (**B, E**) Weight per pup at birth by parity (linear model on Diet × Parity). (**C, F**) Marginal reproductive output per cycle, as the bootstrap median ratio to the CD reference with 95% percentile CI (5,000 iterations). (**G**) Latency to first litter (Wilcoxon rank-sum test). (**H**) Litters per dam (Poisson GLM; IRR, incidence rate ratio). (**I**) Litter size by parity (linear mixed model with dam as random intercept). Full statistical output is in **Data S4**.

To understand how LPD during the juvenile period affected female reproductive output, we followed three arms across 2 parities: CD throughout, LPD throughout, and a Switch arm with CD until conception then LPD for gestation, to account for the effect of LPD during pregnancy itself (**Fig 7D–F**). To assess the effect of diet on conception rate, we pooled females on the CD throughout and Switch arms, as they were both fed CD during the periconception window. Conception per cycle was on average 65% under CD versus 26% under LPD (**Fig. 7D**). Among the females who conceived, there was no evidence that litter size or perinatal mortality differed across arms (**Fig. S7E–F**). However, pup weight at birth differed with gestational diet (**Fig. 7E**) being lowest in Switch litters, while the offspring of LPD mothers had an intermediate weight (1.05 g). Since conception itself was rare in LPD females (26%), we interpret this finding with caution. Expected offspring output per mating cycle collapsed to 1.4 pups/cycle under continuous LPD against 3.75 in CD (63% reduction), while Switch matched CD (4.15 pups/cycle; **Fig. 7F**). Similar to the male deficit, the female LPD gap was dominated by reduced conception rate rather than litter size (**Fig. S7G**). Periconceptional protein restriction thus governs whether a litter is conceived while gestational protein restriction seems to regulate how big pups are at birth.

To test whether the LPD-associated deficit of female reproduction was reversible, we raised weanling B6N females on CD or LPD until adulthood, then switched the LPD group to CD, and paired all females with proven studs for three months (**Fig 7G–I**). We found no evidence that latency to first litter (**Fig. 7G**), the number of litters (**Fig. 7H**) or per-pup birth weight (**Fig. S7H**) differed, nor that the impairment of reproductive capacity (**Fig. 7D–F**) persisted once the nutritional constraint was lifted. One transient imprint remained: litter size showed an overall diet effect, even if rescued dams rapidly converged with CD (**Fig. 7I**). Juvenile protein restriction thus limits female reproductive capacity without permanently preventing it.

While males and females facing juvenile protein restriction adopt very different adaptive strategies in terms of somato-metabolic and reproductive development, culminating in a strong compression of sexually dimorphic traits, their reproductive output is thus similarly decreased, mainly through a diminished conception rate.

## DISCUSSION

Our data show that juvenile protein restriction across the P21–P56 window compresses sexual dimorphism in the mouse at three nested levels (somatic growth, energy metabolism, and hepatic molecular identity). The two sexes are reshaped along largely separate axes: the male is displaced toward the female baseline along the somatic, metabolic and hepatic-identity dimension, with no detectable delay in puberty but with an altered reproductive output, while the female holds the somato-metabolic dimension and instead defers reproductive maturation. This sex-divergent developmental plasticity shows the ability of each sex to absorb the same juvenile constraint along different axes. LPD males do not adapt differently from females; they adapt in the same direction with larger amplitude, which results in the compression of a sexual dimorphism normally present at steady state. Our data illustrate that mammalian sexual dimorphism is not a fixed adult property but rather a developmentally plastic phenotype.

Chronic dietary protein restriction drives a state of growth-hormone resistance: the hepatic GH receptor is preserved, but a post-receptor block lowers IGF-1 output (*15*). The biochemistry of this state has been described for decades, but how the same nutritional signal reshapes the initial sexual dimorphism remains unresolved. Our data argue against an asymmetric nutritional input. In the liver, the one organ where the sex-biased program can be read directly (*41*), the male hepatic regulatory architecture proves the more vulnerable of the two, and not only under nutritional constraint. Consistently, genetic loss of hepatic GH-receptor signaling in *Ghr*-deficient mice likewise disrupts the male-biased hepatic transcriptional program far more than the female one (*42*).

Prepubertal orchidectomy showed that the displacement of the male growth and metabolic phenotype toward the female baseline is not driven by the fall in circulating testosterone. This gonad-independence contrasts with the literature in females: in peripubertal female rats, ovariectomy de-represses circulating IGF-1 under nutritional restriction while hepatic *Igf1* mRNA stays low (*43*). We found no evidence that orchidectomy altered either circulating IGF-1 or hepatic *Igf-1* mRNA in male mice. This is consistent with a sex-asymmetric gonadal input into somatotropic control.

The feminization of the male liver is therefore only partly testosterone-driven, and two observations show why the loss of androgens cannot account for it on its own. First, the amplitudes do not match: LPD lowers circulating androgens only partially (**Fig. 5A**), yet the most strongly repressed *Mup* transcripts fall 14- to 110-fold in LPD livers. Male-biased *Mup* expression is set by the pattern of pituitary GH release rather than by androgen levels: within four days, a continuous, female-like GH profile shifts 86% of strongly male-biased and 68% of strongly female-biased liver genes at least 20% toward the opposite-sex level (*44*) Second, part of the hepatic response does not need the testes at all: several dimorphic transcripts kept their full diet response in castrated males (**Fig. 5J–L**). Because orchidectomy removes gonadal androgens but leaves the chromosomal sex intact, a cell-intrinsic contribution of sex-chromosome complement cannot be excluded: in gonadectomized mice, X-chromosome dosage alone shifts adiposity and hepatic lipid phenotype independently of gonadal hormones (*45*). An additional credible mechanism could be that hepatic amino-acid limitation lowers liver-derived IGF-1 (*15*), an essential feedback normally sculpting GH pulses, which results in a tonic mode that feminizes the pulsatility-read transcripts. Consistently, persistent STAT5b activation, the intracellular signature of continuous GH exposure, is by itself sufficient to feminize the male liver transcriptome, repressing male-biased and inducing female-biased genes (*46*). However, the male hepatic feminization cannot be read as damage alone. The male liver program, characterized by high BCL6 levels, is advantageous under dietary surfeit and pathogen pressure but disadvantageous under scarcity, while a female-biased, BCL6-low, configuration is the less favorable upon pathogen pressure but protective under nutrient scarcity (*25*). Under protein restriction the male therefore shifts toward the state thought to be favored under scarcity, i.e., the state the female constitutively occupies.

In our juvenile cohorts, FGF21 is strongly induced by LPD in both sexes, yet we found no evidence that hepatocyte-specific *Fgf21* deletion rescued any of the male LPD phenotypes. This contrasts, however, with the adult setting, where FGF21 is required for the metabolic and lifespan benefits of protein restriction in males, an effect that is itself strongly sex- and strain-dependent (*24*, *47*). Hepatic FGF21 does, however, pace the second step of the female reproductive delay: *Fgf21* ablation accelerates first estrus under CD, with no detectable shift in vaginal opening, and only mutant LPD females reached first estrus by P56. This is the direction the FGF21 neuroendocrine axis predicts: FGF21 overexpression delays puberty onset and blunts the preovulatory LH surge, whereas removing FGF21 signaling in the forebrain, where its co-receptor β-Klotho is confined to the suprachiasmatic nucleus, shortens the delay to ovulation after a fast (*40*).

The female liver clarifies the asymmetry. LPD females show indirect signs of reduced gonadal steroid output (delayed puberty, sparse cycling, and reduced reproductive tract maturation), yet their hepatic molecular identity is undisturbed. The same drop in sex-steroid tone therefore integrates differently in the two sexes: in the male, falling androgens contribute to hepatic feminization, whereas in the female, the inferred fall in estrogens does not masculinize the liver.

How these two inputs (reduced androgen tone and the nutritionally driven shift of the somatotropic axis) are integrated at the systemic scale remains to be investigated. LPD females instead postpone maturation until a somatic threshold is reached: we found no evidence that body weight at vaginal opening differed between diets, nor of a lasting deficit in conception or cumulative output after dietary repletion, beyond a transient imprint on litter size, consistent with a somatic gate that is reached later rather than moved. This response is consistent with the energetic landscape of mammalian reproduction, where about 90% of the energy cost of reproduction is indirect metabolic overhead borne by the mother, rather than energy deposited in the offspring, even before lactation is counted (*33*). The reproductive readouts add a parallel: in both sexes the cost falls on per-cycle conception rather than on litter quality, and it is of the same order. Two divergent developmental trajectories thus converge on a single endpoint. This is unlike gestational protein restriction, which is imposed and lowers fetal and birth weight instead (*48*, *49*).

Read this way, the compression is the developmental signature of condition-dependent sexual dimorphism (*7*): the more elaborated, costlier a sex-specific program is (here, the male’s) the more sensitive to resource limitation, so dimorphism contracts precisely when resources are scarce. This is the prediction of sexual-selection theory, where more dimorphic traits show greater sex differences in condition dependence (*50*), and it parallels the greater nutritional plasticity of male size under male-biased dimorphism (*9*). The principle generalizes beyond mammals, and the seahorse makes the point with the sexes reversed: because the male bears gestation, juvenile caloric restriction stunts male somatic growth and brood-pouch development while sparing ovarian mass (*51*). Vulnerability tracks the sex carrying the larger juvenile anabolic demand, not the male sex as such. Here, we expand this ecological and macroscopic description to tissue-level, metabolic and molecular traits.

The post-weaning window coincides with the postnatal installation of the sex-specific growth trajectories and hepatic identity besides male and female reproductive maturation. In the liver, this includes the pubertal switch to pulsatile GH and the establishment of sex-biased *Cyp* and *Mup* expression (*12*, *13*, *31*). Roughly three-quarters of male-specific liver genes are induced and about half of female-specific genes repressed between three and eight weeks of age, whereas female-specific expression is largely established before puberty (*52*). LPD applied within this window therefore appears to prevent the installation of the male program rather than perturb an already-built state in females. Adult-onset protein restriction, by contrast, targets an established state and produces phenotypes dominated by energy-balance adjustment, with no compression of male hepatic identity reported (*24*, *26*, *47*). Re-analysing the published Green et al. adult data through our pipeline is consistent with this: the GH/STAT5b dimorphism axis is being selectively spared in adults while the nutrient-stress response is retained. The contracted dimorphism is most parsimoniously read as an arrest of installation rather than an erosion of a finished state, and not of the hepatic program alone but equally of the somatic growth trajectory and the metabolic phenotype, shaped across the same window.

### Limitations

Several limitations bound these conclusions. First, the design contrasts a single low-protein formulation with a single control, so dietary protein content and the protein-to-carbohydrate ratio vary together along one colinear axis; we therefore show that juvenile protein dilution compresses dimorphism but cannot resolve the dose–response shape or place the response on a nutritional reaction surface, which would require several P:C ratios within a geometric framework (*20*). Second, the assignment to the P21–P56 window rests on convergence with developmental data rather than on a factorial dissociation of age from dose and duration: the adult cohort we reanalyzed differs from ours in age, protein dose and exposure duration at once, so that comparison alone cannot isolate the window. Third, our reproductive readouts are proximate output under non-competitive mating, not fitness: the adaptive reading of the compression is an inference consistent with life-history theory, not a demonstration of selective value. Fourth, males were mated with larger outbred OF1 females, although preserved litter quality argues against a global gametic defect, the male conception deficit may partly reflect this size mismatch rather than reproductive physiology alone. Most fundamentally, our data identify a reallocation but do not localize the trade-off that drives it. The deeper axis is plausibly reproduction versus lifespan along a fast–slow pace-of-life continuum, on which the sexes are expected to hold different optima under a shared genome, and which is thought to be mediated, in part, by the very GH/IGF-1 nutrient-sensing pathway we dissect here (*53*). Resolving where each sex sits on that continuum, and how sexual conflict over it is settled, lies beyond a study that measures a truncated reproductive span without survival.

### Conclusions and future directions

Under protein dilution in the post-weaning period, the magnitude of mammalian sexual dimorphism is thus a developmentally plastic quantity rather than a fixed adult property, extending the principle of condition-dependent dimorphism (*7*), long studied in insects and ectotherms, to the mammalian somatic, hepatic, and reproductive program. That the contraction is built within a defined postnatal window, not merely revealed, reframes the amplitude of dimorphism as an outcome of the juvenile nutritional environment acting on a sex-asymmetric regulatory architecture, with the male the more sensitive sex. This recasts a trait long treated as a fixed species character as a conditional, environmentally tuned one. Whether the compression carries a net cost or an adaptive benefit, and where each sex ultimately sits on the pace-of-life continuum, are the questions this work opens.

## MATERIALS AND METHODS

### Animals

All animal experiments were approved by a local animal care and use committee (CECCAPP) and authorized by the French Ministry of Research (APAFIS #38677-2022040517218744 v6, #44718-2023051708064873 v7, #53298-2024120217028336 v3), and were carried out at IGFL (Lyon, France) in accordance with European Directive 2010/63/EU. C57BL/6N mice were purchased from Charles River (France) at 21 days of age, or at 14 days with their mothers. Hepatocyte-specific *Fgf21* knockouts (*Fgf21*^HepKO^) were generated as previously described (*28*). Mice were housed in Innorack IVC Mouse 3.5 disposable cages (Inovive) on a 12:12 light-dark cycle, with water and food ad libitum. At weaning, mice were allocated to diet, and to surgery in the orchidectomy cohort, so that each litter was spread as evenly as possible across groups with sexes balanced; genotype in the *Fgf21*^HepKO^ cohort followed Mendelian inheritance.

### Diets and general design

At postnatal day 21 (P21), mice were weaned onto a standard AIN-93G control diet (CD; 3.82 kcal/g; 18.5% kcal from protein; SAFE Diets, U8978A01R 00260) or a matched-lipid low-protein variant (LPD; 3.62 kcal/g; 4.8% kcal from protein; SAFE Diets, U8978A01R 00261). Full compositions are in **Table S1**. Body weight and naso-anal length were recorded weekly from P21 to P56 under brief isoflurane anesthesia (3%, Isoflurin, VetCare). Unless stated otherwise, mice were kept on their diet ad libitum and tissues collected at P56.

Sample size was set a priori on the Sex × Diet interaction, with body size as primary readout: for an interaction of Cohen’s d = 2 at two-sided alpha = 0.05 and 80% target power, the F-test of the interaction term requires n = 10 per cell (N = 40), for an effective power of 87%. The pre-specified hypotheses were that juvenile protein restriction reduces sexual dimorphism in somatic growth and in hepatic gene expression, tested as the Sex × Diet interaction, and that this compression is larger in males. The analyses of pubertal timing, of reproductive output and of the orchidectomy cohort were also pre-specified. All other analyses, including the feminization score, the trait-wise decomposition of the Mahalanobis distance, the growth-rate threshold and the androgen-dependence index, are post hoc and are identified as such below.

### Cohorts

**Cohort 1** (n = 40, 10 per Sex × Diet). Transferred at P36 to the ANIPHY platform (Lyon), acclimated 3 days; body composition at P38 by low-field NMR (Minispec LF90 II, Bruker); indirect calorimetry P39 to P41 (PhenoMaster, TSE Systems); returned to IGFL at P42; insulin tolerance test at P49. One LPD male was excluded from the entire calorimetry recording: its cumulative food intake was four times the next highest value, a hopper artefact rather than an eating phenotype; it is kept in all other analyses. At P56, mice were fasted 6 h from 8 AM, given buprenorphine (0.1 mg/kg s.c., Buprecare) 30 min and glucose (2 g/kg by gavage) 15 min before terminal anesthesia (ketamine 100 mg/kg i.p. plus subcutaneous lidocaine). Portal and caval blood were collected by midline laparotomy into Z-gel microtubes (Sarstedt), centrifuged (10,000g, 5 min, 4°C) and stored at −80°C. Liver was snap-frozen, femur measured with calipers, and white and brown adipose depots, kidneys and gastrocnemius weighed. The liver transcriptome was profiled on 20 of these mice (5 per group), and portal and caval serum on the 19-amino-acid panel.

**Cohort 2** (n = 42, 10 to 11 per Sex × Diet). Oral glucose tolerance test at P42 with serum insulin at 0, 15 and 30 min; daily monitoring of balanopreputial separation in males and of vaginal opening, first estrus and cyclicity in females. At P56, mice were fasted from 8 AM and killed by cervical dislocation between 2 and 4 PM; submandibular blood was collected, organs (including testes, seminal vesicles, uterus) weighed, and hepatic glycogen and triglycerides measured. Epididymal sperm count was performed in males. The liver metabolome was profiled on 24 of these mice (6 per group).

**Cohort 3** (n = 28, 7 per Sex × Diet). Postprandial tail-vein blood at P35, P42 and P49 for longitudinal hormone profiling. At P56, postprandial portal and caval blood were collected between 8 and 10 AM, immediately followed by cervical dislocation, and analyzed on the amino-acid panel.

**Cohort 4** (n = 83; F-CD 21, F-LPD 20, M-CD 21, M-LPD 21). Allocated at weaning to sacrifice at P28, P42 or P70, between 8 and 10 AM without fasting. Tail-vein blood at P28, P35, P42 and P49 for hormone profiling; liver taken for RT-qPCR of a sex-dimorphic panel.

**Cohort 5** (females, n = 28, 14 per diet; dietary rescue). Kept on CD or LPD from P21 to P70, then all on CD. Vaginal opening, first estrus and cytology were recorded daily, cyclicity between P56 and P70. From P70 each female was paired with a proven C57BL/6J stud for about three months; latency to first litter, per-parity litter size and per-pup birth weight were recorded.

**Cohort 6** (males, n = 40, 10 per Diet × Surgery). Bilateral orchidectomy (ORX) or sham surgery at P21, weaning onto CD or LPD the same day. Buprenorphine (0.1 mg/kg i.p.) 30 min before induction; isoflurane 5% for induction, 1 to 2% for maintenance. After aseptic preparation, a 2-mm ventral midline scrotal incision was made, the muscle layer incised, the spermatic vessels clamped and each testis excised after cord transection; sham animals received the incision only. Muscle and skin were closed with resorbable suture, and buprenorphine (0.1 mg/kg s.c.) was given every 8 h on the day of surgery and the next, and longer if a structured pain grid indicated residual discomfort. OGTT at P42; sacrifice at P56 at 8 AM after submandibular blood collection.

**Cohort 7** (*Fgf21*^HepKO^, C57BL/6J background; n = 92, 8 to 14 per Sex × Diet × Genotype). Alb-Cre^+/−^ *Fgf21*^fl/fl^ mice were crossed with Alb-Cre^−/−^ *Fgf21*^fl/fl^ controls to give littermates of both sexes, weaned at P21 onto CD or LPD. Tail-vein blood at P28 and P56 for IGF-1 and at P56 for FGF21; daily balanopreputial separation and vaginal opening; OGTT at P42; sacrifice at P56 between 8 and 10 AM without fasting.

**Cohort 8** (females, three-arm reproductive output). From P21, females were allocated to CD throughout, LPD throughout, or Switch: CD at baseline, a pulsed transition to LPD at day 5 of each mating cycle and a return to CD at parturition, so that conception occurs on CD and only gestation is exposed to LPD. From P70, each female was paired with a confirmed-fertile B6N male for four consecutive nights per cycle, over up to two cycles. Maternal weight, litter size, perinatal mortality and per-pup birth weight were recorded at P0; conception was inferred from gestational weight gain and confirmed by delivery.

**Cohort 9** (males, non-competitive reproductive output). Males raised on CD or LPD from P21 to P70 were paired at P70 with individual age-matched virgin hyperfertile OF1 females (Charles River) in single-male cages, over up to three cycles at about 21-day intervals, each cycle being four consecutive nights in the female’s cage; males returned to their diet-segregated cage between cycles. Gestation was tracked by maternal weight; litters were counted, sexed and weighed at P0.

### Reproductive development

#### Females

Mice were examined daily from weaning. Puberty onset was the day of vaginal opening, after which estrous stage was assessed daily by vaginal cytology (*38*); first estrus was the first appearance of cornified epithelial cells. In Cohort 5, cyclicity was counted between P56 and P70 as complete transitions through proestrus, estrus, metestrus and diestrus. Uterus weight at sacrifice indexed reproductive maturation.

#### Males

Puberty onset was assessed daily by balanopreputial separation (*29*, *54*), and testis and seminal vesicle weights at P56 indexed testicular growth and androgenic status. Epididymal sperm count followed (*55*): epididymides stored at −80°C were thawed, weighed, minced in 1 mL PBS and shaken, and intact sperm heads counted under brightfield microscopy.

### Metabolic phenotyping

#### ITT

At P49, Cohort 1 mice were fasted 6 h from 8 AM and injected with human insulin (0.5 U/kg i.p.; NovoRapid, Novo Nordisk). Tail-vein glucose was measured with a glucometer (OneTouch Verio Reflect, LifeScan) before gavage ant at 15, 30, 45, 90 and 120 min. The humane endpoint was blood glucose below 30 mg/dL: the test was stopped for that animal, the remaining timepoints recorded as missing and glucose given as rescue (1 g/kg i.p.). Four LPD mice reached it.*OGTT.* At P42, Cohort 2 mice were fasted 6 h from 8 AM and given D-glucose by gavage (2 g/kg, 20% solution at 5 mL/kg; Sigma G7021) with the same sampling schedule. Extra blood was collected at 0, 15 and 30 min for serum insulin.

#### Circulating hormones

IGF-1 (Mouse/Rat IGF-1 Quantikine, R&D Systems MG100), FGF21 (Mouse/Rat FGF21, BioVendor RD291108200R) and insulin (UltraSensitive Mouse Insulin, Crystal Chem 90080) were measured by ELISA according to the manufacturers’ protocols.

#### Serum steroids

Testosterone (along with androstenedione, progesterone, corticosterone and 11-deoxycorticosterone) was measured at P56 in the orchidectomy cohort by liquid chromatography coupled to tandem mass spectrometry on a 6495D triple quadrupole (Agilent), on 100 µL of serum. The limit of quantification was 0.13 nmol/L for testosterone. Intermediate precision at two control levels is reported in the Supplementary Methods. Only testosterone is reported here; the four other analytes are in the deposited dataset.

### Liver biochemistry and histology

#### Triglycerides

Snap-frozen liver (50 to 100 mg) was homogenized in 0.25 M sucrose, 10 mM HEPES with a Precellys 24 homogenizer (Bertin), and triglycerides were measured by a glycerol-3-phosphate oxidase colorimetric assay (Biolabo 87319), expressed per g liver wet weight.

#### Glycogen

Liver (50 to 100 mg) was homogenized in 6% perchloric acid and neutralized to pH 6.5 to 8.5 with 3.2 M K_2_CO_3_. Glycogen was hydrolyzed with alpha-amyloglucosidase (1 mg/mL, 1 h, 45°C), the released glucose phosphorylated by hexokinase (150 U/mL, 30 min, room temperature) and oxidized by glucose-6-phosphate dehydrogenase (200 U/mL) with coupled NADP^+^ reduction; NADPH absorbance was read at 340 nm (SPECTRO Star nano, BMG Labtech). Glycogen was expressed per g liver wet weight.

#### Oil Red O

Livers from 19 mice (5 per group, except F-LPD n = 4) were dehydrated overnight in 30% sucrose, embedded in OCT and stored at −80°C. Sections (10 µm, Leica CM 3050S) were fixed in 4% paraformaldehyde and stained with Oil Red O (Sigma O-0625). Ten random fields per mouse were imaged at 20× (Leica DM6000). Droplets were segmented in Fiji with a modified macro of Nishad et al. (*56*) (threshold 0 to 95, circularity 0.70 to 1.00, size 10 to 250 µm^2^) and summarized per mouse. Acquisition and analysis were single-blind to diet.

### RT-qPCR

Total RNA was extracted with the NucleoSpin RNA kit (Macherey-Nagel 740955.250) and quantified on a NanoDrop 2000. Reverse transcription used 1 µg of RNA (SensiFAST cDNA Synthesis, Meridian BIO-65053) and qPCR used Takyon No ROX SYBR MasterMix blue dTTP (Eurogentec UF-NSMT-B0701) on a Bio-Rad CFX96. Analysis followed the MIQE 2.0 guidelines (*57*); expression was normalized to two validated reference genes (*Rpl32*, *Actb*), stability assessed with geNorm, reported on the log_2_ scale and calibrated to the mean of the M-CD (or CD-Sham) group. Primers are listed in **Table S2**. The Mup cluster is a pan-*Mup* readout, amplifying the cluster rather than a single paralog.

### Urinary MUP Western blot

Urine collected at P56 was stored at −20°C. For each sample, 3 µL of urine was mixed with 8.1 µL of denaturing buffer (20% glycerol, 10% 2-mercaptoethanol, 10% SDS, 62.5 mM Tris) and 18.9 µL of water and denatured at 95°C for 10 min; 10 µL was loaded onto a 4 to 20% Mini-PROTEAN TGX precast gel (Bio-Rad) and run at 200 V for 30 min. Proteins were transferred to nitrocellulose (Trans-Blot Turbo, Bio-Rad) and total protein content was verified. Membranes were blocked for 1 h in TBS-T with 5% milk, incubated for 1 h with anti-MUP (C-7) mouse monoclonal IgG1 (Santa Cruz sc-166429, 1:500), washed three times in TBS-T, incubated for 20 min with StarBright Blue 700 goat anti-mouse IgG (Bio-Rad 12004159; 1:2,500) and washed three times in PBS-T. Blots were imaged at 660 nm on an Amersham ImageQuant 800 (Cytiva). **Fig. 4F** shows the ladder and the six male lanes cropped from a single membrane that also carried the female lanes; a global rotation, the crop and a linear inversion and windowing were the only operations, with no gamma adjustment, local editing or lane splicing, and the uncropped acquisition is shown in **Fig. S8**.

### Serum amino acids

Amino acids were quantified by HPLC on an Agilent 1100 system with a Zorbax Eclipse-AAA C18 column (3.5 µm, 150 × 4.6 mm) and a guard cartridge, using a method developed for this setup. Samples were buffered with borate at pH 10.2 and derivatized at room temperature with ortho-phthalaldehyde for primary amines and 9-fluorenylmethyl chloroformate for secondary amines (Agilent 1313A autosampler). Separation was at 40°C, 2 mL/min, with 40 mM NaH_2_PO_4_ pH 7.8 (eluent A) and acetonitrile/methanol/water 45:45:10 (eluent B), on a 20 to 80% B gradient; detection was by fluorescence at 340/450 nm (266/305 nm for proline). Cystine was not detected, so 19 amino acids were quantified, with norvaline as internal standard and response factors from a 250 pmol/µL standard mix (ChemStation for LC 3D). Four datasets were analyzed separately: postabsorptive portal and caval serum (Cohort 1, 6-h fast) and postprandial portal and caval serum (Cohort 3, non-fasted).

### Liver metabolomics

Polar metabolites were extracted from about 30 mg of frozen liver in 80% methanol with sonication and homogenization (Precellys Cryolys), clarified (15,000 rpm, 15 min, 4°C), dried and reconstituted in methanol proportionally to pellet protein content (BCA). Profiling used UHPLC-MS/MS in multiple-reaction-monitoring mode on a 6495 iFunnel triple quadrupole (Agilent), in two chromatographic modes with positive and negative electrospray ionization, covering central carbon metabolism (*58–60*). Data were processed in MassHunter B.07.00; features with a coefficient of variation above 30% across QC samples were excluded. The 189-feature by 24-sample table was deduplicated across ionization modes by keeping, for each duplicated compound, the copy with the lower coefficient of variation (ties within 5% broken by higher mean signal), leaving 165 metabolites, which were log_2_-transformed. Ambiguous identifiers (for example beta-alanine/sarcosine, UDP-hexose) were kept as such. Per-metabolite tables and the deduplication log are in **Data S1**.

### Liver RNA sequencing

RNA integrity and concentration were checked on a Qubit 4.0 and a TapeStation 4150, and mRNA purified from 1 µg of total RNA (Poly(A) mRNA Magnetic Isolation Module, NEB). Libraries were prepared with the CORALL Total RNA-Seq kit (Lexogen), fragmentation-free and carrying 12-bp unique molecular identifiers, and sequenced on a NextSeq500 (Illumina), single-end 1 × 86 bp: 31 to 44 million reads per sample. Adapters were trimmed with Cutadapt v3.5, reads mapped to GRCm39 (Ensembl 110) with STAR v2.7.11a, deduplicated with UMI-tools v1.1.2 and counted with htseq-count v2.0.3. Statistical modelling is described below.

### Quantification of dimorphism

The same framework was applied at three levels: five somato-metabolic traits (**Fig. 2J**), 165 hepatic metabolites (**Fig. 3C**) and the liver transcriptome (**Fig. 3I**). Each sample-by-feature matrix was reduced, where features outnumbered samples, by principal-component analysis on the centered matrix, retaining the fewest leading components reaching 80% cumulative variance. Within each diet, sex separation was tested by PERMANOVA on Euclidean distances (`vegan::adonis2`; 9,999 permutations), and the male-female distance was computed as a Mahalanobis distance:

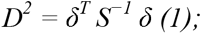

where δ is the vector of differences between male and female group means within a diet and S is the within-group covariance matrix pooled across the four Sex × Diet cells. Compression is reported as:

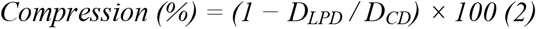

and tested by permuting sex labels within each diet (9,999 permutations) on the statistic D^2^ – D^2^_LPD_; 95% confidence intervals come from a stratified bootstrap within each cell.

For the somato-metabolic level, five candidate traits from Cohort 2 (body weight at P56, fasting glycemia and fasting insulin at P42, hepatic glycogen and hepatic triglycerides at P56) passed a pre-specified two-step selection performed on CD animals only: a dimorphism filter (|Hedges’ g| > 0.5 and Welch p < 0.05) and a collinearity filter (one of any pair with |r| > 0.80 retained, decision logged). All five were retained. Robustness was assessed by re-running the compression on every combination of three to five of them. For the transcriptome, two feature sets were analyzed in parallel: the full prefiltered transcriptome and the sex-biased set, defined as genes with adjusted p < 0.05 in the sex contrast under CD alone, a selection orthogonal to the compression hypothesis and therefore non-circular; sensitivity analyses used the top 500 and top 1,000 genes by absolute Wald statistic.

#### Feminization score (post hoc)

Each liver sample was projected onto a sex-discriminating axis built in PC space from the strictly sex-biased set (adjusted p < 0.05 and |shrunken log_2_ fold-change| > 1 in the CD sex contrast; 321 genes). The axis is the Fisher linear discriminant between M-CD and F-CD, learned on CD samples only: CD samples were scored by leave-one-out cross-validation and LPD samples projected out of sample. Scores were rescaled so that the M-CD median is 0 and the F-CD median is 1 (Supplementary Methods). Group contrasts used aligned-rank-transform ANOVA (`ARTool` v0.11.2) with Holm correction, and two one-sided one-sample tests against the midpoint 0.5 were applied to the M-LPD group, for majority feminization and for non-inferiority.

#### Androgen-dependence index (post hoc)

In the orchidectomy cohort, per-gene expression was fit by a four-group cell-means model (`limma::lmFit` on ∼ 0 + Group, empirical Bayes moderation), from which four contrasts were extracted: the diet effect within each surgery, the surgery effect under CD, and the Diet × Surgery interaction. For each gene, ρ is the ratio of the diet log_2_ fold-change under ORX to the diet log_2_ fold-change under Sham: ρ near 1 indicates an androgen-independent diet response, ρ near 0 a response already saturated by orchidectomy, and ρ below 0 a sign reversal. Per-gene values are in **Data S3**.

### Statistical analyses

Analyses were performed in R v4.4.3. Software, package versions and their citations are listed in the Supplementary Methods; resampling and permutation used a fixed seed.

#### Reporting convention

α = 0.05, is used as a landmark, not as a verdict: results are graded by the strength of the evidence rather than split into significant and non-significant (*61*), and an effect we did not detect is reported as an absence of evidence, not as evidence of absence. Estimated marginal means are given with 95% confidence intervals and contrasts with a standardized effect size. Each symbol in the figures marks a single comparison, so no correction for multiple testing applies; where a family of tests is larger, the correction is named with the analysis (Holm, or Benjamini-Hochberg for the omics). **Data S4** lists, panel by panel, the model, the test, N and any correction.

#### Somatic growth

Body weight and naso-anal length from P21 to P56 were modeled by linear mixed-effects regression with a natural cubic-spline basis on age:

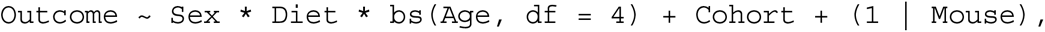

by REML with the `bobyqa` optimizer; Surgery replaced Sex in Cohort 6 and Genotype was added in Cohort 7. Type-III analysis of variance was applied to the fixed effects and estimated marginal means and within-sex diet contrasts extracted at each weigh day. Growth rate is the first derivative of the fitted spline.

#### Circulating hormones

IGF-1 and FGF21 followed approximately log-linear time courses, so no spline was used:

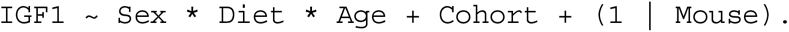

The same model was used on log(FGF21). Values below the FGF21 assay limit of detection (7 pg/mL; 60 of 244 observations) were set to half that limit for fitting, with a left-censored Tobit refit as sensitivity analysis. Cross-sectional FGF21 in Cohort 7 was modeled directly by Tobit regression with bootstrap 95% confidence intervals (2,000 resamples). Serum testosterone in Cohort 6 was left-censored at the limit of quantification (0.13 nmol/L). After orchidectomy every LPD male fell below it, so that cell contains no quantified value and a Diet × Surgery model has no identified parameters: neither the interaction nor the diet contrast within castrated males can be estimated, and neither is reported. The diet effect was therefore tested in sham males, in whom every value is quantified, by exact Wilcoxon rank-sum test with the Hodges-Lehmann shift. Detectability was compared by Fisher’s exact test on the number of males above the limit of quantification.

#### Cross-sectional endpoints and allometry

Femur length, fasting glycemia and insulin, hepatic glycogen and triglycerides and cross-sectional liver qPCR were analyzed by two- or three-way type-III ANOVA. Hepatic lipid droplets were reduced to a per-mouse geometric mean area and analyzed on the log scale, the mouse being the unit of replication. Metabolic organ weights at P56 were analyzed by log-log ANCOVA with body weight as covariate, testing the Sex × Diet or Diet × Surgery interaction explicitly, and body composition at P38 by log-log ANCOVA with backward elimination of the slope interactions, the retained model being reported on the panel. Reproductive organ weights followed a per-organ scale selection: body weight was kept as covariate when its correlation with the organ was significant and informative by F-test (testis, seminal vesicle, uterus; dropped for sperm count), and the raw, log-log and Box-Cox scales were compared by corrected AIC, leave-one-out RMSE and the DHARMa Kolmogorov-Smirnov test before the best-scoring scale was reported.

#### Tolerance tests

Glycemic time courses were modeled by

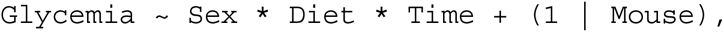

with Time categorical, and serum insulin identically on three timepoints. Areas under the curve were computed by the trapezoidal rule and analyzed by 2 × 2 ANOVA; for the ITT, timepoints after the humane endpoint were imputed by last observation carried forward for the AUC calculation only.

#### Amino acids and metabolites

Serum amino acids and the 165 hepatic metabolites were log_2_-transformed and modeled with `limma` on a cell-means parameterization of the four Sex × Diet groups, each amino-acid dataset separately. Three contrasts were extracted, the diet effect within each sex and the Sex × Diet interaction, with robust empirical Bayes moderation and Benjamini-Hochberg adjustment within each contrast.

#### Liver transcriptome

Gene-level counts were analyzed with `DESeq2` v1.46.0. A single interaction model, `∼ Sex * Diet`, was fit per cohort on genes with at least five counts in at least five samples, and five contrasts were derived from that single fit so that they share one dispersion estimate: the male diet effect, the female diet effect (`Diet_LPD_vs_CD + SexF.DietLPD`), the Sex × Diet interaction (signed so that positive values indicate a larger female response), and the sex effect under each diet. Multiple testing was controlled within each contrast by the Benjamini-Hochberg procedure. The Wald test of the interaction was complemented by a likelihood-ratio test of `∼ Sex * Diet` against `∼ Sex + Diet`, re-fit on the same matrix with the same size factors; the conservative interaction set used throughout is the intersection of the two. Differentially expressed genes and all counts reported in the figures use unshrunk log_2_ fold-changes with adjusted p < 0.05 and |log_2_ fold-change| > 1. Shrunk estimates (`apeglm` v1.28.0 for the three named coefficients, `ashr` v2.2-63 for the two compound contrasts) were used for display coordinates only, never for inference. Ensembl identifiers were mapped to MGI symbols with `org.Mm.eg.db` v3.20.0. Differential-expression tables are in **Data S2**.

#### Cross-study comparison

The liver RNA-seq dataset of Green et al. (*24*) (<u>GSE181301</u>; 24 adult C57BL/6J samples, 6 per Sex × Diet) was re-analyzed downstream from the published RSEM expected counts, rounded to integers, with the same QC, model, contrasts and thresholds.

#### Gene sets and enrichment

Six curated sets of at most 15 genes each were assembled from the literature, two for the nutrient-stress arm (ATF4 and PPARα targets (*62–67*)) and four for the GH-driven sex-biased arm (STAT5b targets, BCL6 targets, GH-axis dimorphism, hepatic feminization (*12*, *13*, *23*, *52*, *68*, *69*)); membership is in Supplementary Methods and deposited as CSV files. Enrichment among DEGs was tested by a one-sided Fisher exact test within the prefiltered background, against the male, female and interaction contrasts separately. Gene set enrichment analysis used mouse-native Hallmark sets (`MSigDB MH`, `msigdbr` v25.1.1) and `fgseaMultilevel` (`fgsea` v1.32.4, sets of 15 to 500 genes, eps = 0), on three rankings by unshrunk Wald statistic: male diet effect, female diet effect and interaction. Results are in **Data S2**.

#### Pubertal timing and cyclicity

Time-to-event endpoints were analyzed by Kaplan-Meier estimation with log-rank tests and Cox proportional-hazards models, stratified by cohort where females from Cohorts 2 and 5 were pooled. Proportionality was checked on scaled Schoenfeld residuals; where it was rejected (first estrus) a Weibull accelerated failure time model was fit and reported. Where censoring was complete or nearly so (LPD *Fgf21*^fl/fl^ females), Firth-penalized Cox regression was used. The vaginal-opening-to-first-estrus interval was compared by Wilcoxon rank-sum test with a bootstrap confidence interval on the median difference, and the number of complete cycles by Poisson GLM, with a quasi-Poisson refit and a Wilcoxon test as robustness checks.

#### Growth-puberty coupling (post hoc)

Body weight at the exact day of the pubertal event was interpolated between bracketing weigh days, and the prepubertal growth rate taken as the slope of body weight on age up to that day. Age at vaginal opening was fit against growth rate by the reciprocal model a + b / rate (Levenberg-Marquardt damped least squares), with bootstrap 95% prediction intervals (5,000 iterations). A breakpoint was located by segmented regression (0.30 ± 0.05 g/day), and the residual dispersion of the reciprocal model was compared above and below it, which is the test presented in **Fig. S6D**. Mediation of the diet effect on age at vaginal opening by body weight was estimated in the potential-outcomes framework (*70*) with a Bayesian bivariate Gaussian system (`brms`, 4 chains × 4,000 iterations, weakly informative empirically calibrated priors, right-censored ages handled by `cens()`), and total, natural direct and natural indirect effects obtained by G-computation over the posterior; convergence and priors are detailed in Supplementary Methods.

#### Reproductive output

Conception was analyzed per mating cycle by logistic regression, with a per-animal random intercept where non-singular, and persistence by a binomial GLM on per-male success counts; a pre-specified sensitivity analysis excluded OF1 females that failed with all partners. In the three-arm cohort, cycles were classified by the effective diet at fertilization, with a planned carryover contrast of Switch against CD at parity 2. Conditional outcomes among confirmed deliveries (total births, live pups, per-pup weight) were modeled by Diet × Parity linear models, and perinatal mortality by a binomial GLM with a Diet effect only, because rare events caused complete separation in the full design. In the dietary-rescue cohort, latency to first litter was compared by Wilcoxon rank-sum test, litters per dam by Poisson GLM, and per-parity litter size and pup weight by linear mixed models with a random intercept by dam. Marginal output per cycle was estimated as the product of conception probability and conditional litter size by non-parametric bootstrap over cycles (5,000 iterations) and is reported as a ratio to CD with percentile intervals; the gap was decomposed additively into a conception-stage and a litter-size contribution.

#### Diagnostics

Residuals were assessed visually and by simulated-residual diagnostics (DHARMa). Departures from assumptions were handled case by case and logged: log transformation for severe skewness (FGF21, sperm count, droplet area), data-driven Box-Cox scale selection for reproductive organ allometry, Tobit regression for assay censoring, replaced by rank-based and exact tests where a design cell contained no quantified value, a quasi-Poisson refit for overdispersion, a Weibull accelerated failure time model when proportional hazards failed, Firth correction for monotone likelihood, non-parametric tests alongside parametric models in small samples, and last-observation-carried-forward imputation in the ITT.

## Data and code availability

All raw data, R scripts, processed outputs, and metadata are available on Zenodo: 10.5281/zenodo.16411410. Raw sequencing data (FASTQ files) have been deposited in the Gene Expression Omnibus (GEO) under accession number GSE302336 and are publicly available.

## ACKNOWLEDGMENTS

We thank Jean-Louis Thoumas for assistance with mouse handling and strain management. We are grateful to Raghav Galgali for his contributions to liver transcriptome analysis and sperm count assays. We also acknowledge Manuela Forero for her work on *Fgf21* knockout mouse experiments, and Lucie Cressot for her work on the ORX cohort.

We thank Daniela Fernandois and Marialetizia Rastelli for their help in setting up the puberty-related assays and their contributions to the interpretation of the data.

We thank Julijana Ivanisevic, Hector Gallart Ayala, and Tony Teav (Metabolomics Unit, University of Lausanne) for performing targeted liver metabolomics and for helpful discussions regarding the analysis and interpretation of metabolomics data.

## Use of AI

The authors used Anthropic’s Claude (Opus) to assist with improving English phrasing and grammar during manuscript preparation, and to refine and optimize R scripts used for data analysis and figure generation. All scientific content, interpretation, and conclusions were developed and verified by the authors.

## Author contributions

Conceptualization: AJ, FDV, FL

Methodology: AJ, LR, FDV, FL

Software: FDV

Formal analysis: AJ, LR, FDV, FL

Investigation: AJ, LR, AL, EC, SH, BG, IR, PDS, IP, CR, MZ

Resources: AF, JB, SH, BG, IR, PDS, IP, HG

Data curation: LR, FDV

Writing – original draft: AJ, FDV

Writing – review & editing: AJ, LR, SH, BG, AF, JB, HG, FDV, FL

Visualization: AJ, FDV

Supervision: FDV, FL

Project administration: FDV, FL

Funding acquisition: FL

---

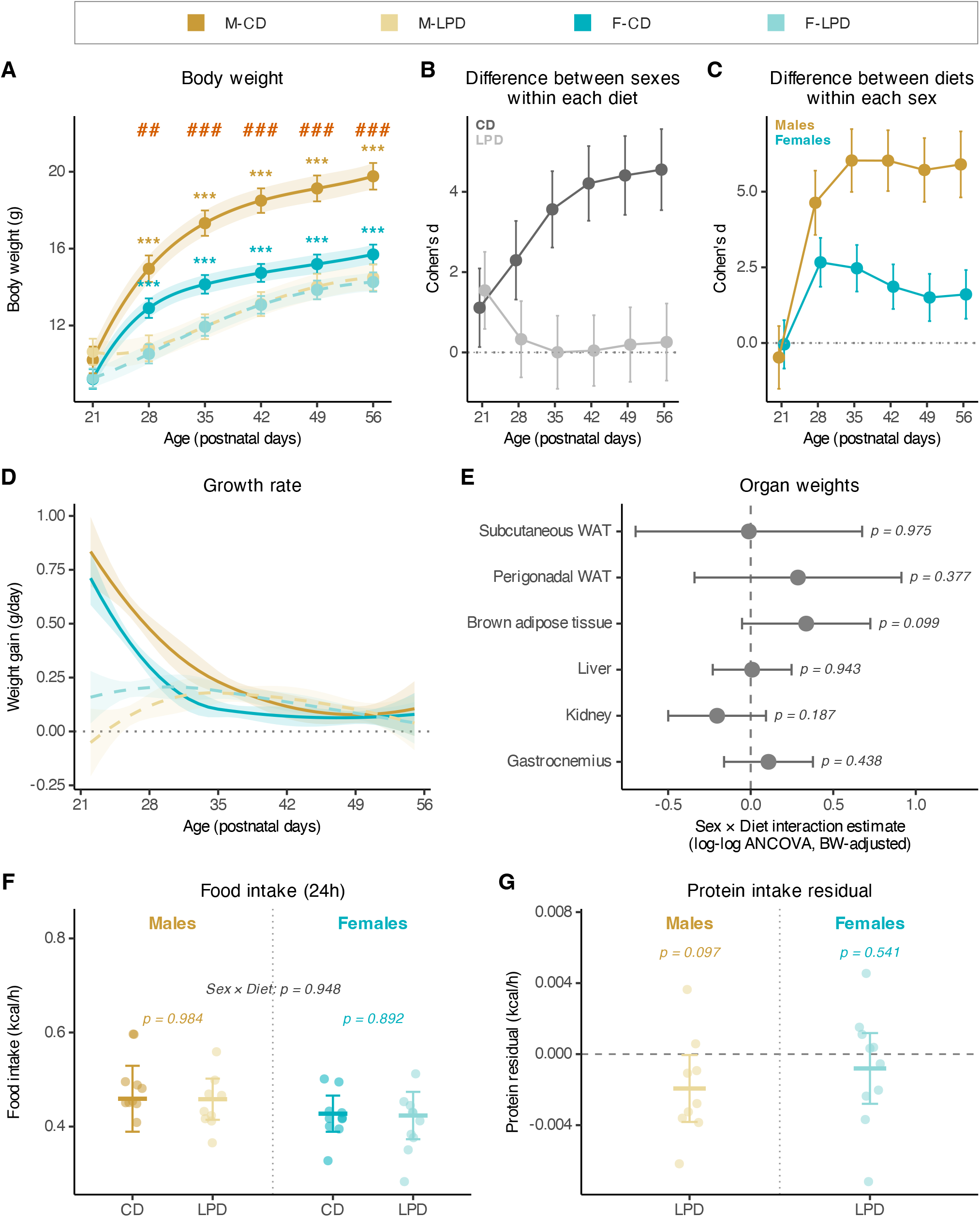

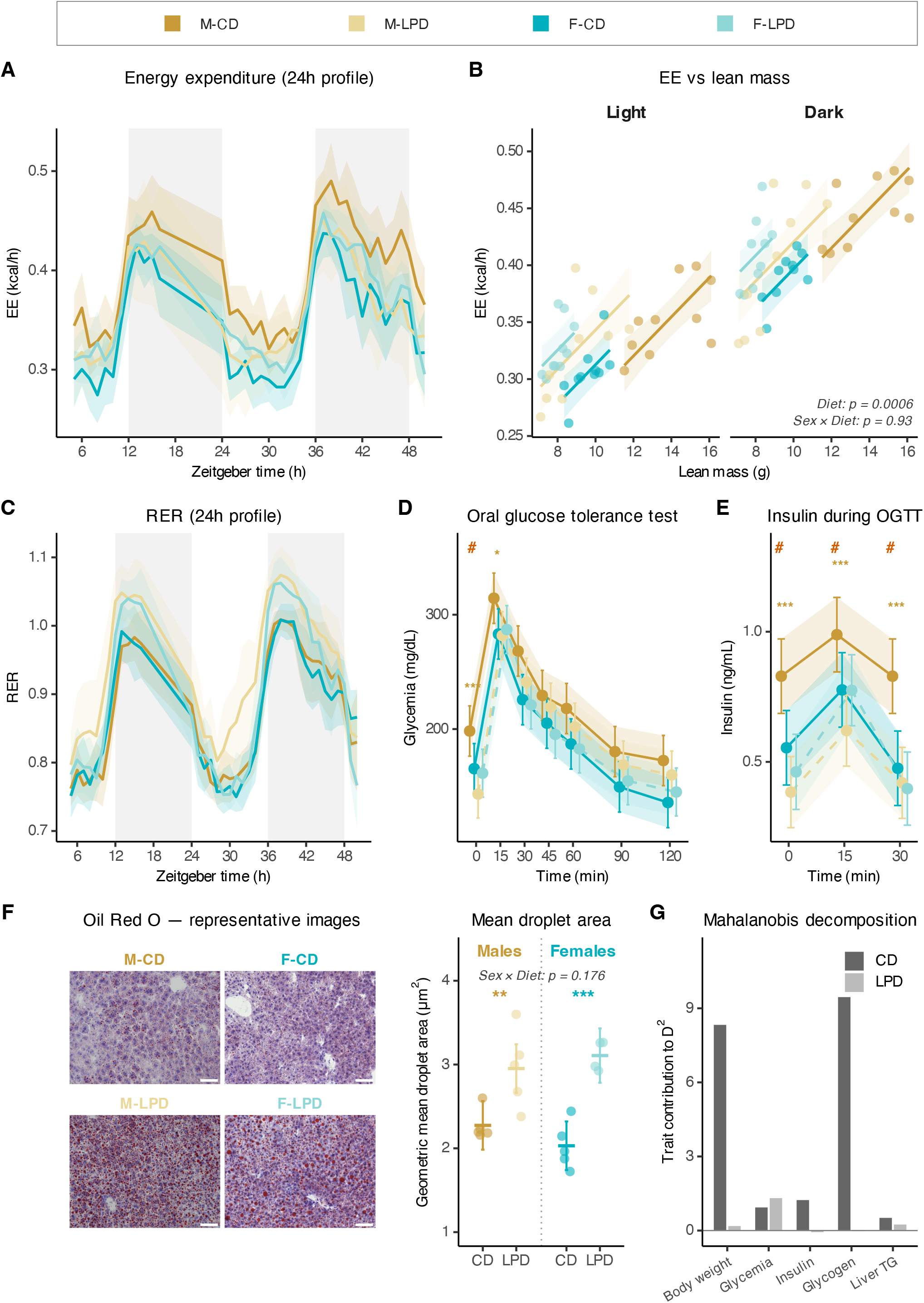

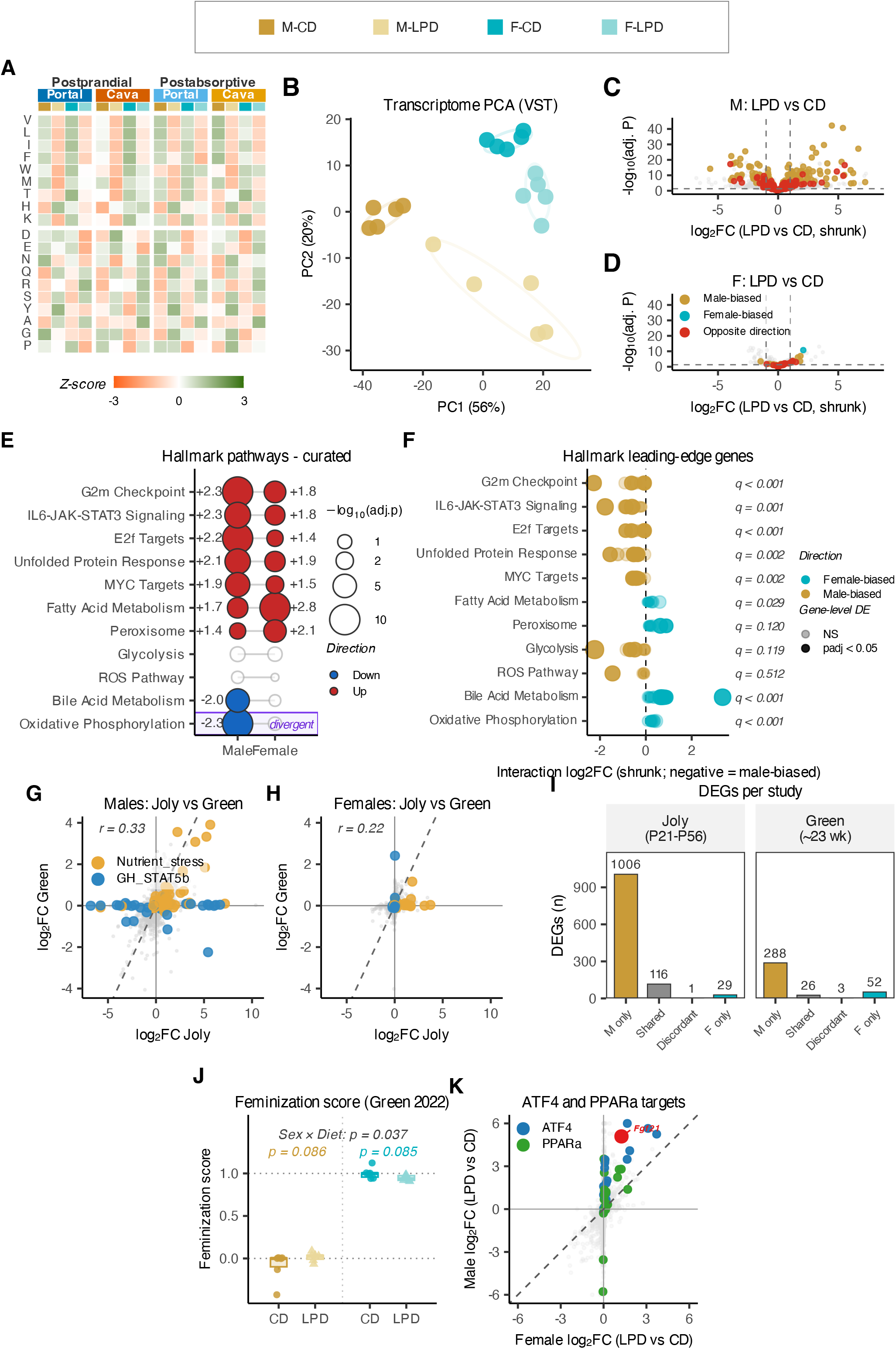

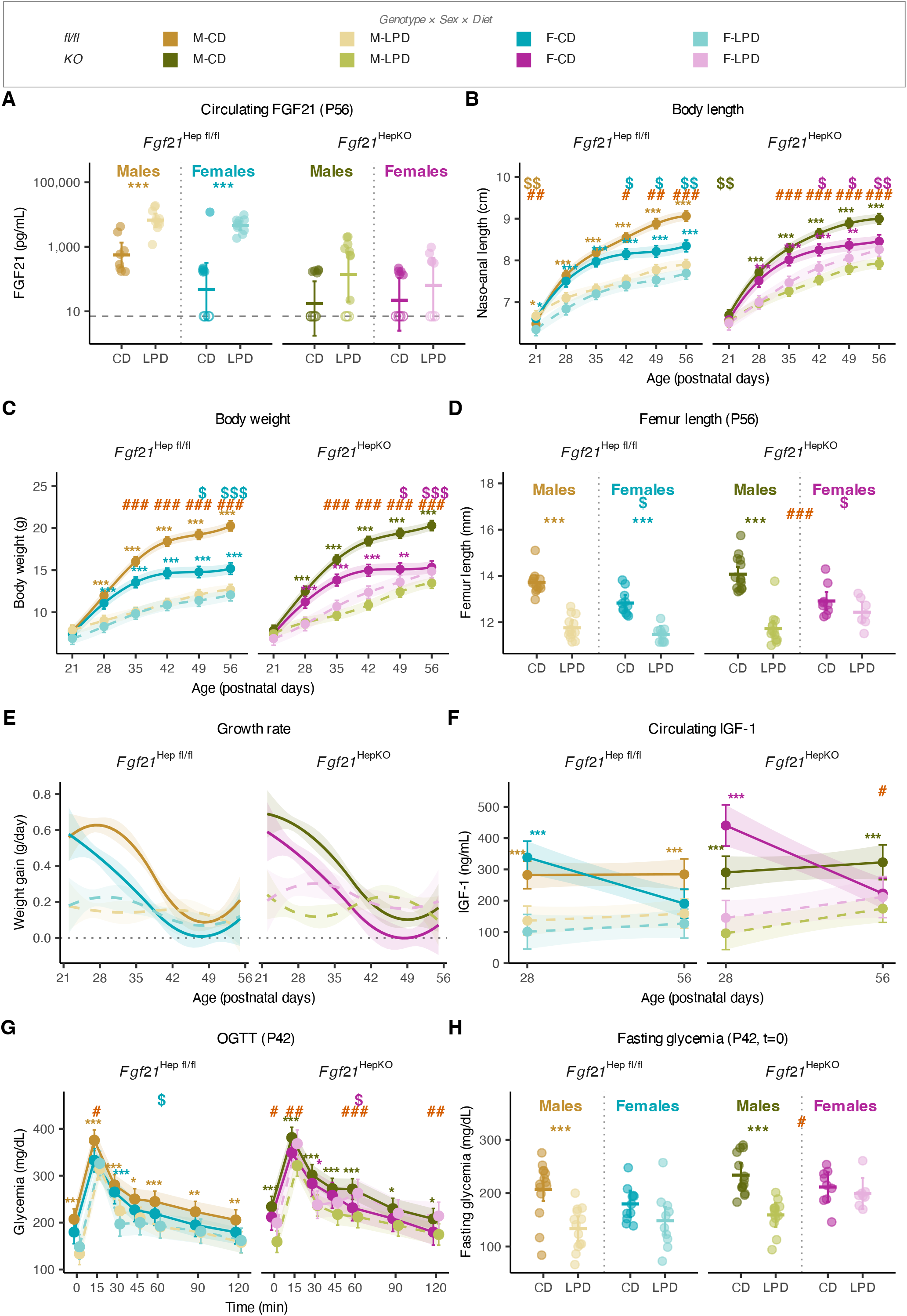

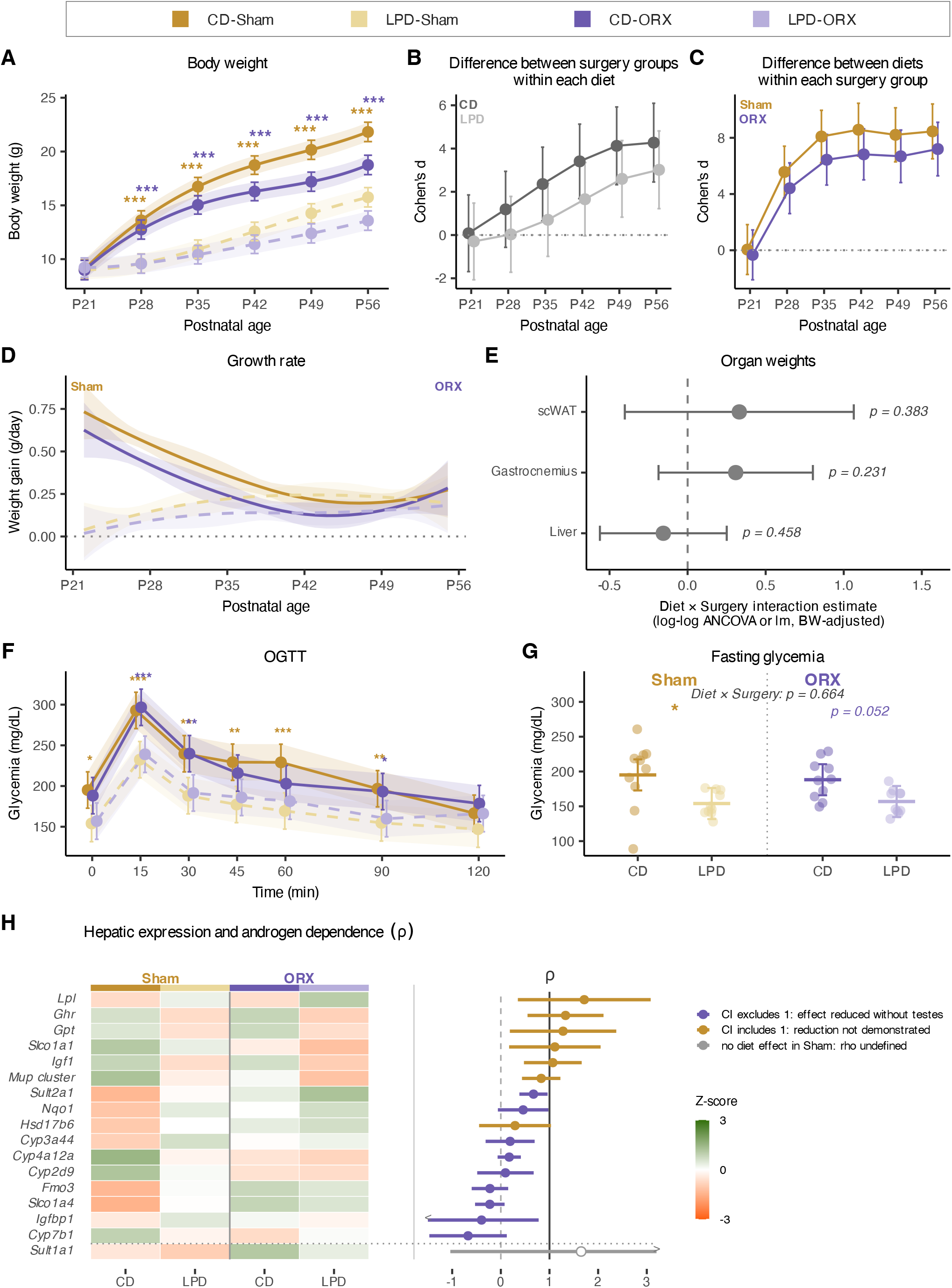

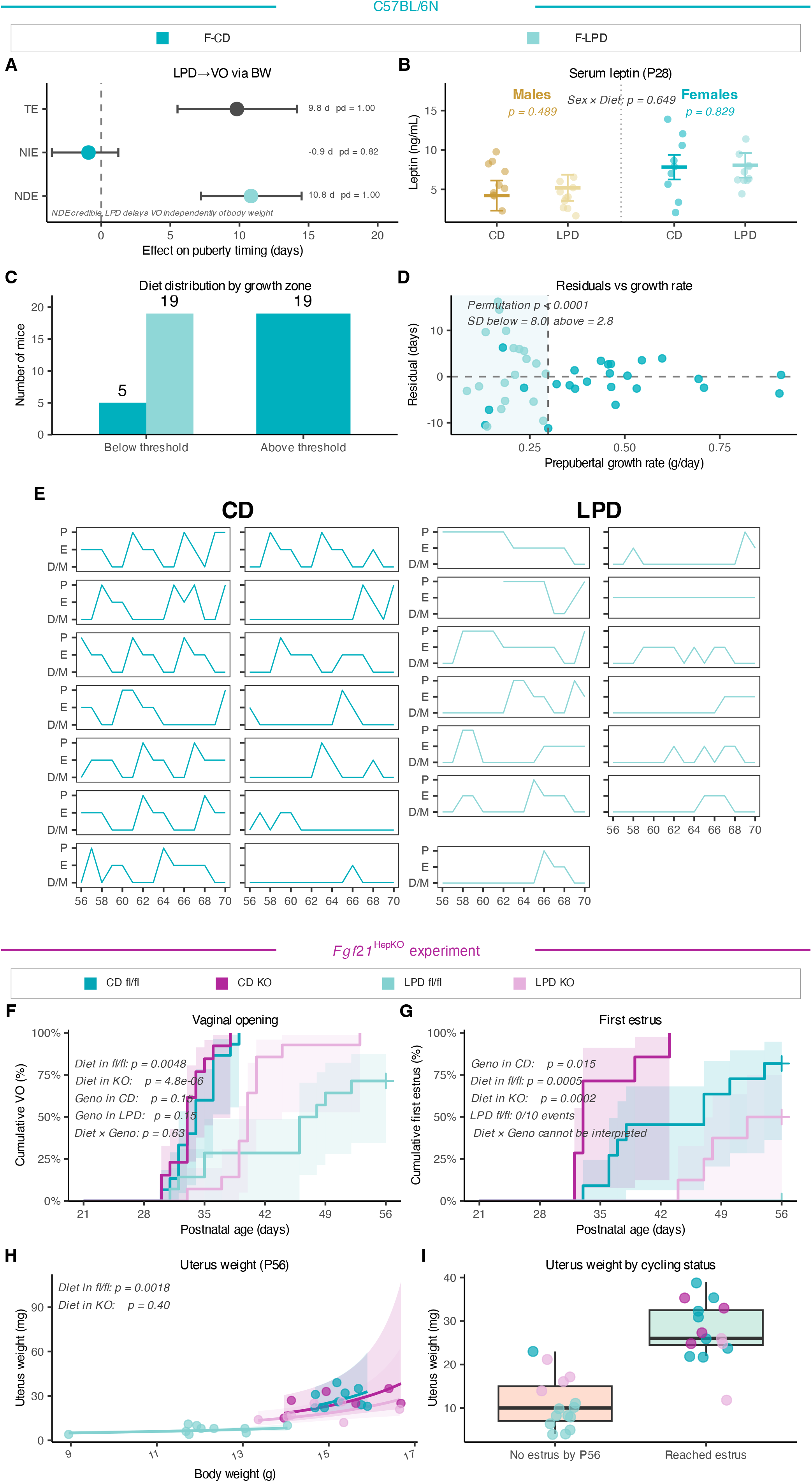

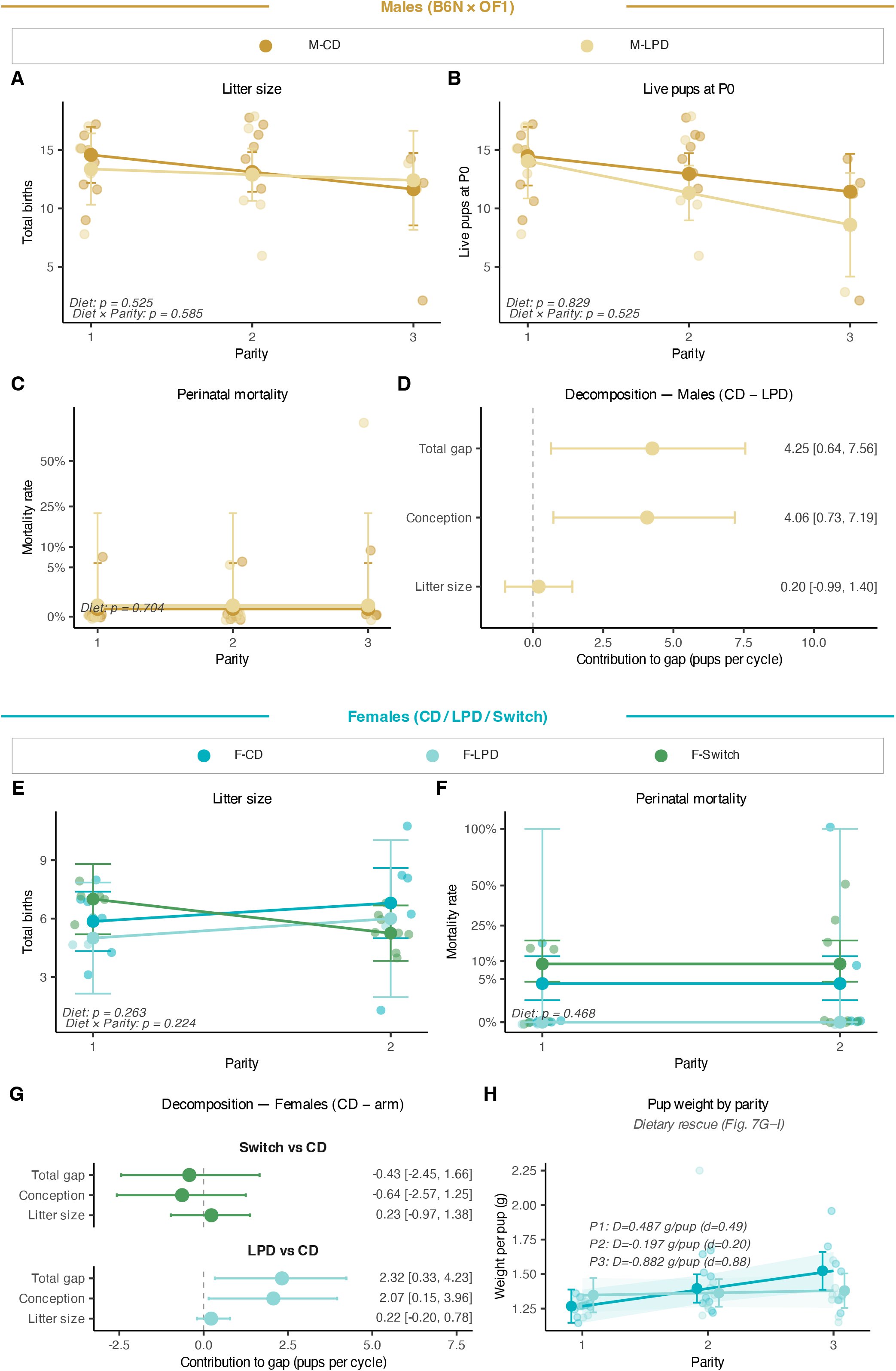

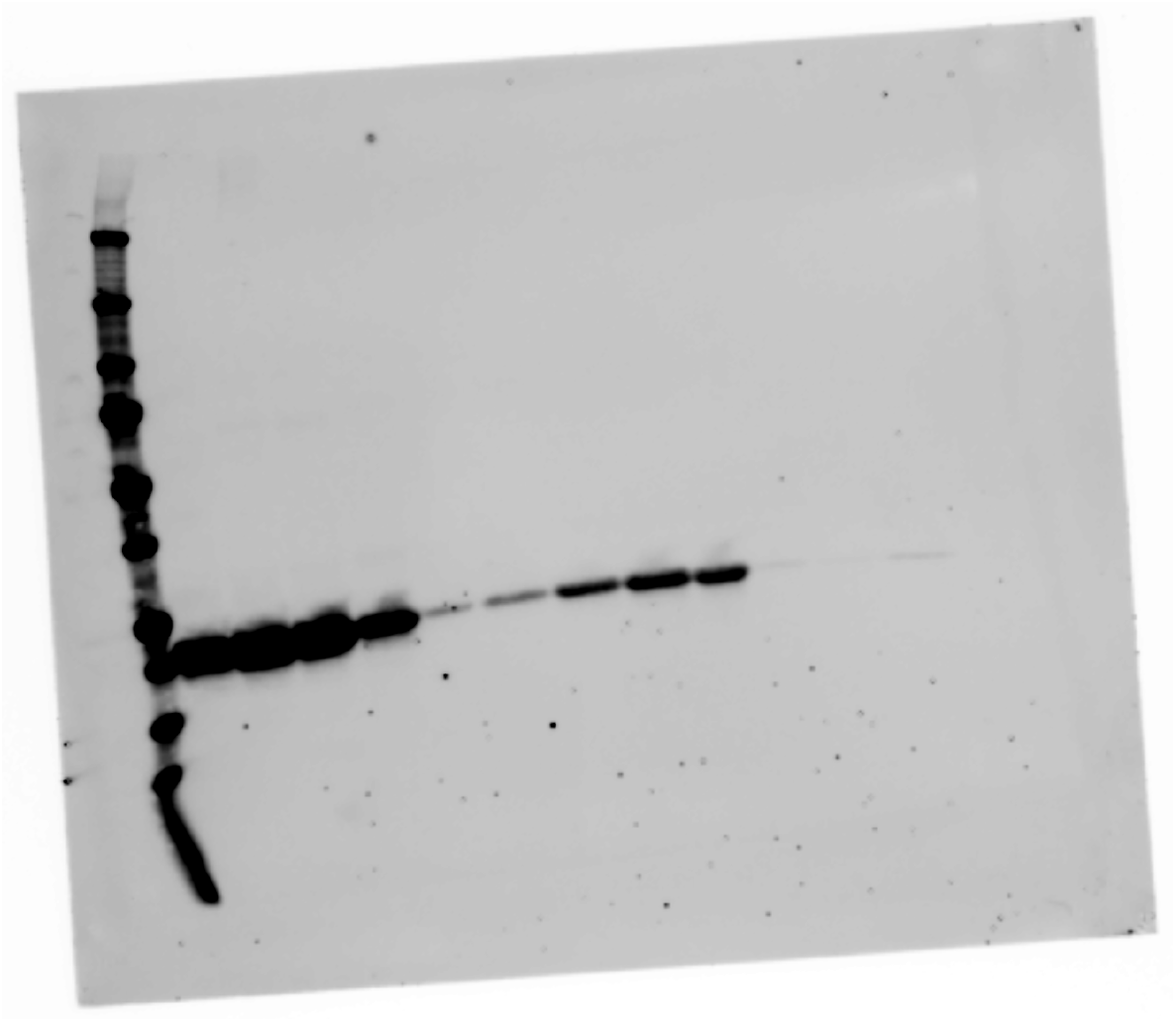

