## Supplementary Data for "Juvenile protein restriction compresses sexual dimorphism in mice"

**Short title:** Juvenile protein scarcity compresses dimorphism

**Authors:** Amélie Joly[1]*, Lucas Rebiffé[1]*, Anne Lambert[2], Estelle Caillon[3], Sandrine Hughes[3], Benjamin Gillet[3], Mickael Zergane[1], Isabelle Rahioui[4], Pedro Da Silva[4], Chantal Rigaud[5], Ingrid Plotton[5–7], Justine Bruse[8], Hervé Guillou[8], Anne Fougerat[8], François Leulier[3]†, Filipe De Vadder[3]†#

**Affiliations:**

[1]École Normale Supérieure de Lyon, Centre National de la Recherche Scientifique, Institut de Génomique Fonctionnelle de Lyon UMR5242; 69364 Lyon Cedex 07 France.

[2]Université Claude Bernard Lyon 1, École Normale Supérieure de Lyon, Centre National de la Recherche Scientifique, Institut de Génomique Fonctionnelle de Lyon UMR5242; 69364 Lyon Cedex 07 France.

[3]Centre National de la Recherche Scientifique, École Normale Supérieure de Lyon, Institut de Génomique Fonctionnelle de Lyon UMR5242; 69364 Lyon Cedex 07 France.

[4]INSA Lyon, INRAE, BF2I, UMR203; 69621 Villeurbanne, France.

[5]Hospices Civils de Lyon, Laboratoire de Biologie Médicale Multi-Sites, Laboratoire de Biologie Endocrinienne; 69500 Bron, France.

[6]Hospices Civils de Lyon, Service de médecine et biologie de la Reproduction; 69500 Bron, France.

[7]Université Claude Bernard Lyon 1, INSERM U1208, INRAE, Stem Cell and Brain Research Institute (SBRI); 69500 Bron, France

[8]Toxalim (Research Center in Food Toxicology), INRAE, ENVT, INP-PURPAN, UMR1331, Université de Toulouse; 31027 Toulouse Cedex 3, France.

* These authors contributed equally to this work

† Co-senior authors

**This PDF file includes:**

Supplementary Methods

Figs. S1 to S8

Tables S1 and S2

**Other Supplementary Materials for this manuscript include the following:**

Data S1 to S4

### Supplementary Methods

This section develops the procedures summarized in Materials and Methods. Equations are numbered S1 to S18.

#### 1. Multivariate quantification of dimorphism

*Pooled covariance.* With $G = 4$ groups (M-CD, M-LPD, F-CD, F-LPD), $n_g$ the size of group $g$ and $\mathbf{S}_g$ its sample covariance matrix, the pooled within-group covariance is

$$\Sigma_{\text{pooled}} = \frac{\sum_{g=1}^{G} (n_g - 1)\, \mathbf{S}_g}{\sum_{g=1}^{G} (n_g - 1)} \tag{S1}$$

*Mahalanobis distance.* Let $\delta_c = \bar{\mathbf{x}}_{M,c} - \bar{\mathbf{x}}_{F,c}$ denote the vector of differences between male and female group means within diet $c$, with $c \in \{\text{CD}, \text{LPD}\}$. The squared distance and its square root are

$$D_c^2 = \delta_c^{\top}\, \Sigma_{\text{pooled}}^{-1}\, \delta_c, \qquad D_c = \sqrt{D_c^2} \tag{S2}$$

The symbol $\delta$ is used here for the between-sex mean difference; $d$ is reserved throughout the manuscript for Cohen's standardized effect size.

*Compression and its test.* Compression is reported as

$$\text{Compression (\%)} = \left(1 - \frac{D_{\text{LPD}}}{D_{\text{CD}}}\right) \times 100 \tag{S3}$$

and tested by permuting sex labels within each diet independently (9,999 permutations) on the statistic $T = D_{\text{CD}}^2 - D_{\text{LPD}}^2$. Confidence intervals on the compression percentage come from a stratified bootstrap (9,999 resamples within each Sex × Diet cell). All multivariate analyses used `set.seed(42)`.

*Covariance-aware companion index.* A Bhattacharyya distance was computed in parallel and decomposed into a mean and a covariance term, $D_B = D_{\text{mean}} + D_{\text{cov}}$, with

$$D_{\text{mean}} = \tfrac{1}{8}\,\delta^{\top}\mathbf{S}^{-1}\delta, \qquad \mathbf{S} = \frac{\mathbf{S}_M + \mathbf{S}_F}{2} \tag{S4}$$

so that the part of the separation that comes from the difference in centroids can be read apart from the part that comes from the difference in dispersion. This index is reported in the deposited tables and is not shown in the figures.

*Feature sets.* The somato-metabolic level used the five traits retained by the two-step selection described in Materials and Methods; subset robustness was assessed by re-running the compression on every combination of three to five of them (median and range reported in the legend of Fig. 2J). The metabolome level used the 165-metabolite $\log_2$ matrix directly. The transcriptome level used, in parallel, the full prefiltered matrix and the CD sex-biased set, with sensitivity analyses on the top 500 and top 1,000 genes by absolute Wald statistic.

#### 2. Feminization score

The score is a scalar projection of each P56 liver sample onto a sex-discriminating axis, built in principal-component space from the 321 strictly sex-biased genes (adjusted $P < 0.05$ and $|\text{shrunken } \log_2 \text{ fold-change}| > 1$ in the CD sex contrast), a set distinct from the 1,255 genes used for the transcriptome Mahalanobis distance.

*Axis.* The discriminant direction is given in closed form by Fisher's linear discriminant,

$$\mathbf{w} = (\mathbf{S}_{\text{CD}} + \lambda\mathbf{I})^{-1}\,(\bar{\mathbf{x}}_{F\text{-CD}} - \bar{\mathbf{x}}_{M\text{-CD}}) \tag{S5}$$

where $\mathbf{S}_{\text{CD}}$ is the within-CD pooled covariance matrix in PC space, the vector $\bar{\mathbf{x}}_{F\text{-CD}} - \bar{\mathbf{x}}_{M\text{-CD}}$ points from the male-CD centroid toward the female-CD centroid, and $\lambda = 10^{-6}$ is a numerical regularizer ensuring invertibility when the sample size approaches the PC dimension. Multiplying by the inverse covariance whitens the variability within each sex, leaving an axis that carries between-sex signal only.

*Projection.* The raw score of a sample $\mathbf{x}_i$ is the inner product $s_i = \mathbf{w}^{\top}\mathbf{x}_i$, which increases from typical M-CD toward typical F-CD values. CD samples were scored by leave-one-out cross-validation, the axis being refit on the remaining CD samples for each held-out sample; LPD samples were projected on the full CD-trained axis and are therefore out of sample throughout.

*Rescaling.* Raw scores were rescaled so that the M-CD median maps to 0 and the F-CD median to 1,

$$s_i' = \frac{s_i - \text{median}\{s_j : j \in M\text{-CD}\}}{\text{median}\{s_j : j \in F\text{-CD}\} - \text{median}\{s_j : j \in M\text{-CD}\}} \tag{S6}$$

A value near 0 indicates a male-typical hepatic identity, a value near 1 a female-typical one, and values outside that interval a hyper-masculine or hyper-feminine shift. Group contrasts used aligned-rank-transform ANOVA with Holm correction. Two one-sided one-sample tests were applied to the M-LPD group against the midpoint 0.5, one for majority feminization ($H_1 : \mu > 0.5$) and one for non-inferiority to it.

##### 3. Cross-study comparison with Green *et al.*

The two datasets were generated with different library chemistries and processed with different pipelines. Those technical differences are shared by every sample of a study, so they cannot be told apart from the study itself, and expression levels cannot be compared between the two studies directly. A fold change computed inside one study escapes the problem, because the technical factor is common to its two diet arms and cancels when one is divided by the other. The comparison was therefore built from within-study diet fold changes only, as a per-gene $\log_{10}$ ratio of absolute unshrunk $\log_2$ fold-changes,

$$r_g = \log_{10} \frac{\left|\text{LFC}_g^{\text{this study}}\right|}{\left|\text{LFC}_g^{\text{Green}}\right|} \tag{S7}$$

within two functional modules: a nutrient module (union of ATF4 and PPAR$\alpha$ targets) and a dimorphic module (union of STAT5b, BCL6, GH-axis and hepatic feminization targets). A ratio of this kind can still carry a study-wide scale factor, since the two experiments differ in protein dose, exposure duration and age as well. The quantity tested is therefore not $r_g$ itself, but the difference in $r_g$ between the two modules, in which any study-wide factor cancels a second time. Module dissociation was tested by Mann-Whitney $U$ test on $r_g$, with a 10,000-permutation test on module labels as a sensitivity analysis and an ANOVA on the raw $|\text{LFC}|$ values (Module $\times$ Study $\times$ Sex) as a follow-up. A sensitivity analysis after deduplication of genes shared between sets is reported in parallel. The difference in feminization score between diets, per sex and per study,

$$\Delta s = \text{median}\left(s'_{\text{LPD}}\right) - \text{median}\left(s'_{\text{CD}}\right) \tag{S8}$$

was compared between cohorts by stratified bootstrap (9,999 resamples) and is reported as a point estimate with a 95% confidence interval.

#### 4. Curated gene sets

Each set was kept to at most 15 genes to preserve the power of the per-set enrichment test. Membership is also deposited as CSV files.

**ATF4 targets**: *Asns*, *Psat1*, *Phgdh*, *Mthfd2*, *Aldh1l2*, *Fgf21*, *Chac1*, *Atf3*, *Ddit3*, *Trib3*, *Nupr1*, *Gadd45a*, *Pck2*, *Ddit4*, *Slc7a1*. Canonical ATF4 transcripts of the amino-acid and integrated stress responses, covering the one-carbon serine and glycine branch, asparagine biosynthesis, the PERK/ATF4 effectors and *Fgf21*.

**PPAR$\alpha$ targets**: *Acot1*, *Acot2*, *Acot3*, *Acot4*, *Acot6*, *Cyp4a10*, *Cyp4a14*, *Cyp4a31*, *Ehhadh*, *Pdk4*, *Acadm*, *Fgf21*. Hepatic targets of fatty-acid $\beta$- and $\omega$-oxidation, including the *Cyp4a* subfamily, the acyl-CoA thioesterase cluster, *Acadm* and *Pdk4*. *Fgf21* is shared with the ATF4 set, reflecting its dual upstream control.

**STAT5b targets**: *Bcl6*, *Igf1*, *Socs2*, *Ghr*, *Mup1*, *Mup7*, *Mup20*, *Cyp2d9*, *Cyp4a12a*, *Ugt2b38*, *Ces1g*. Direct STAT5b targets in mouse liver, spanning the GH-pulse signaling axis, the male-biased *Mup* cluster and a representative male-biased Cyp and transferase panel.

**BCL6 targets**: *Bcl6*, *Cux2*, *Cyp2a4*, *Fmo3*, *Cyp2d9*, *Cyp4a12a*, *Cyp4a12b*, *Cyp2u1*, *Mup20*. Sex-biased hepatic effectors whose adult expression depends on BCL6 occupancy and on the dimorphic GH-pulse architecture.

**GH-axis dimorphism**: *Mup1*, *Mup7*, *Mup20*, *Cyp4a12a*, *Cyp2d9*, *Cyp2b13*, *Sult2a1*, *Cyp3a44*, *Elovl3*, *Tff3*, *Serpina1e*, *Cyp7b1*, *Slco1a4*, *Ugt2b38*, *C4a*. Liver genes whose expression follows the dimorphic GH secretion pattern.

**Hepatic feminization**: *Bcl6*, *Cux2*, *Cyp2a4*, *Fmo3*, *Igfbp1*, *Abcc12*, *Hsd17b6*, *Prlr*, *Slc22a26*, *Slc22a27*. Female-biased hepatic markers that, when induced in male liver, indicate a shift toward a female-pattern hepatic identity.

#### 5. Androgen-dependence index and pattern classification

Per-gene expression in the orchidectomy cohort was fit by a cell-means model on the four Diet $\times$ Surgery groups (`limma::lmFit` on `~ 0 + Group`), and four contrasts were extracted with empirical Bayes moderation: the diet effect within Sham, the diet effect within ORX, the surgery effect under CD, and the Diet $\times$ Surgery interaction. For each gene,

$$\rho_g = \frac{\mathrm{LFC}_g(\text{diet within ORX})}{\mathrm{LFC}_g(\text{diet within Sham})} \quad \text{(S9)}$$

A value near 1 indicates an androgen-independent diet response, near 0 a response already saturated by orchidectomy alone, between 0 and 1 a partly additive response, and below 0 a sign reversal under orchidectomy. Genes were assigned to mutually exclusive descriptive patterns from the per-contrast $q$-values and $\rho$, evaluated in this order: no diet effect ($q_{\mathrm{Sham}}$ and $q_{\mathrm{ORX}}$ both $\geq 0.05$); androgen-independent ($q_{\mathrm{Sham}} < 0.05$, $q_{\mathrm{ORX}} < 0.05$, $q_{\mathrm{int}} \geq 0.1$); saturated by orchidectomy ($q_{\mathrm{Sham}} < 0.05$, $q_{\mathrm{ORX}} \geq 0.2$, $q_{\mathrm{int}} < 0.1$, $0 \leq \rho < 0.4$); additive ($q_{\mathrm{Sham}} < 0.05$, $q_{\mathrm{ORX}} < 0.05$, $q_{\mathrm{int}} < 0.1$, $0.4 \leq \rho \leq 0.9$); sign reversal ($\rho < -0.1$ and $q_{\mathrm{Sham}} < 0.05$); partial androgen dependence ($q_{\mathrm{Sham}} < 0.05$, $q_{\mathrm{int}} < 0.1$, $0 < \rho < 0.4$). The cut-offs were fixed in advance rather than tuned. The classification annotates the continuous $\rho$ axis of Fig. S5H and no inference is drawn from the discrete classes. Per-gene values are in Data S3.

#### 6. Bayesian mediation of pubertal timing

Let $A$ denote diet ($0 = \mathrm{CD}$, $1 = \mathrm{LPD}$), $M$ the mediator (body weight at the pubertal event, interpolated between bracketing weigh days) and $Y$ the age at the event. The mediator-outcome system was fit as a multivariate Gaussian regression in `brms`,

$$M \mid A \sim \mathcal{N}\left(\alpha_0 + \alpha_A A + \alpha_C C,\ \sigma_M^2\right) \quad \text{(S10)}$$

$$Y \mid A, M \sim \mathcal{N}\left(\beta_0 + \beta_A A + \beta_M M + \beta_C C,\ \sigma_Y^2\right) \quad \text{(S11)}$$

with $C$ a per-mouse cohort term for the female analyses (Cohorts 2 and 5 pooled) and right-censored ages handled by `cens()`. Priors were weakly informative and empirically calibrated: $\mathcal{N}(\mu, 2\sigma)$ centered on the empirical marginal mean for each intercept, $\mathcal{N}(0, 2\sigma)$ for each slope, and student-t$(2, 0, \sigma)$ for each residual standard deviation. Four chains of 4,000 iterations with 1,000 warm-up gave 12,000 post-warm-up draws. Causal effects were obtained by G-computation:

for each draw, counterfactual outcomes $Y(a, M(a'))$ were simulated for $a, a' \in \{0, 1\}$ and averaged over the empirical covariate distribution, giving posteriors for

$$\mathrm{TE} = \mathbb{E}\left[Y(1) - Y(0)\right] \tag{S12}$$
$$\mathrm{NDE} = \mathbb{E}\left[Y(1, M(0)) - Y(0, M(0))\right] \tag{S13}$$
$$\mathrm{NIE} = \mathbb{E}\left[Y(0, M(1)) - Y(0, M(0))\right] \tag{S14}$$

with $\mathrm{TE} = \mathrm{NDE} + \mathrm{NIE}$ by construction. Posterior medians and 95% credible intervals are reported together with the probability of direction, $\mathrm{pd} = \max\left(\Pr(\theta > 0), \Pr(\theta < 0)\right)$. Convergence was assessed by $\hat{R} < 1.01$ on all monitored parameters, bulk and tail effective sample sizes above half the post-warm-up draws, and trace inspection. Full posterior summaries are in the Zenodo deposit.

#### 7. Effect sizes

For cross-sectional traits, Cohen's $d$ is the standardized mean difference, $d = (\bar{x}_1 - \bar{x}_2)/s_{\mathrm{pooled}}$, with

$$s_{\mathrm{pooled}} = \sqrt{\frac{(n_1 - 1)s_1^2 + (n_2 - 1)s_2^2}{n_1 + n_2 - 2}} \tag{S15}$$

For small samples ($n_1 + n_2 < 50$, as in the cross-sectional figures) the bias-corrected Hedges' $g$ is reported instead, $g = d\left(1 - \frac{3}{4(n_1 + n_2) - 9}\right)$. Approximate 95% confidence intervals come from the asymptotic variance

$$\mathrm{Var}(d) = \frac{n_1 + n_2}{n_1 n_2} + \frac{d^2}{2(n_1 + n_2)} \tag{S16}$$

as $d \pm 1.96\sqrt{\mathrm{Var}(d)}$. For contrasts derived from mixed models (longitudinal traits and tolerance tests) the standardized effect at each timepoint is the estimated marginal mean contrast divided by the residual standard deviation of the model, obtained with `emmeans::eff_size` and Satterthwaite degrees of freedom. The sign convention follows the figure legends (LPD − CD, or F − M).

#### 8. Reproductive output: two-stage decomposition

Marginal output per mating cycle was estimated as the product of the two stages,

$$\mathbb{E}\left[\text{births} \mid \text{diet}\right] = \Pr\left(\text{conception} \mid \text{diet}\right) \times \mathbb{E}\left[\text{litter size} \mid \text{conception}, \text{diet}\right] \tag{S17}$$

by non-parametric bootstrap over cycles (5,000 iterations); the LPD/CD and Switch/CD ratios are reported as bootstrap medians with percentile intervals. The gap between CD and an arm was decomposed additively (Kitagawa decomposition) into a conception-stage and a litter-size contribution,

$$E_{\text{CD}} - E_{\text{arm}} = \underbrace{s_{\text{CD}}\left(p_{\text{CD}} - p_{\text{arm}}\right)}_{\text{conception stage}} + \underbrace{p_{\text{arm}}\left(s_{\text{CD}} - s_{\text{arm}}\right)}_{\text{litter size}} \tag{S18}$$

where $p$ is the per-arm conception rate and $s$ the conditional litter size. Each contribution is reported as a bootstrap median with a percentile interval.

#### 9. Serum steroid assay

*Limits of quantification.* The analytical laboratory declared the following limits of quantification: 0.13 nmol/L for testosterone, androstenedione and 11-deoxycorticosterone, 0.40 nmol/L for corticosterone and 0.50 nmol/L for progesterone. Results below a limit were returned as numerical values rather than as censored records, so the censoring is applied here, at the declared limit, and not inherited from the laboratory report. Only testosterone is reported in the manuscript; the four other analytes are in the deposited dataset.

*Intermediate precision.* Precision was characterized during method validation on the same platform, at two internal control levels, against the minimum performance limit of the European Federation of Clinical Chemistry and Laboratory Medicine (EFLM). Every coefficient of variation lies below its limit, corticosterone at the high level being the closest to it.

| Analyte | Level | *n* | Mean (nmol/L) | SD (nmol/L) | CV (%) | EFLM limit (%) |
|---|---|---|---|---|---|---|
| Testosterone | low | 20 | 1.79 | 0.10 | 5.61 | 9.80 |
| | high | 20 | 16.61 | 1.11 | 6.71 | 9.80 |
| Δ4-androstenedione | low | 20 | 1.21 | 0.08 | 6.58 | 13.60 |
| | high | 20 | 13.22 | 0.82 | 6.23 | 13.60 |
| Progesterone | low | 20 | 6.24 | 0.48 | 7.63 | 20.00 |
| | high | 20 | 9.60 | 0.92 | 9.57 | 15.00 |
| Corticosterone | low | 18 | 2.65 | 0.21 | 7.82 | 15.00 |
| | high | 18 | 45.94 | 6.49 | 14.14 | 15.00 |
| 11-deoxycorticosterone | low | 18 | 0.61 | 0.04 | 6.30 | 15.00 |
| | high | 18 | 5.29 | 0.44 | 8.39 | 15.00 |

Both control levels lie above the testosterone concentrations measured in orchidectomized animals, so these data do not characterize precision at the bottom of that range. This is a further reason to treat values below the limit of quantification as censored rather than as measurements.

*Consequence for the testosterone model.* Thirteen of the 37 assayed samples fall at or below 0.13 nmol/L, all of them in the two orchidectomized cells (5 of 10 under CD, 8 of 8 under LPD). A cell in which nothing is quantified has no identified mean: the likelihood keeps improving as that mean is pushed towards minus infinity, and because the sum-contrast coefficients of a Diet $\times$ Surgery model are each a combination that contains it, no coefficient of that model is identified either. The diet effect on testosterone was therefore tested in sham males alone, where every value is quantified, by exact Wilcoxon rank-sum test, and detectability was compared by Fisher's exact test on the number of males above the limit of quantification. Neither the Diet $\times$ Surgery interaction nor the diet contrast within orchidectomized males is estimable, and neither is reported.

#### 10. Software

All analyses were run in R v4.4.3. The table below lists every tool and package used, with the version applied here and its citation; these are not repeated in the numbered reference list. Random number generation used a fixed seed wherever resampling or permutation was involved.

| Software or package | Version | Citation | DOI |
|---|---|---|---|
| R | 4.4.3 | R Core Team (2025) | — |
| tidyverse | 2.0.0 | Wickham *et al.*, *JOSS* 4, 1686 (2019) | 10.21105/joss.01686 |
| emmeans | 1.11.2-8 | Lenth & Piaskowski (2017) | 10.32614/CRAN.package.emmeans |
| lme4 | 1.1-37 | Bates *et al.*, *J. Stat. Softw.* 67 (2015) | 10.18637/jss.v067.i01 |
| lmerTest | 3.1-3 | Kuznetsova *et al.*, *J. Stat. Softw.* 82 (2017) | 10.18637/jss.v082.i13 |
| car | 3.1-3 | Fox & Weisberg, *An R Companion to Applied Regression*, 3rd ed. (2019) | — |

| **Software or package** | **Version** | **Citation** | **DOI** |
|---|---|---|---|
| MASS | 7.3-65 | Venables & Ripley, *Modern Applied Statistics with S*, 4th ed. (2002) | — |
| survival | 3.8-3 | Therneau (2001) | 10.32614/CRAN.package.survival |
| coxphf | 1.13.4 | Heinze *et al.* (2007) | 10.32614/CRAN.package.coxphf |
| minpack.lm | 1.2-4 | Elzhov *et al.* (2022) | 10.32614/CRAN.package.minpack.lm |
| nlstools | 2.1-0 | Baty *et al.* (2007) | 10.32614/CRAN.package.nlstools |
| segmented | 2.1-4 | Fasola *et al.*, *Comput. Stat.* 33, 997–1015 (2018) | 10.1007/s00180-017-0740-4 |
| brms | 2.23.0 | Bürkner, *J. Stat. Softw.* 100 (2021) | 10.18637/jss.v100.i05 |
| bayestestR | 0.17.0 | Makowski *et al.* (2019) | 10.32614/CRAN.package.bayestestR |
| ARTool | 0.11.2 | Kay *et al.* (2014) | 10.32614/CRAN.package.ARTool |
| vegan | 2.7-2 | Oksanen *et al.* (2001) | 10.32614/CRAN.package.vegan |
| DHARMa | 0.4.7 | Hartig (2016) | 10.32614/CRAN.package.DHARMa |
| limma | 3.62.2 | Ritchie *et al.*, *Nucleic Acids Res.* 43, e47 (2015) | 10.1093/nar/gkv007 |
| DESeq2 | 1.46.0 | Love *et al.*, *Genome Biol.* 15, 550 (2014) | 10.1186/s13059-014-0550-8 |
| apeglm | 1.28.0 | Zhu *et al.*, *Bioinformatics* 35, 2084–2092 (2019) | 10.1093/bioinformatics/bty895 |
| ashr | 2.2-63 | Stephens *et al.* (2016) | 10.32614/CRAN.package.ashr |
| fgsea | 1.32.4 | Korotkevich *et al.* (2016, preprint) | 10.1101/060012 |
| msigdbr | 25.1.1 | Dolgalev (2018) | 10.32614/CRAN.package.msigdbr |
| org.Mm.eg.db | 3.20.0 | Carlson (2024) | 10.18129/B9.bioc.org.Mm.eg.db |
| AnnotationDbi | 1.68.0 | Pagès *et al.* (2017) | 10.18129/B9.bioc.AnnotationDbi |
| pheatmap | 1.0.12 | Kolde (2010) | 10.32614/CRAN.package.pheatmap |
| Cutadapt | 3.5 | Martin, *EMBnet.journal* 17, 10–12 (2011) | 10.14806/ej.17.1.200 |
| STAR | 2.7.11a | Dobin *et al.*, *Bioinformatics* 29, 15–21 (2013) | 10.1093/bioinformatics/bts635 |
| UMI-tools | 1.1.2 | Smith *et al.*, *Genome Res.* 27, 491–499 (2017) | 10.1101/gr.209601.116 |
| HTSeq | 2.0.3 | Anders *et al.*, *Bioinformatics* 31, 166–169 (2015) | 10.1093/bioinformatics/btu638 |
| Fiji / ImageJ | — | Schindelin *et al.*, *Nat. Methods* 9, 676–682 (2012) | 10.1038/nmeth.2019 |

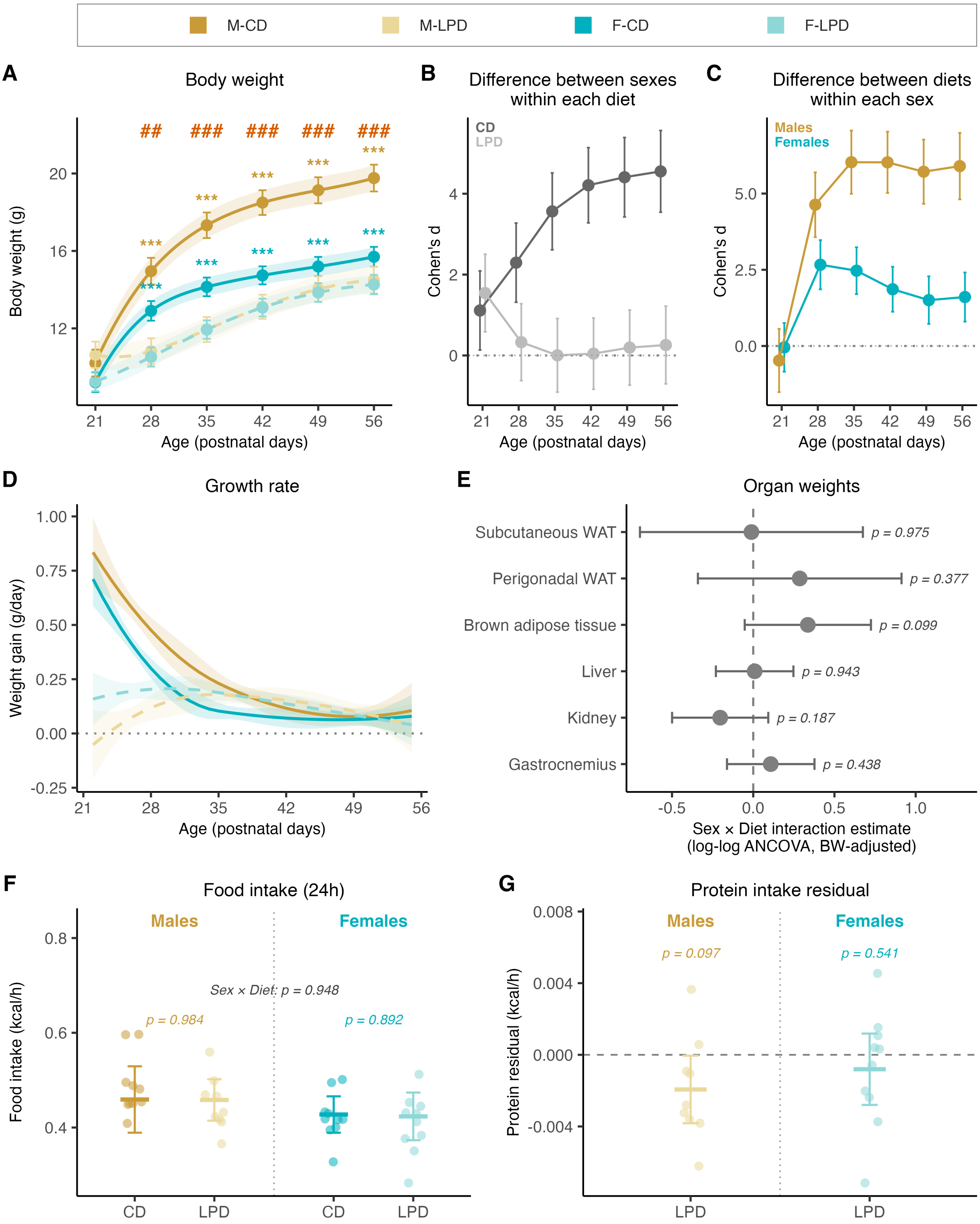


**Figure S1. Body weight, growth rate, organ allometry, and protein leverage under juvenile protein restriction.** Related to Figure 1.

C57BL/6N mice fed CD or LPD from P21 to P56. (**A**) Body weight from P21 to P56, with (**B**) the sex difference within each diet and (**C**) the diet effect within each sex, measured as Cohen's d with 95% CI. (D) Growth rate, the first derivative of the mixed-model body weight spline. (E) Sex × Diet interaction estimates from log-log ANCOVA on organ weights at P56, with body weight as covariate: subcutaneous and perigonadal white adipose tissue, brown adipose tissue, liver, kidney and gastrocnemius. Symbols, interaction coefficient; horizontal bars, 95% CI; p-value, two-sided test of the interaction term. (**F**) 24-h food intake during indirect calorimetry (P39–P41), adjusted for lean mass. (**G**) Protein intake residual, computed in LPD animals only as the difference between measured 24h protein intake and the prediction assuming no caloric compensation (CD protein intake × dilution ratio); negative values indicate compensation toward CD protein intake. Residuals were tested against zero by Wilcoxon signed-rank test (9 males, 10 females). Longitudinal traits (**A, D**) are shown as estimated marginal means with 95% CI from linear mixed-effects models with a cubic spline on age; cross-sectional measurements as individual values with mean and 95% CI. Within-sex pairwise comparisons of the diet effect are indicated by colored asterisks (yellow, males; teal, females); the Sex × Diet interaction by red hash marks (***$p < 0.001$; ##$p < 0.01$; ###$p < 0.001$). Statistical models are detailed in Methods. Sample sizes and full statistical output are in **Data S4**.

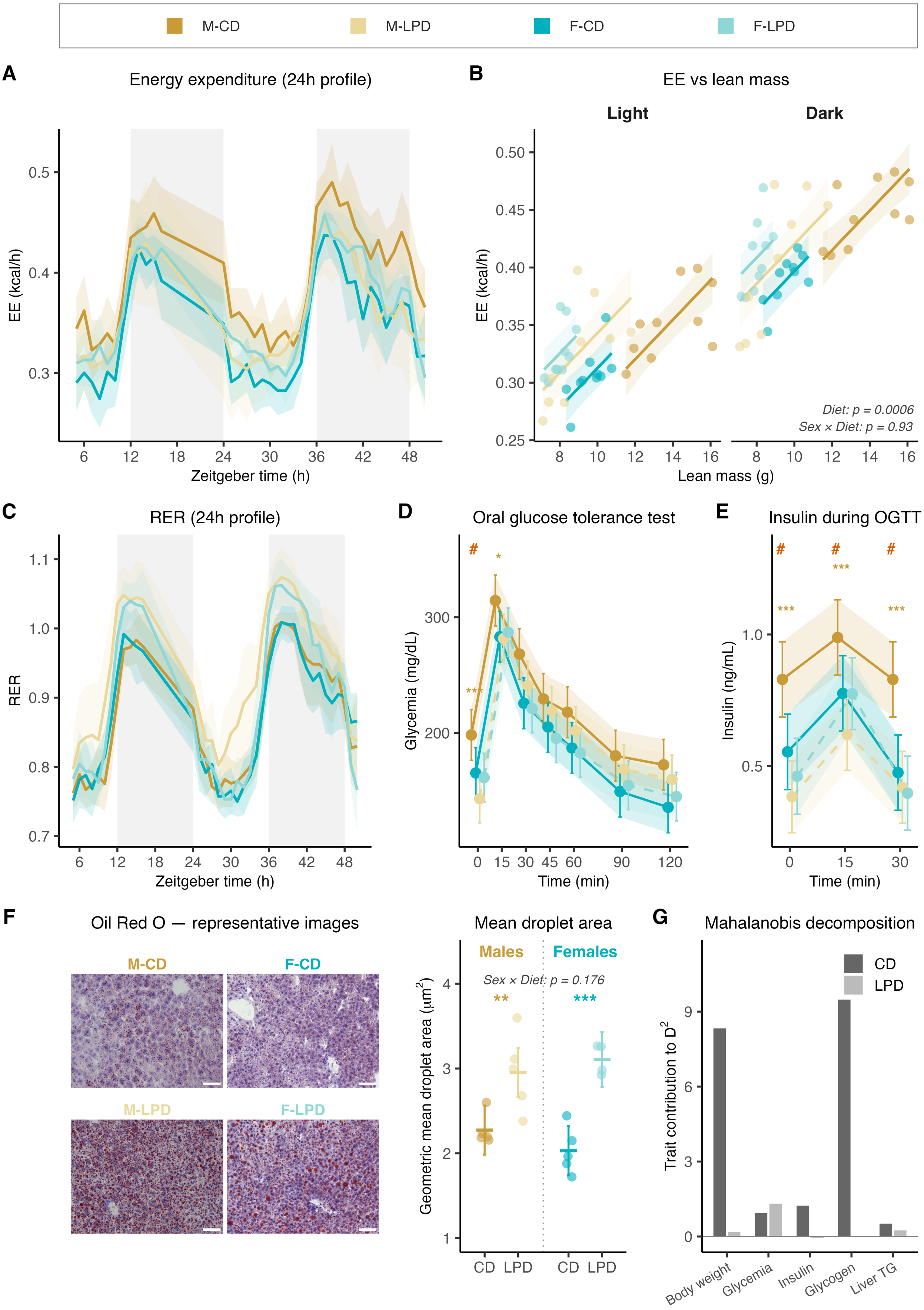

M-CD
M-LPD
F-CD
F-LPD
A
Energy expenditure (24h profile)
EE (kcal/h)
Zeitgeber time (h)
B
EE vs lean mass
Light
Dark
EE (kcal/h)
Lean mass (g)
Diet: p = 0.0006
Sex × Diet: p = 0.93
C
RER (24h profile)
RER
Zeitgeber time (h)
D
Oral glucose tolerance test
Glycemia (mg/dL)
Time (min)
E
Insulin during OGTT
Insulin (ng/mL)
Time (min)
F
Oil Red O — representative images
M-CD
F-CD
M-LPD
F-LPD
Mean droplet area
Males
Females
Sex × Diet: p = 0.176
Geometric mean droplet area (μm²)
CD
LPD
CD
LPD
G
Mahalanobis decomposition
CD
LPD
Trait contribution to D²
Body weight
Glycemia
Insulin
Glycogen
Liver TG

**Figure S2. Energy expenditure, glucose handling, hepatic lipid droplet morphology, and trait-wise decomposition of dimorphism compression.** Related to Figure 2.

C57BL/6N mice fed CD or LPD from P21. (**A**) 24-h energy expenditure (EE) profile during indirect calorimetry (P39–P41). Shaded grey bands, dark phases. (**B**) EE against lean mass in the light (left) and dark (right) phase. Lines, group-specific predicted EE from linear mixed-effects ANCOVA (`EE ~ Diet × Sex + lean mass + (1|Mouse_ID)`); shaded bands, 95% CI; p-values shown on the panel. (**C**) 24h respiratory exchange ratio (RER) profile during indirect calorimetry. (**D**) Oral glucose tolerance test at P42. (**E**) Serum insulin during the oral glucose tolerance test at P42 (t = 0, 15, 30 min). (**F**) Hepatic lipid droplet morphology at P56. Left, representative Oil Red O-stained liver sections, one mouse per group. Right, per-mouse geometric mean droplet area. (**G**) Trait-wise decomposition of the Mahalanobis $D^2$ distance between male and female centroids reported in **Fig. 2J**, into additive per-trait contributions under CD (dark grey) and LPD (light grey): body weight, fasting glycemia, fasting insulin, hepatic glycogen and liver triglycerides. Longitudinal traits (**A, C**) are shown as estimated marginal means with 95% CI from linear mixed-effects models with a 24-h-cyclic structure on Zeitgeber time. Tolerance tests (**D, E**) were analyzed by linear mixed-effects models with repeated measures on time. Mean droplet area (**F**) was analyzed by two-way ANOVA on log-transformed values. Within-sex pairwise comparisons of the diet effect are indicated by colored asterisks (yellow, males; teal, females; *p < 0.05; **p < 0.01; ***p < 0.001); the Sex × Diet interaction by red hash marks (#p < 0.05; ##p < 0.01; ###p < 0.001). Statistical models are detailed in Methods. Sample sizes and full statistical output are in **Data S4**.

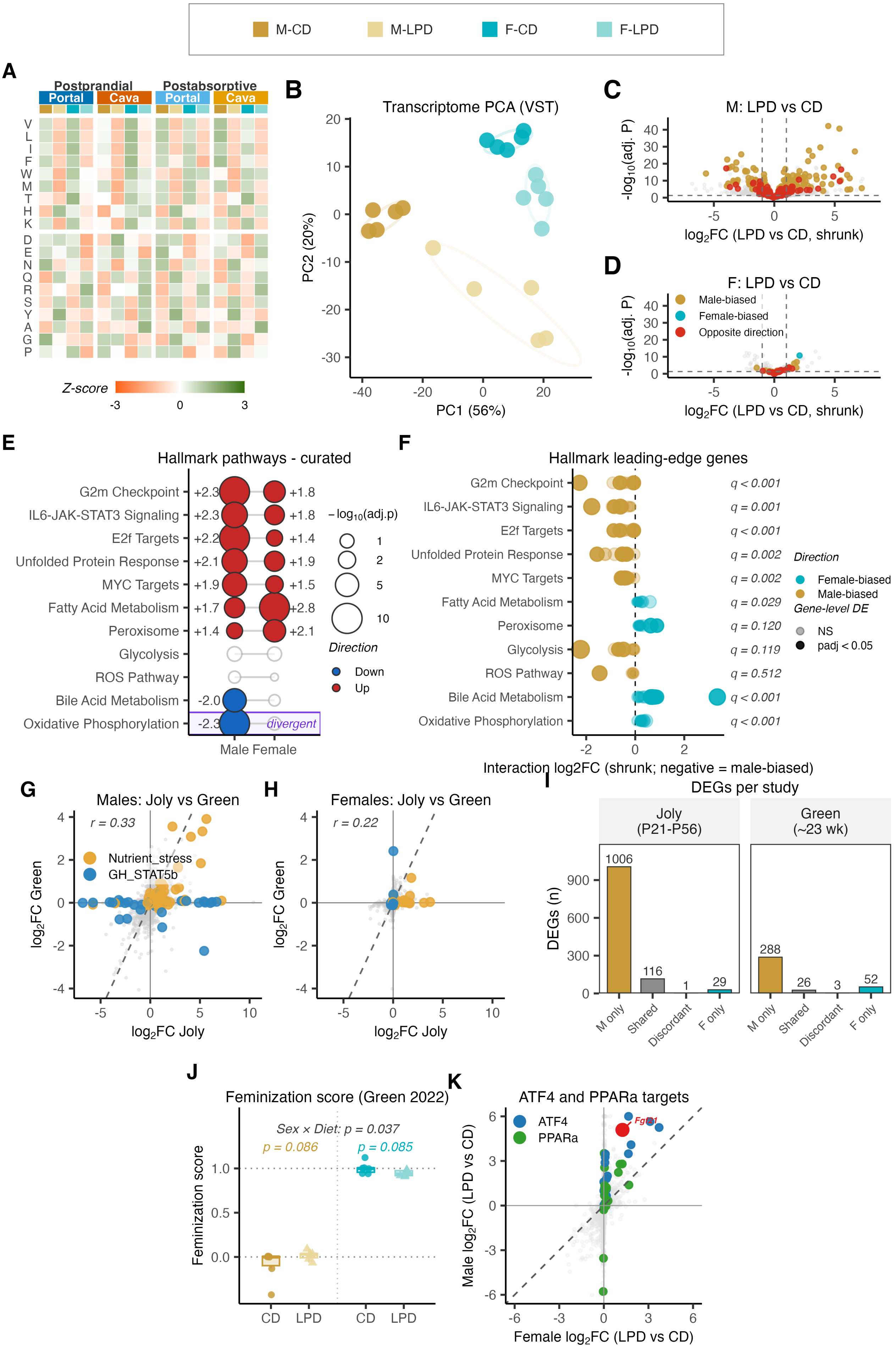

M-CD
M-LPD
F-CD
F-LPD
A
Postprandial
Postabsorptive
Portal
Cava
Portal
Cava
V L I F W M T H K D E N Q R S Y A G P
Z-score
-3
0
3
B
Transcriptome PCA (VST)
PC1 (56%)
PC2 (20%)
C
M: LPD vs CD
-log10(adj. P)
log2FC (LPD vs CD, shrunk)
D
F: LPD vs CD
Male-biased
Female-biased
Opposite direction
-log10(adj. P)
log2FC (LPD vs CD, shrunk)
E
Hallmark pathways - curated
G2m Checkpoint +2.3 +1.8
IL6-JAK-STAT3 Signaling +2.3 +1.8
E2f Targets +2.2 +1.4
Unfolded Protein Response +2.1 +1.9
MYC Targets +1.9 +1.5
Fatty Acid Metabolism +1.7 +2.8
Peroxisome +1.4 +2.1
Glycolysis
ROS Pathway
Bile Acid Metabolism -2.0
Oxidative Phosphorylation -2.3
divergent
Male Female
– log10(adj.p)
1
2
5
10
Direction
Down
Up
F
Hallmark leading-edge genes
G2m Checkpoint q < 0.001
IL6-JAK-STAT3 Signaling q = 0.001
E2f Targets q < 0.001
Unfolded Protein Response q = 0.002
MYC Targets q = 0.002
Fatty Acid Metabolism q = 0.029
Peroxisome q = 0.120
Glycolysis q = 0.119
ROS Pathway q = 0.512
Bile Acid Metabolism q < 0.001
Oxidative Phosphorylation q < 0.001
Direction
Female-biased
Male-biased
Gene-level DE
NS
padj < 0.05
Interaction log2FC (shrunk; negative = male-biased)
G
Males: Joly vs Green
r = 0.33
Nutrient_stress
GH_STAT5b
log2FC Green
log2FC Joly
H
Females: Joly vs Green
r = 0.22
log2FC Green
log2FC Joly
I
DEGs per study
Joly (P21-P56)
Green (~23 wk)
DEGs (n)
1006
116
1
29
288
26
3
52
M only
Shared
Discordant
F only
J
Feminization score (Green 2022)
Sex × Diet: p = 0.037
p = 0.086
p = 0.085
Feminization score
CD
LPD
CD
LPD
K
ATF4 and PPARa targets
ATF4
PPARa
Fgf21
Male log2FC (LPD vs CD)
Female log2FC (LPD vs CD)

**Figure S3. Hepatic transcriptome remodeling under juvenile protein restriction is male-skewed and developmentally specific.** Related to Figure 3.

Liver bulk transcriptome and circulating amino acids profiled in C57BL/6N mice fed CD or LPD from P21. (**A**) Heatmap of serum amino acids in the postprandial (PP) and postabsorptive (PA) states at P56, sampled from the portal and the cava vein. Z-score per amino acid; essential amino acids on top, non-essential at the bottom. Amino acids reported by their one-letter code. (**B**) Principal component analysis of the liver bulk transcriptome (VST-transformed) at P56 (n = 5 per group). (**C, D**) Volcano plots of differential expression LPD versus CD in males (**C**) and females (**D**), colored as in **Fig. 3E**. (**E**) Curated Hallmark gene sets with significant enrichment in either sex. (**F**) Leading-edge dotplot for Hallmark gene sets with a significant Sex × Diet interaction. (**G, H**) Per-gene LPD-versus-CD shrunken $\log_2$ fold-change, this study (x) versus Green et al. 2022 (y; GSE181301, adult C57BL/6J, re-analyzed with the same pipeline), in males (**G**) and females (**H**). Amber, nutrient-stress module (ATF4/PPARα targets); blue, GH/STAT5b module. (**I**) Diet-responsive DEG counts per study, categorized as male only, shared, discordant or female only, as in **Fig. 3D**. (**J**) Feminization score computed as in **Fig. 3H** on the Green et al. 2022 dataset; the Sex × Diet interaction and the within-sex diet contrasts are given on the panel (aligned rank transform ANOVA). (**K**) Per-gene LPD-versus-CD $\log_2$ fold-change for curated ATF4 (blue) and PPARα (green) target genes, females (x) versus males (y); *Fgf21* highlighted in red. Correlation coefficients in (**G, H**) are Pearson's r. DEGs were defined as genes with Benjamini–Hochberg-adjusted $p < 0.05$ and $|\log_2 \text{fold-change}| > 1$. Statistical models are detailed in Methods. Sample sizes and full statistical output are in **Data S4**.

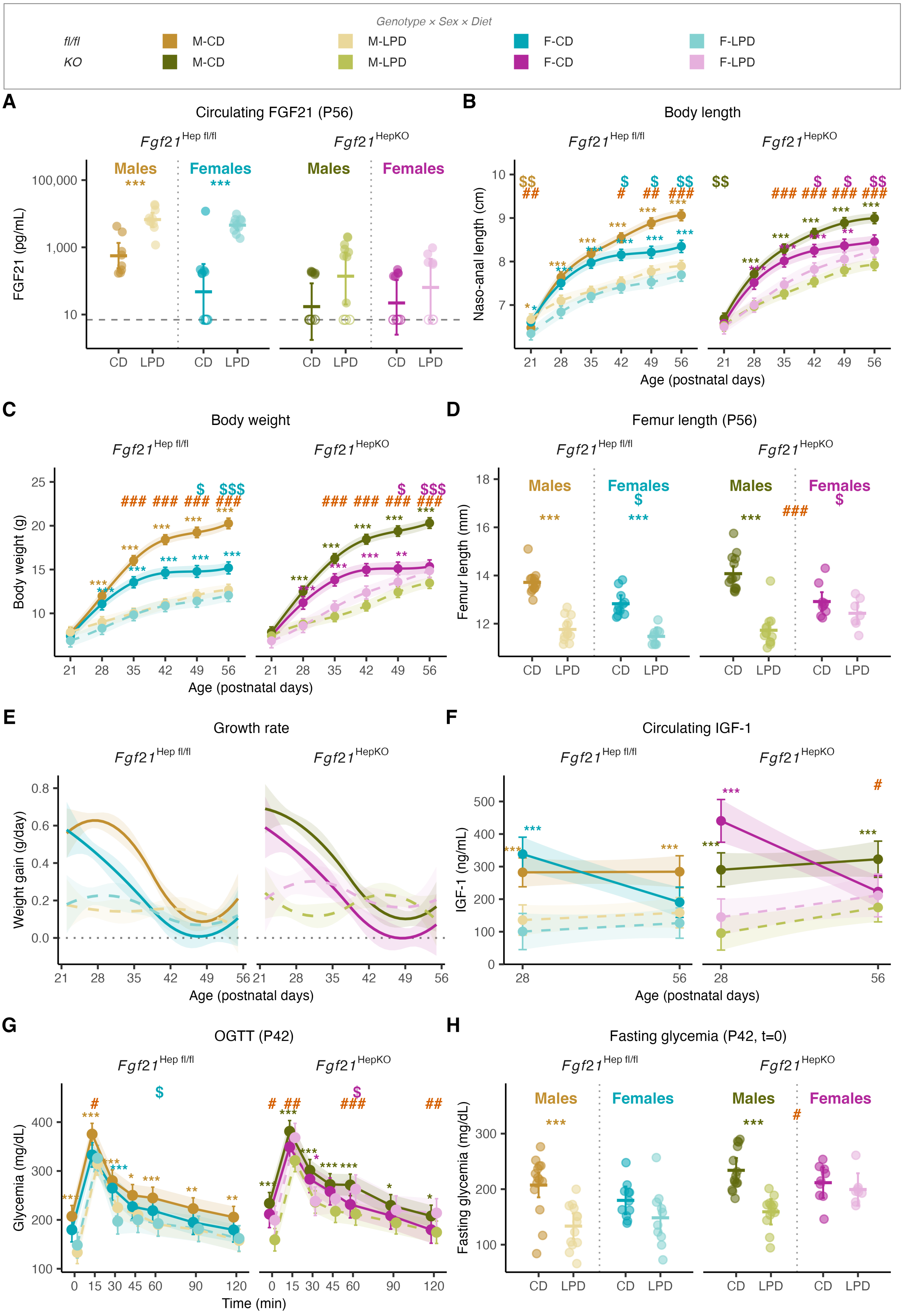
Genotype × Sex × Diet
fl/fl
KO
M-CD
M-LPD
F-CD
F-LPD
A
Circulating FGF21 (P56)
Fgf21 Hep fl/fl
Fgf21 HepKO
Males
Females
FGF21 (pg/mL)
100,000
1,000
10
CD
LPD
B
Body length
Naso-anal length (cm)
Age (postnatal days)
C
Body weight
Body weight (g)
D
Femur length (P56)
Femur length (mm)
E
Growth rate
Weight gain (g/day)
F
Circulating IGF-1
IGF-1 (ng/mL)
G
OGTT (P42)
Glycemia (mg/dL)
Time (min)
H
Fasting glycemia (P42, t=0)
Fasting glycemia (mg/dL)

**Figure S4. Hepatocyte-specific Fgf21 deletion partially rescues late female somatic growth without affecting male phenotypes under juvenile protein restriction.**

*Fgf21*$^{\text{Hepfl/fl}}$ (fl/fl) and *Fgf21*$^{\text{HepKO}}$ (KO) littermates were fed CD or LPD from P21 to P56. (**A**) Serum FGF21 concentration at P56, log scale. Dashed horizontal line, detection limit (7 pg/mL); open symbols, samples below the detection limit. (**B**) Naso-anal length from P21 to P56. Left facet, fl/fl; right facet, KO. (**C**) Body weight from P21 to P56. (**D**) Femur length at P56. (**E**) Growth rate, the first derivative of the mixed-model body weight spline. (**F**) Circulating IGF-1 at P28 and P56. (**G**) Oral glucose tolerance test at P42. (**H**) Fasting glycemia at P42 (t = 0 of the OGTT in G). FGF21 (**A**) was analyzed by Tobit regression (left-censored at the detection limit) with bootstrap 95% CI (2,000 resamplings). Longitudinal traits (**B, C, E, F, G**) were analyzed by linear mixed-effects models with a Sex × Diet × Genotype factorial design; cross-sectional measurements (**D, H**) by three-way ANOVA. Longitudinal traits are shown as estimated marginal means with 95% CI; cross-sectional measurements as individual values with mean and 95% CI. Within-sex pairwise comparisons of the diet effect are indicated by colored asterisks following the legend on top of the figure (*$p < 0.05$; **$p < 0.01$; ***$p < 0.001$). Pairwise comparisons of the diet effect within each Sex × Genotype combination are indicated by colored asterisks (one color per Sex × Genotype group, matching the legend). The Sex × Diet interaction within each genotype is indicated by red hash marks (#$p < 0.05$; ##$p < 0.01$; ###$p < 0.001$); the Diet × Genotype interaction within each sex by dollar marks ($\$p < 0.05$; $\$\$p < 0.01$; $\$\$\$p < 0.001$). Statistical models are detailed in Methods. Sample sizes and full statistical output are in **Data S4**.

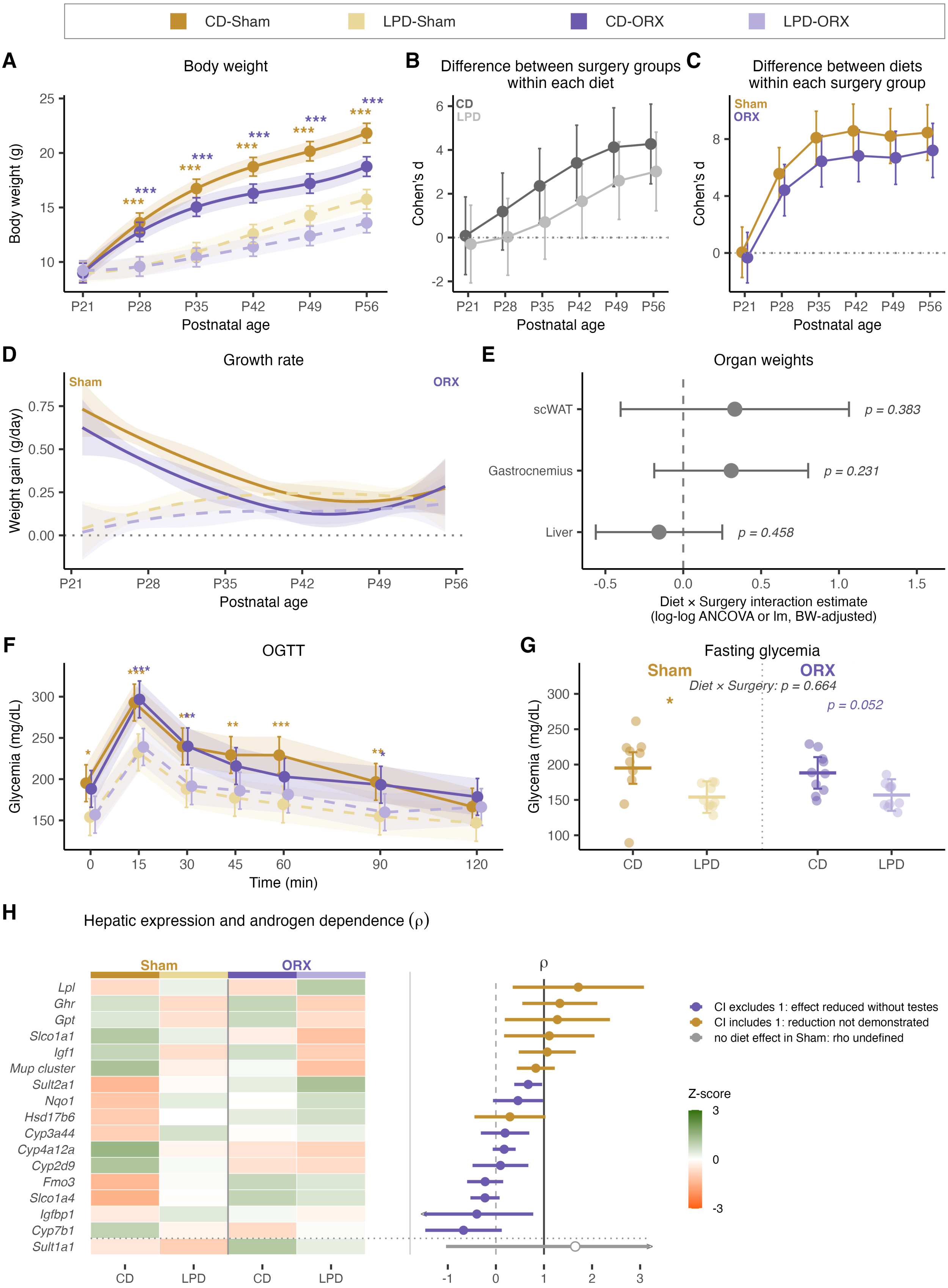


**Figure S5. Orchidectomy reproduces part of the hepatic response to juvenile protein restriction, indexed by androgen dependence.** Related to Figure 5.

C57BL/6N males underwent bilateral orchidectomy (ORX) or sham surgery at P21 and were fed CD or LPD until P56 ($n = 40$, 10 per Diet × Surgery group). (**A**) Body weight from P21 to P56, with (**B**) the surgery effect within each diet and (**C**) the diet effect within each surgery group, measured as Cohen's d with 95% CI. (**D**) Growth rate, the first derivative of the mixed-model body weight spline. (**E**) Diet × Surgery interaction estimates from ANCOVA on organ weights at P56, with body weight as covariate. Symbols, interaction coefficient; horizontal bars, 95% CI; p-value, two-sided test of the interaction term. (**F**) Oral glucose tolerance test at P42. (**G**) Fasting glycemia at P42 ($t = 0$ of the OGTT in F). (**H**) Left, heatmap of hepatic mRNA expression at P56 for 17 transcripts; columns, group means; rows, transcripts; color, Z-score per transcript. Right, per-transcript androgen-dependence index $\rho$, the ratio of the LPD $\log_2$ fold-change under ORX to the LPD $\log_2$ fold-change under Sham, with 95% CI; rows are ordered by $\rho$. The Mup cluster row corresponds to a pan-*Mup* qPCR readout (see Methods). Longitudinal traits are estimated marginal means with 95% CI; cross-sectional measurements are individual values with mean and 95% CI. Within-surgery pairwise comparisons of the diet effect are indicated by colored asterisks (yellow, Sham; purple, ORX; *$p < 0.05$; **$p < 0.01$; ***$p < 0.001$). Sample sizes and full statistical output are in **Data S4**.

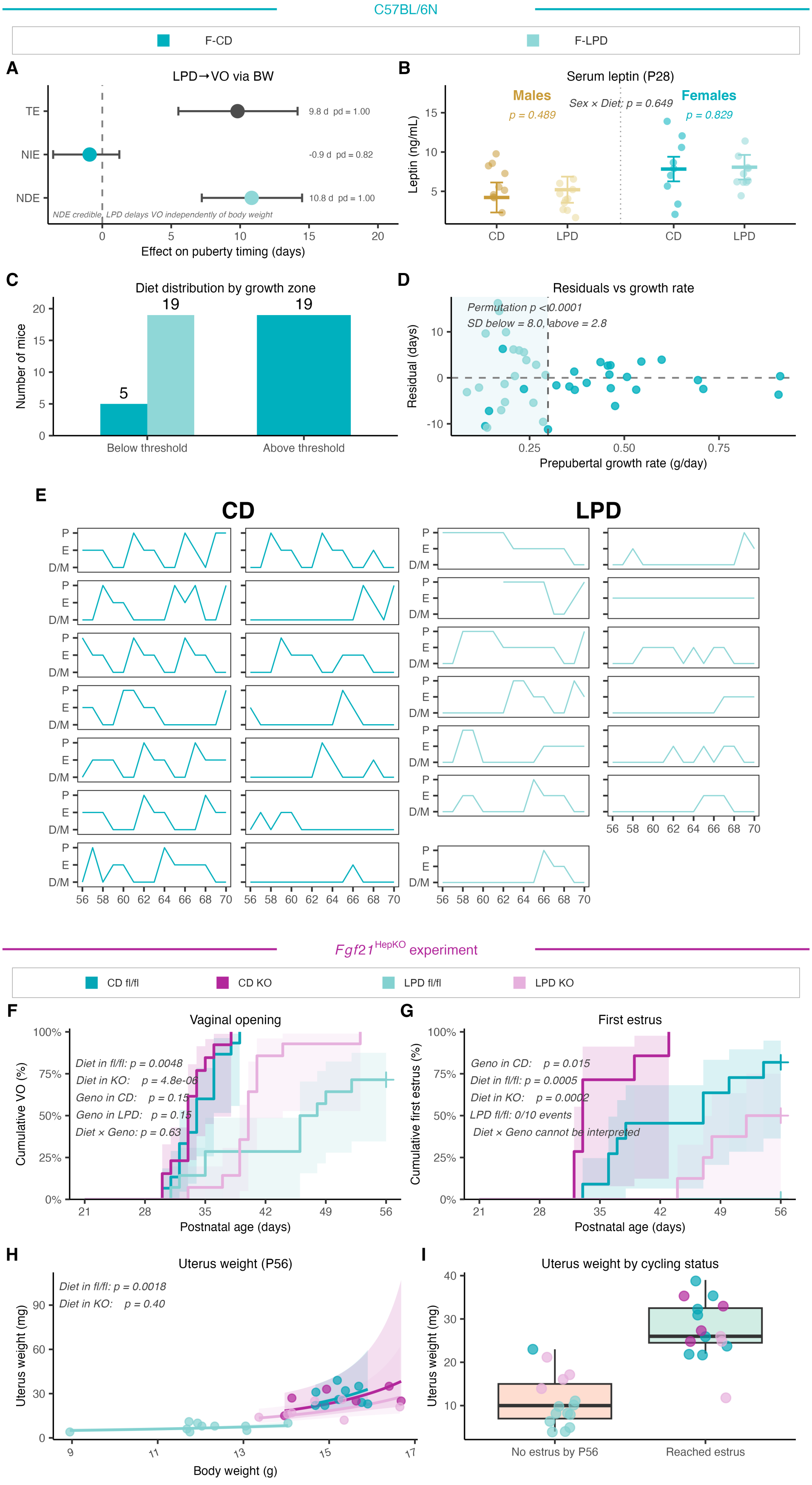

C57BL/6N
F-CD
F-LPD
A
LPD→VO via BW
TE
NIE
NDE
9.8 d pd = 1.00
-0.9 d pd = 0.82
10.8 d pd = 1.00
NDE credible, LPD delays VO independently of body weight
Effect on puberty timing (days)
B
Serum leptin (P28)
Males
p = 0.489
Sex × Diet: p = 0.649
Females
p = 0.829
Leptin (ng/mL)
CD
LPD
C
Diet distribution by growth zone
Number of mice
Below threshold
Above threshold
D
Residuals vs growth rate
Permutation p < 0.0001
SD below = 8.0, above = 2.8
Residual (days)
Prepubertal growth rate (g/day)
E
CD
LPD
P
E
D/M
Fgf21 HepKO experiment
CD fl/fl
CD KO
LPD fl/fl
LPD KO
F
Vaginal opening
Diet in fl/fl: p = 0.0048
Diet in KO: p = 4.8e-06
Geno in CD: p = 0.15
Geno in LPD: p = 0.15
Diet × Geno: p = 0.63
Cumulative VO (%)
Postnatal age (days)
G
First estrus
Geno in CD: p = 0.015
Diet in fl/fl: p = 0.0005
Diet in KO: p = 0.0002
LPD fl/fl: 0/10 events
Diet × Geno cannot be interpreted
Cumulative first estrus (%)
Postnatal age (days)
H
Uterus weight (P56)
Diet in fl/fl: p = 0.0018
Diet in KO: p = 0.40
Uterus weight (mg)
Body weight (g)
I
Uterus weight by cycling status
Uterus weight (mg)
No estrus by P56
Reached estrus

**Figure S6. Mediation of vaginal opening timing in C57BL/6N, growth-rate gating, and reproductive phenotypes in Fgf21HepKO females.** Related to Figure 6.

C57BL/6N females (**A–E**) and *Fgf21*$^{\mathrm{Hepfl/fl}}$ (fl/fl) and *Fgf21*$^{\mathrm{HepKO}}$ (KO) littermate females (**F–I**) were fed CD or LPD from P21. (**A**) Bayesian causal mediation of the LPD effect on age at vaginal opening (VO) through body weight at VO. Total effect (TE), natural direct effect (NDE; LPD → VO) and natural indirect effect (NIE; LPD → mediator → VO), as posterior median with 95% credible interval; pd, probability of direction. (**B**) Serum leptin at P28, with the within-sex diet contrasts and the Sex × Diet interaction given on the panel. (**C**) Distribution of CD and LPD females above and below the prepubertal growth-rate threshold of 0.30 g/day identified in **Fig. 6C**. (**D**) Residuals of the reciprocal model of **Fig. 6C** against prepubertal growth rate. Below the threshold the residual scatter widens (SD 8.0 days versus 2.8 days above it; permutation $p < 0.0001$). (**E**) Individual estrous cycles between P56 and P70 in all C57BL/6N females. P, proestrus; E, estrus; D/M, diestrus/metestrus. (**F**) Cumulative incidence of vaginal opening in *Fgf21*$^{\mathrm{Hepfl/fl}}$ (fl/fl) and *Fgf21*$^{\mathrm{HepKO}}$ (KO) females. Shaded areas, 95% CI of Kaplan–Meier estimates. (**G**) Cumulative incidence of first estrus in the same animals. The Diet × Genotype interaction cannot be tested because no LPD fl/fl female reached estrus by P56 (0/10 events). (**H**) Uterus weight at P56 against body weight. (**I**) Uterus weight at P56 in females that did or did not reach first estrus by P56, pooled across diet and genotype. Mediation (**A**) was performed by Bayesian models (`brms`; 4 chains × 4,000 iterations; weakly informative, empirically calibrated priors), with right-censored ages handled by `cens()`; convergence and posterior summaries are reported in Methods. Diet and Genotype contrasts for time-to-event endpoints (**F, G**) were assessed by Firth-corrected Cox proportional hazards models stratified by genotype or by diet. Uterus weight scaling (**H**) was assessed by ANCOVA on Box–Cox transformed values with body weight as covariate; the within-genotype diet p-values are shown on the panel. Sample sizes and full statistical output are in **Data S4**.

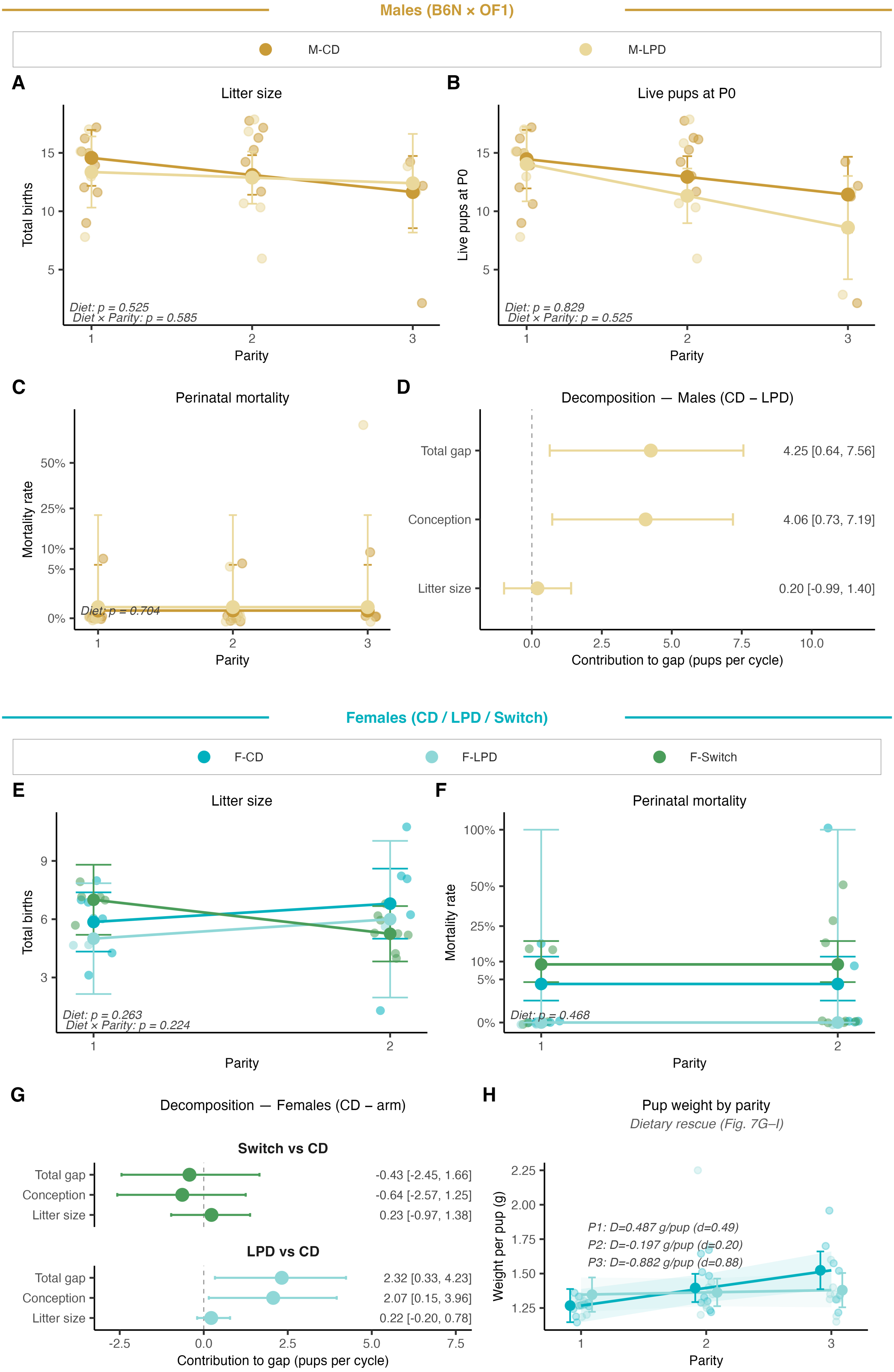
Males (B6N × OF1)
M-CD
M-LPD
A
Litter size
Total births
Parity
Diet: p = 0.525
Diet × Parity: p = 0.585
B
Live pups at P0
Live pups at P0
Parity
Diet: p = 0.829
Diet × Parity: p = 0.525
C
Perinatal mortality
Mortality rate
Parity
Diet: p = 0.704
D
Decomposition — Males (CD – LPD)
Total gap
Conception
Litter size
4.25 [0.64, 7.56]
4.06 [0.73, 7.19]
0.20 [-0.99, 1.40]
Contribution to gap (pups per cycle)
Females (CD / LPD / Switch)
F-CD
F-LPD
F-Switch
E
Litter size
Total births
Parity
Diet: p = 0.263
Diet × Parity: p = 0.224
F
Perinatal mortality
Mortality rate
Parity
Diet: p = 0.468
G
Decomposition — Females (CD – arm)
Switch vs CD
Total gap
Conception
Litter size
-0.43 [-2.45, 1.66]
-0.64 [-2.57, 1.25]
0.23 [-0.97, 1.38]
LPD vs CD
Total gap
Conception
Litter size
2.32 [0.33, 4.23]
2.07 [0.15, 3.96]
0.22 [-0.20, 0.78]
Contribution to gap (pups per cycle)
H
Pup weight by parity
Dietary rescue (Fig. 7G–I)
Weight per pup (g)
P1: D=0.487 g/pup (d=0.49)
P2: D=-0.197 g/pup (d=0.20)
P3: D=-0.882 g/pup (d=0.88)
Parity

**Figure S7. Conditional litter outcomes given conception are preserved across diets, localizing the reproductive cost to the conception stage.** Related to Figure 7.

(**A–C**) Male conditional litter outcomes (LPD versus CD), restricted to cycles with a confirmed delivery: (**A**) total births per litter, (**B**) live pups at P0, (**C**) perinatal mortality rate. Dots, individual litters; lines and bars, parity-stratified estimated marginal means with 95% CI from Diet × Parity linear models (mortality: mixed binomial GLM with a per-male random intercept, Diet only). Per-pup weight at birth for the same cohort is in **Fig. 7B**. (**D**) Kitagawa decomposition of the male per-cycle output gap (CD − LPD) into conception-stage and litter-size contributions; points, medians; bars, 95% bootstrap CI. (**E**, **F**) Female three-arm conditional outcomes given conception: (E) total births per litter; (F) perinatal mortality rate. The gestation-specific pup-weight effect that defines the three-arm cohort is shown in Fig. 7E. (**G**) Kitagawa decomposition of the female per-cycle output gap (CD − arm) into conception-stage and litter-size contributions, for LPD and Switch versus CD. (**H**) Pup weight at birth by parity in the dietary-rescue cohort (linear mixed model with a random intercept by dam; parity $\leq 3$); per-parity differences and standardized effect sizes are given on the panel. Estimated marginal means are shown with 95% CI. Sample sizes and full statistical output are in **Data S4**.

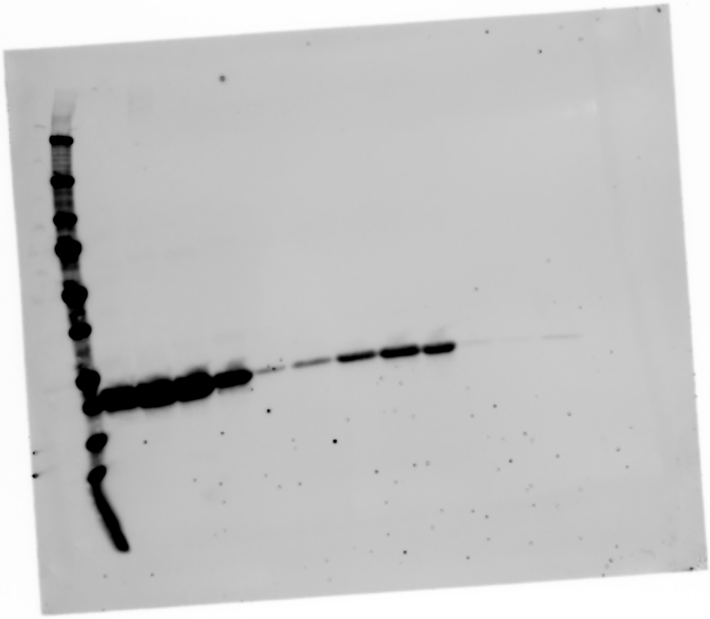

**Figure S8. Uncropped Western Blot presented in Figure 4.**

Lanes 1–3: CD males; lanes 4–6: LPD males; lanes 7–9: CD females; lanes 10–12: LPD females. Only males are shown on Figure 4.

**Table S1. Composition of the experimental diets.**

*Control (CD) and juvenile low-protein (LPD) casein-based isolipidic diets fed from P21 to P56.*

| | Control (CD) | Low-protein (LPD) |
|---|---|---|
| **Diet identity** | | |
| Supplier | SAFE (Augy, France) | SAFE (Augy, France) |
| Definition | AIN-93G-based, 20% casein, +BHT | AIN-93G-based, low-protein, +BHT |
| Catalogue number | U8978A01R 00260 | U8978A01R 00261 |
| Base | Casein-based, isolipidic | Casein-based, isolipidic |
| **Ingredient sources** | | |
| Protein source | Casein | Casein |
| Fat source | Soybean oil | Soybean oil |
| Carbohydrate sources | Cornstarch, maltodextrin, sucrose | Cornstarch, maltodextrin, sucrose |
| Fiber source | Cellulose | Cellulose |
| Mineral mix | PM AIN-93M_G (3.5%) | PM AIN-93M_G (3.5%) |
| Vitamin mix | PV AIN-93M_G (1%) | PV AIN-93M_G (1%) |
| Other | L-cystine, choline bitartrate, BHT | L-cystine, choline bitartrate, BHT |
| **Macronutrients (g/kg)** | | |
| Crude protein | 177 | 44 |
| Crude fat | 74 | 72 |
| Nitrogen-free extract (NFE) | 611 | 699 |
| of which starch | 470 | 536 |
| of which sugars | 128 | 136 |
| Crude fiber | 36 | 92 |
| Crude ash | 27 | 25 |
| Moisture | 75 | 68 |
| **Energy (% kcal, Atwater)** | | |
| Protein | 18.5 | 4.8 |
| Carbohydrate (NFE) | 64.0 | 77.2 |
| Fat | 17.5 | 18.0 |
| ME Atwater (kcal/g) | 3.82 | 3.62 |

#### Table S2. Primer sequences used for qPCR.

*Forward and reverse primer sequences and amplification efficiencies for hepatic RT-qPCR. Efficiency was determined from standard curves and is expressed as a percentage. Reference genes are highlighted in bold. Because class-B Mup paralogs share more than 97% sequence identity, the "Mup cluster" primer pair amplifies the conserved cluster and is not gene-specific; the melting curve is a single reproducible peak, which confirms a Tm-coherent amplicon family but does not establish paralog specificity.*

| Gene | Forward primer (5′→3′) | Reverse primer (5′→3′) | Efficiency (%) |
|---|---|---|---|
| ***Actb*** | **CATCCGTAAAGACCTCTATGCCAAC** | **ATGGAGCCACCGATCCACA** | **106.3** |
| ***Rpl32*** | **CCTCTGGTGAAGCCCAAGATC** | **TCTGGGTTTCCGCCAGTTT** | **98.0** |
| *Igf1* | ACCAAAATGACCGCACCTGC | AACACTCATCCACAATGCCTGTC | 103.0 |
| *Ghr* | AATCCAAGCCTGGGGACAAG | GTCTCCAGTTCAGGGGAACG | 104.6 |
| *Sult1a1* | CCAGCCCCACGGATCATTAAG | CGGGCAACGTAGATCACCTTG | 104.2 |
| *Cyp2d9* | TAATGCATTCCCGATACTCTTGCGT | TCTCAGGATTCCCTTTGGCCTTC | 99.8 |
| *Cyp3a44* | ATTCCATCTTATGCTCTTCACCATGAC | GATCAATGCTGCCCTTGTTCTCC | 100.5 |
| *Cyp4a12a* | AGTGTCCTCTAATGGCTGCAAG | GATTTGATCACTTGGTCTGTGTG | 103.3 |
| *Mup cluster* | TGTCTTGGAGAATTCCTTAG | GTTCTCGGCCATAGAGC | 100.4 |
| *Slco1a1* | GTGCATACCTAGCCAAATCACT | CCAGGCCCATAACCACACATC | 103.1 |
| *Sult2a1* | GTTCCAAGGCCAAGGCGAT | GTTCCGAGTGACCCTGGATTC | 109.3 |
| *Slco1a4* | GCTTTTCCAAGATCAAGGCATTT | CGTGGGGATACCGAATTGTCT | 103.9 |
| *Cyp7b1* | AACACCATTCCAGCTATGTTCTG | CCTCAAGAATAGTGCTTTCCAGG | 114.1 |
| *Hsd17b6* | GTTTGACCCAGGTGACTATAAGC | ACTTGGAGCAACTGTAGAATCCT | 118.0 |
| *Fmo3* | ACTGGTGGTACACAAGGCAG | ATGGTCCCATCCTCAAACACA | 105.5 |
| *Gpt* | AGCCTTTTACTGAGGTTATCCGT | TCAGAAGATTGGGGTAGACACA | 114.6 |
| *Lpl* | ATGGATGGACGGTAACGGGAA | CCCGATACAACCAGTCTACTACA | 113.7 |
| *Nqo1* | TGGCCGAACACAAGAAGCTG | GCTACGAGCACTCTCTCAAACC | 105.9 |
| *Igfbp1* | CTGCCAAACTGCAACAAGAATG | GGTCCCCTCTAGTCTCCAGA | 98.8 |
